# Multiplexed hormone immunoassays for women’s health on a regenerable silicon photonic chip

**DOI:** 10.64898/2026.09.26.754695

**Authors:** Samantha M. Grist, Maggie Wang, Lauren S. Puumala, Myra Wei, Karyn Newton, Ben Cohen-Kleinstein, Kowsar Heydaria, Sajida Chowdhury, Jaden Sequeira, Kithmin Wickremasinghe, Zavary Koehn, Nicholas Tang, Lukas Chrostowski, Sudip Shekhar, Karen C. Cheung

**Affiliations:** School of Biomedical Engineering, University of British Columbia, 251-2222 Health Sciences Mall, Vancouver, BC V6T 1Z3, Canada; Centre for Blood Research, University of British Columbia, 2350 Health Sciences Mall, Vancouver, BC V6T 1Z3, Canada; Department of Electrical and Computer Engineering, University of British Columbia, 5500-2332 Main Mall, Vancouver, BC V6T 1Z4, Canada

## Abstract

Hormone dynamics are central to a multitude of conditions in women’s health; however, limitations in the cost, convenience, and multiplexing capacity of current hormone assays preclude routine longitudinal hormone testing. This limitation severely restricts our understanding in this historically neglected field, presenting a need for improved bioanalytical tools to support hormone-informed care. Silicon photonic resonator sensors have excellent multiplexing capacity and scalable fabrication, and offer strong potential to meet this important need. Here we demonstrate, for the first time, multiplexed silicon photonic detection of two urinary hormone biomarkers, follicle-stimulating hormone (FSH) and pregnanediol-3-glucuronide (PdG), using sub-wavelength grating ring resonators and a polydopamine-based, regenerable surface chemistry. We combine a sandwich immunoassay for FSH and an antigen-down competitive immunoassay for PdG on the same chip, both amplified using a commercially available quantum-dot–streptavidin conjugate and with on-chip regeneration over multiple measurement cycles. In artificial urine, this proof-of-principle multiplexed assay achieves limits of detection of 0.594 ng/mL for FSH and 19.1 ng/mL for PdG. This architecture is inherently scalable to tens of hormone targets on a millimetre-scale chip (a >10× multiplexing improvement over typical lateral-flow immunoassays) enabling future low-cost, quantitative at-home hormone panels that could transform research, diagnostics, and treatment in women’s health.

## 1. Introduction

There is an important unmet need in women’s health for longitudinal, quantitative measurement of multiple hormones. Although it has been historically neglected, measurement of hormones relevant to women’s health has wide-ranging applications, including understanding the menstrual cycle and cycle disturbances caused by conditions like polycystic ovarian syndrome, endometriosis, and the postpartum period^1^, as well as more broadly understanding fertility, contraception^2,3^, and pregnancy loss^4^, improving personalized medicine^5,6^, and improving the standard of care for the menopausal transition^7^.

Hormone levels are dynamic, and several hormones work together to regulate the menstrual cycle and women’s health overall; as such, single or infrequent hormone measurements cannot provide the same actionable data as time-course measurements^1^. There is a current lack of information-rich longitudinal testing that means clinicians and researchers cannot capture the dynamic nature of women’s physiology^7,8^. Hormone fluctuations have historically been cited as a reason to exclude female patients and non-human mammals from drug trials^9,10^ leading to increased incidence of adverse drug reactions in female patients. Although the hypothesis that females are more variable than males due to these hormone fluctuations (a rationale previously used to justify male-only studies) has been disproven and there are growing calls for inclusion of females in biomedical research and drug testing^9,11^, there is evidence that differences in hormone levels from the menstrual cycle, life stage, pregnancy, and steroidal contraceptives could impact biomarker expression, pharmacokinetics, and pharmacodynamics^5,6^. Including hormone measurements in biomedical studies and drug trials could thus not only lead to improved understanding of the factors leading to variability, but also inform personalized medicine approaches that include hormone levels in diagnostic cutoffs, drug choice, and dosing^6,11^.

Meeting this need requires clinically accepted measurement of multiple hormones relevant to women’s health (e.g., follicle-stimulating hormone and estrogen) and hormone metabolites (e.g., pregnanediol, a progesterone metabolite) in urine^7^ or another conveniently accessible body fluid to track hormone level dynamics, in a setting and at a cost point that does not make this testing hard for patients. Today’s hormone testing platforms are subject to trade-offs in their ability to meet this need, including cost, time-to-result, multiplexing, sensitivity, and specificity. While mass spectrometry as well as the enzyme-linked immunosorbent assay (ELISA), radioimmunoassays (RIAs), and other lab-grade immunoassays are quantitative gold-standard biomarker tests, they require centralised laboratory analysis, trained technicians, long time-to-result, and expensive equipment. Mail-in hormone panel tests (e.g., the DUTCH test^12^, Everlywell^13^) similarly have long time-to-result and cost hundreds of dollars per test, preventing their application for daily monitoring. Rapid, low-cost assays like lateral-flow assays (e.g., Mira^14^, OVRY^15^, Proov^16^, Clearblue^17^ fertility hormone monitoring systems) are not able to meet the multiplexing requirements for this application (a need to measure >3 hormone markers simultaneously)^18–22^, and some are semi-quantitative and low-sensitivity. Despite improvements in quantitative lateral flow technology in recent years, there are significant barriers to high-quantity multiplexing and extended dynamic ranges for analytes of interest. For example, there are more than a dozen hormones and metabolites known to impact the menopausal transition, but multiplexing beyond 2-3 analytes on a single LFA has detrimental impacts on sensitivity and dynamic range^21^. There is thus an open need for innovation to generate the kind of dynamic hormone monitoring data required to better understand and treat the menopausal transition and women’s health more broadly.

A range of sensors for hormones and hormone metabolites in body fluids like blood, saliva, sweat, and urine have been proposed and demonstrated^23^. One promising technology to meet this open need is based on silicon photonic (SiP) integrated circuits^24^. This technology leverages semiconductor manufacturing economies of scale^25^, with excellent capacity for integration, permitting the fabrication of multiple sensors on a single chip^26^. Each sensor can have its own functionalization chemistry for multiplex detection^27^, enabling many high-sensitivity assays in a single compact device^28^ as well as references, controls, and replicates. This integration, combined with the sensors’ flexibility and information-rich monitoring of both the steady-state and dynamics of binding makes SiP sensors an attractive next-generation technology for decentralized testing.

These sensors measure changes in optical refractive index (RI) introduced by analytes binding to specific receptor chemistries immobilized on the surface of the sensor (e.g., by reading out changes in resonance wavelength of a resonator device in a fluid sample)^29^. SiP sensors are compatible with a range of different classes of bioreceptors, immobilization chemistries, and patterning techniques^30^. Although the application of SiP biosensors to decentralized testing like at-home hormone monitoring have historically been limited by the size and cost of the readout system, recent innovation in sensor architecture and laser integration could open the door for this technology to meet this need^31–33^. Figure 1(a) presents a vision for future quantitative hormone monitoring using SiP sensors.

**Figure 1.**
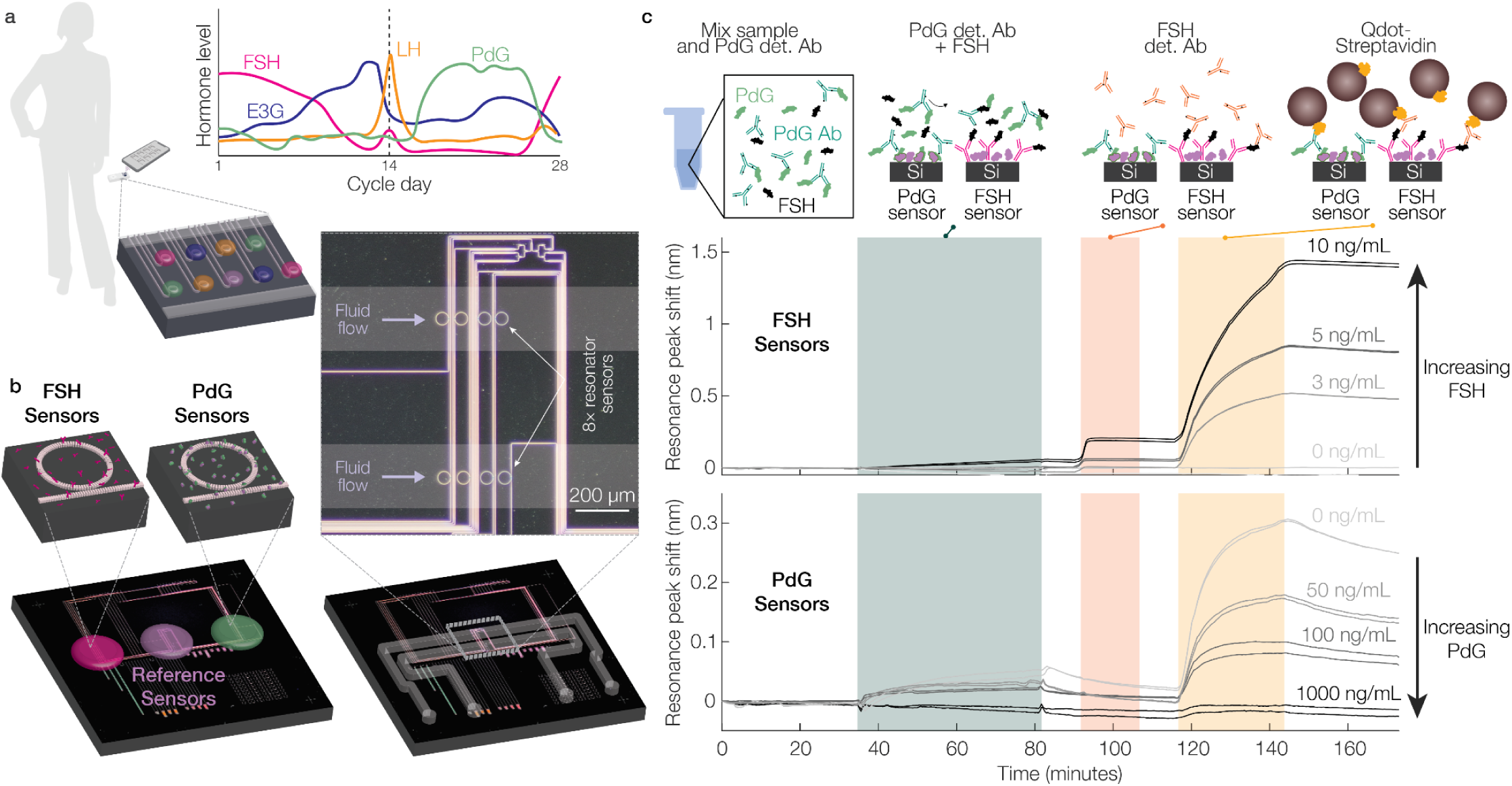
Silicon photonics-based hormone detection towards portable, low-cost urinalysis for women’s health. (a) Vision for a handheld quantification system for longitudinal monitoring of hormonal dynamics relevant to women’s health. Integration of multiplexed silicon photonic sensors will permit measurement of several hormone targets, permitting tracking of their dynamics during the menstrual cycle. Four example hormone markers (estrogen metabolite estrone-3-glucuronide, E3G; progesterone metabolite pregnanediol-3-glucuronide, PdG; luteinizing hormone, LH; and follicle-stimulating hormone, FSH) are illustrated. (b) Pilot hormone measurement system used in this work for measurement of two relevant hormone markers: follicle-stimulating hormone (FSH) and pregnanediol-3-glucuronide (PdG). Cartoons (top 3D) and micrographs (bottom and inset) depict the multiplexed SiP chip used for these experiments during functionalization spotting and integration with microfluidics. The photonic circuit and fluidic designs are described in section 8.1 of the Supplementary Information. (c) Experimental sensorgrams depicting pilot multiplexed measurements of several concentrations of FSH and PdG. FSH is detected via a sandwich immunoassay, so FSH sensor signal increases with increasing FSH concentration. PdG is detected via an antigen-down competitive binding immunoassay, so PdG sensor signal decreases with increasing PdG concentration. To increase the detected signal, both assays use amplification via CdSe nanoparticles (Qdots). Hormone dynamics illustrations depicted in (a) are adapted from Bouchard et al. 2023^1^.

SiP biosensors have been previously proposed for measurement of hormones like estradiol and progesterone^34^ without functionalization or experimental results demonstrating specific detection. Proof-of-principle detection of hormones like progesterone^35,36^, estradiol^37^, cortisol^38–40^, and testosterone^41,42^ has been reported using silicon-based sensors and similar sensors functionalized with molecularly-imprinted polymers, but experimental results demonstrating specific measurement of other hormone markers relevant to women’s health have not yet been reported. SiP biosensors are particularly attractive to meet the open need for hormone quantification due to the capability of integrated photonics technologies to deliver multiplexed hormone quantification. Our team’s previous work on SiP biosensors, which underpins the present study, has included optimizing sub-wavelength grating ring resonator biosensors^43^, understanding and improving immunoassay replicability and assay yield^44,45^, and advancing portable SiP biosensor systems for decentralized diagnostics^31,32^. There are a range of specific hormones and metabolites of interest, including estrogens and their metabolites (e.g., estrone-3-glucuronide, E3G, with a clinical range of ∼2-45 ng/mL^46^ in urine (or higher for patients undergoing gonadotropin stimulation for IVF^47^), estrone with a clinical range of ∼3-21 ng/mL^48^, estradiol with a clinical range of ∼0-11 ng/mL^48^, estrone-3-glucuronide, E1C, an estrone metabolite with clinical range of ∼500-800 pg/mL^49^), progesterone and its metabolites (e.g., pregnanediol-3-glucuronide, PdG, with a clinical range of ∼1-47 μg/mL in urine in menstruating women^46,49^), follicle-stimulating hormone (FSH, with a clinical range in urine of ∼1-70 mIU/mL^46,48^ or 0.339-24 ng/mL using the 2023 WHO standard of 1 mIU = 0.3390 ng^50^), and luteinizing hormone (LH, with a clinical range in urine of ∼1-70 mIU/mL^46,49^). Although glycoprotein hormones like FSH can be detected with a sandwich immunoassay, small hormone metabolites like PdG and E3G are typically detected using a competitive binding immunoassay. Although competitive binding assays for these targets have not yet been demonstrated on a SiP biosensor platform to the authors’ knowledge, antigen-down competitive binding immunoassays for other small-molecule targets have shown good performance, with lower limits of detection (LLoDs) down to the ng/mL or pg/mL level^51–55^.

In this work we demonstrate, for the first time, multiplexed detection of two hormone biomarkers (FSH and PdG) using sub-wavelength grating (SWG) waveguide ring resonator SiP biosensors (Figure 1). We choose these biomarkers to demonstrate the flexible power of multiplexed SiP biosensors to simultaneously detect different types of hormone markers using different types of immunoassays. FSH is a ∼35 kDa glycoprotein heterodimer^56^ often detected using a sandwich immunoassay^1^, while PdG is a small urinary steroid hormone metabolite that is often detected using a competitive immunoassay^1,57^. Here, we demonstrate a covalent sensor functionalization approach leveraging polydopamine (PDA) for bioreceptor immobilization to realize both a sandwich immunoassay to detect FSH and an antigen-down competitive binding immunoassay to detect PdG. We also leverage quantum dots for signal amplification, demonstrating an amplification factor of ∼20 while maintaining potential for sensor regeneration. We characterize the performance of both the FSH and PdG assays independently, and then assess the quantitative performance of the multiplexed assay to detect FSH and PdG in an artificial urine matrix. We then perform multiplexed FSH and PdG detection on SiP sensors functionalized via piezoelectric inkjet spotting to demonstrate the feasibility of highly multiplexed hormone detection on this platform. We anticipate that this proof-of-principle towards a quantitative, multiplexed hormone detection platform based on silicon photonic resonators will open the door to future highly multiplexed, portable devices to quantify hormone dynamics and catalyze data-driven improvements in women’s health.

## 2. Results

### 2.1 Quantum dot-mediated signal amplification yields a ∼20× increase in signal for FSH detection

For multiplexed detection of hormone targets in urine, we sought detection assays that yielded high enough sensitivity and specificity for detection of each marker at physiological levels, coupled with the ability to regenerate the sensor surface for multiple rounds of detection. Assays that employ signal amplification are commonly used for evanescent-field biosensors to increase the detected signal from small and low-concentration analytes, lowering the detection limit to meet the physiological range. Strategies for signal amplification^58^ include the use of enzymatic amplification^59–63^ (e.g., using horseradish peroxidase (HRP)-conjugated detection antibodies and flow of a precipitate-forming substrate solution like 4-chloronaphthol), the use of micro- or nanoparticles^62,64,65^, CRISPR-dCas9 (which binds its target with high specificity but does not cleave it) to amplify the signal from nucleic acid binding^66^, and high-contrast cleavage detection, which uses CRISPR-mediated cleavage of surface-bound nanoparticle reporters to detect nucleic acid targets^67–70^. While attractive due to their ability to generate large signal shifts with small, fast-diffusing enzymes, CRISPR-based strategies are limited to detection of nucleic acids and thus not well-suited to hormone marker detection. Although precipitate-forming enzymatic strategies produce very high degrees of signal amplification, we have found that amplification using 4-chloronaphthol (4CN) and 3,3′,5,5′-Tetramethylbenzidine (TMB) present challenges for regenerating the sensor surface and removing bound precipitate to prepare the sensor for subsequent binding rounds^71^. Another challenge of enzymatic amplification for applications in decentralized testing is the necessity for a separate amplification stage following the immunoassay, which increases complexity and time-to-result. In contrast, particle-based amplification strategies can be integrated into the immunoassay (e.g., using a single flow stage for antibody-particle conjugates, as in lateral flow assays). One limitation of particle-based strategies for signal amplification is the slow diffusion time of large particles^67^ due to the inverse relationship between diffusivity and particle radius via the Stokes-Einstein equation; slower diffusivity limits binding to the sensor surface^72^. There thus exists a tradeoff between diffusivity and refractive index change with particle size, because small particles are more limited in the refractive index change that they are able to impart.

For this work, we aimed to identify a regenerable amplification strategy that balanced this tradeoff. The evanescent field penetration depth in SiP biosensors is typically ∼40-200 nm, depending on polarization and waveguide geometry^73,74^; this presents an upper limit on particle size, because using particle diameters larger than the evanescent field penetration depth is expected to impart minimal benefit while reducing diffusivity. Since the functionalization chemistry and other assay components (e.g., bioreceptor, conjugation chemistry, analyte, detection antibody if a sandwich assay is used) also lie within the evanescent field, the usable detection region is thus expected to be smaller than 40-200 nm. In contrast to previous work using bead-based amplification on silicon photonic microring resonator biosensors, which used streptavidin-coated 114 nm polystyrene/iron oxide beads^65^, we aimed to evaluate the signal amplification capability of ∼15-20 nm commercially available CdSe quantum dot conjugates. CdSe has a high index of refraction (n ≈ 2.5^75,76^) in comparison to that of aqueous fluid (n ≈ 1.32^75,77^). We thus expect that bound quantum dots will result in a larger change in the effective refractive index of the propagating optical mode in the resonator (and thus more resonance peak shift signal) than lower-index particles like polystyrene or silica (SiO_2_, n ≈ 1.44^75,78^).

We tested the signal amplification capability of a commercially available quantum dot (Qdot) conjugated to streptavidin (Qdot^®^ 705 streptavidin conjugate). This product has broad visible light fluorescence excitation with suggested excitation wavelength of 400 nm, and fluorescence emission maximum near 705 nm. Although fluorescence absorption may be expected to result in optical loss and quality factor degradation in the resonator sensors if there is overlap in the relevant spectral ranges, the Qdots’ fluorescence behaviour is at wavelengths far shorter than the C-band near-infrared wavelengths used for our sensors (near 1550 nm) so this is not expected to be a dominant factor. The proprietary ∼15-20 nm Qdot nanomaterial uses a CdSe/CdSeTe semiconductor core, ZnS semiconductor shell, and polymer shell that facilitates bioconjugation of 5-10 streptavidin molecules per Qdot^79^. As shown in Figure 2, we designed a detection sandwich assay for FSH that employed a biotinylated detection antibody and biotin-streptavidin bioaffinity interaction to facilitate quantum dot-mediated amplification. We compared the signal amplification performance of the quantum dots with that of a protein-based amplification approach that our team has previously demonstrated that permits sensor regeneration^71^ (high-sensitivity streptavidin-HRP).

**Figure 2.**
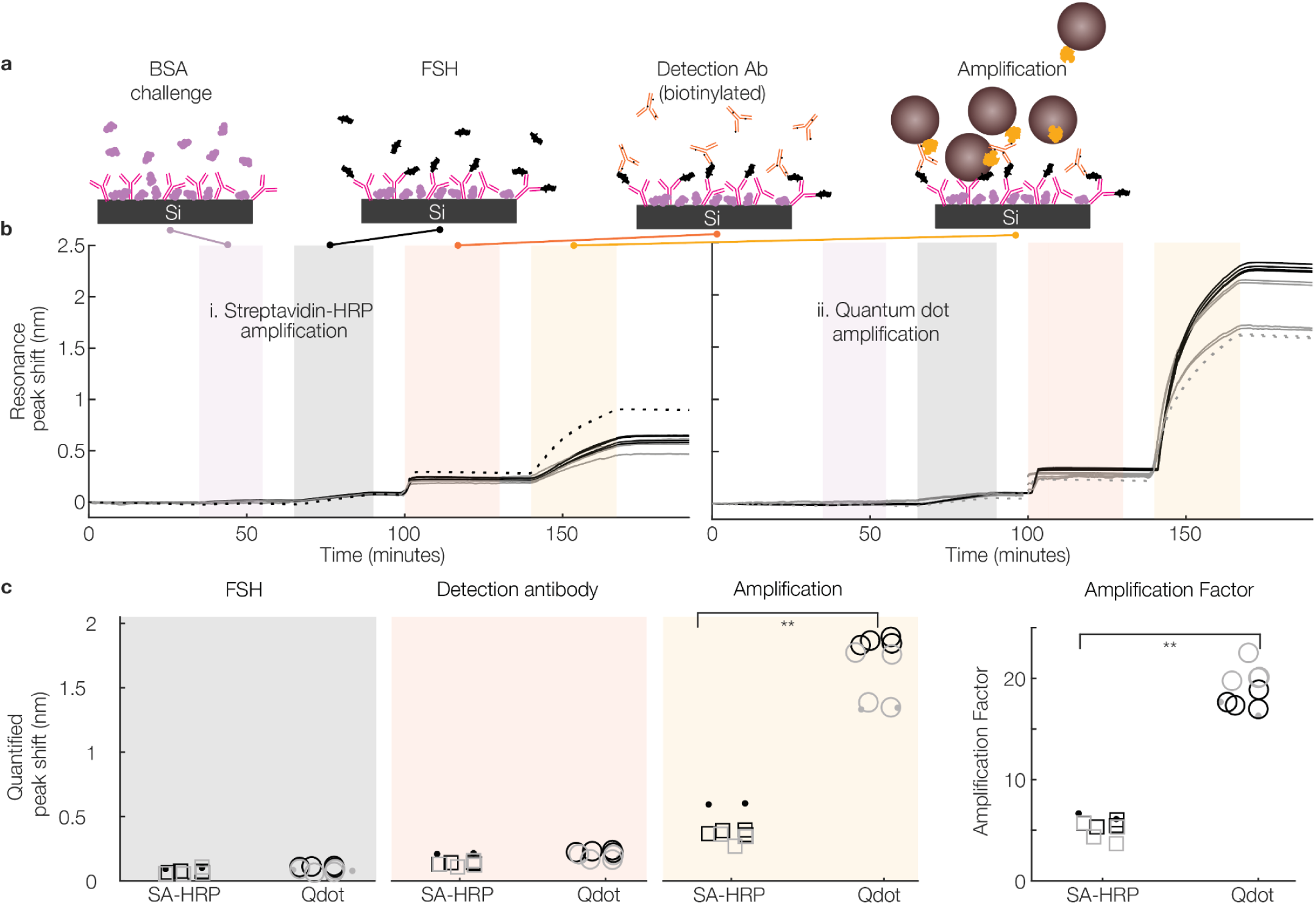
Quantum dot-mediated amplification using streptavidin-conjugated CdSe quantum dots facilitates silicon photonic detection of 25 ng/mL FSH and performs better than signal amplification using high-sensitivity streptavidin-HRP (SA-HRP). (a) Cross-sectional schematics illustrating the stages of the amplified assays, including specificity challenge, FSH binding, detection antibody binding, and amplification. (b) Example sensorgrams depicting the reference-subtracted sensor signal throughout the assay for amplification using (i) streptavidin-HRP and (ii) quantum dots (same axis scale). (c) Quantified resonance peak shift sensor signal during the FSH, detection antibody, and amplification stages for the assays using each amplification strategy, as well as computed amplification factor. Each datapoint represents the quantified signal from one sensor, and marker styles/colours denote microfluidic channel replicates (n = 3 channel replicates, each with 2-4 replicate sensors, per amplification condition). **p < 0.001, Mann-Whitney U test.

For our on-chip FSH detection assay (Figure 2(a)), we began with a SiP sensor chip functionalized with FSH capture antibodies using a straightforward polydopamine-mediated biofunctionalization process^44,80,81^, blocked with bovine serum albumin (BSA), coated with an immunoassay stabilizer, and dried prior to integration with microfluidics. We then sequentially delivered solutions of 1 mg/mL BSA to test for nonspecific binding, 25 ng/mL FSH as our test sample, 2 μg/mL biotinylated FSH detection antibody, and either 1 μM Qdot-streptavidin conjugate (Qdot) or 2 μg/mL streptavidin-HRP (SA-HRP). We included reference sensors on our SiP sensor chips which were coated with BSA during functionalization. The averaged signal from these sensors was subtracted from that of the functionalized sensors to correct for noise sources like thermal drift, nonspecific binding, and bulk refractive index changes as described in Section 8.4 of the Supplementary Information. Figure 2(b) presents sensorgrams depicting the reference-subtracted resonance peak shift vs. time during all assay steps. The amplification signal (yellow region) is markedly higher for the Qdot amplification than for the SA-HRP amplification. Quantifying these resonance peak shifts (Figure 2(c)) shows significantly higher amplification signal using Qdots (1.7 ± 0.3 nm) than using SA-HRP (0.42 ± 0.11 nm): a ∼4-fold improvement in amplification signal. Normalizing the amplification signal for each sensor by its quantified native FSH binding signal yields the amplification factor, or the degree of signal amplification provided by each approach. We measure amplification factors of 18.7 ± 1.9 for the Qdot approach and 5.4 ± 0.9 for the SA-HRP, again showing a significant improvement in the amplification performance.

The increased amplification signal from the Qdot amplification compared to the protein-only approach is expected due to the higher refractive index as well as the larger size of the nanoparticle (∼20 nm diameter; ∼4200 nm^3^ volume) compared to the protein conjugate (∼52-60 kDa for the streptavidin tetramer plus 44 kDa for horseradish peroxidase yields a minimum of ∼100 kDa for SA-HRP, which is expected to result in a minimum radius of ∼3.05 nm^82^ and a minimum spherical volume of ∼120 nm^3^). The Qdots are also expected to have higher refractive index than the proteins; refractive index of adsorbed protein layers on various surfaces has been measured at ∼1.34-1.5^83,84^, with layers of larger molecular weight proteins tending to have less dense layers with lower average indices^83^. A quantitative comparison of the expected amplification performance is challenging because the precise molecular weight, structure, and refractive index of both the commercial SA-HRP and Qdot products are proprietary; however, our observation of significantly improved amplification signal is in line with expectation.

As expected, the amplification performance of the particle-based approach is modest compared to enzymatic amplification approaches, which show amplification factors of ∼10^4^ (with the limitation of challenging regeneration using this approach^71^). Compared to previously demonstrated particle-based amplification using ∼100 nm diameter beads^65^, the amplification-stage peak shift signals that we observe are higher than the maximum observed for C-reactive protein (CRP) concentrations from 100 pg/mL to 10 μg/mL (maximum ∼0.3 nm). The amplification factor in that work was found to be highly dependent on the analyte concentration, with amplification factors decreasing with increasing analyte concentration due to amplification signal saturation near 1 ng/mL CRP. At 10 μg/mL CRP, the beads yielded ∼6-fold enhancement of the CRP binding signal; at 1 ng/mL CRP (which produced no detectable primary/native binding signal), the beads yielded 100-fold signal enhancement compared to the detection antibody signal. Another work reported ∼50-fold signal enhancement of an mRNA assay using a similar ∼114 nm diameter bead-based approach^85^. Because the primary FSH binding signal was easily resolvable in our system, we similarly expect to observe an increasing amplification factor with decreasing FSH concentration. Overall, these results demonstrate the feasibility of quantum dot-mediated amplification for silicon photonic biosensors for physiologically relevant levels of FSH detection in a sandwich assay format, and we moved to demonstrating regeneration as well as specificity of FSH detection.

Seeking to demonstrate the regeneration performance and specificity of our sandwich assay using Qdot-mediated amplification, we next ran replicate experiments comparing detection of a clinically relevant 25 ng/mL concentration of FSH in a 1:1 mixture of artificial urine in immunoassay running buffer with that of a negative control (0 ng/mL FSH in the same 1:1 mix). We demonstrated 3 binding cycles of detection, regenerating the sensor in between binding cycles using a pH 2.2 glycine-HCl regeneration solution. The results of this experiment are described in Section 1 of the Supplementary Information, and highlight specific detection of FSH in artificial urine as well as promise towards regeneration of the sensor surface for multiple immunoassay binding cycles.

### 2.2 SiP biosensors facilitate specific detection of 10 μg/mL PdG in artificial urine using an antigen-down competitive-binding immunoassay

Having demonstrated SiP-based FSH detection, we next sought to detect another hormone marker using a different class of immunoassay. We designed an antigen-down competitive binding assay to detect PdG in the same artificial urine matrix used for FSH detection in Section 1 of the Supplementary Information. Figure 3(a) presents the steps of this assay: we first functionalized the sensor surface with a PdG-BSA conjugate (using the same PDA-mediated functionalization process used for FSH capture antibody attachment), blocked the sensor with BSA, coated with an immunoassay stabilizer, and dried prior to integration with microfluidics. We then delivered a solution of 1 mg/mL BSA to test for nonspecific binding. For the PdG assay, the sample (either 0 ng/mL PdG (control) or 10 μg/mL PdG in a 1:1 mix of artificial urine and running buffer as our test sample) was mixed off-chip with 1000 ng/mL final concentration biotinylated PdG detection antibody and then delivered to the sensor. When this combined solution is delivered, the PdG detection antibody binds to the immobilized PdG antigen on the sensor surface. This detection antibody also binds to PdG in the sample (competing for binding sites with the immobilized PdG), resulting in decreased detection antibody binding to the sensor with higher PdG concentration in the sample in this competitive assay format. In the final stage of the assay, the detection antibody signal was amplified with the same Qdot solution used for the FSH assay before the sensor is regenerated with the pH 2.2 glycine-HCl regeneration solution.

**Figure 3.**
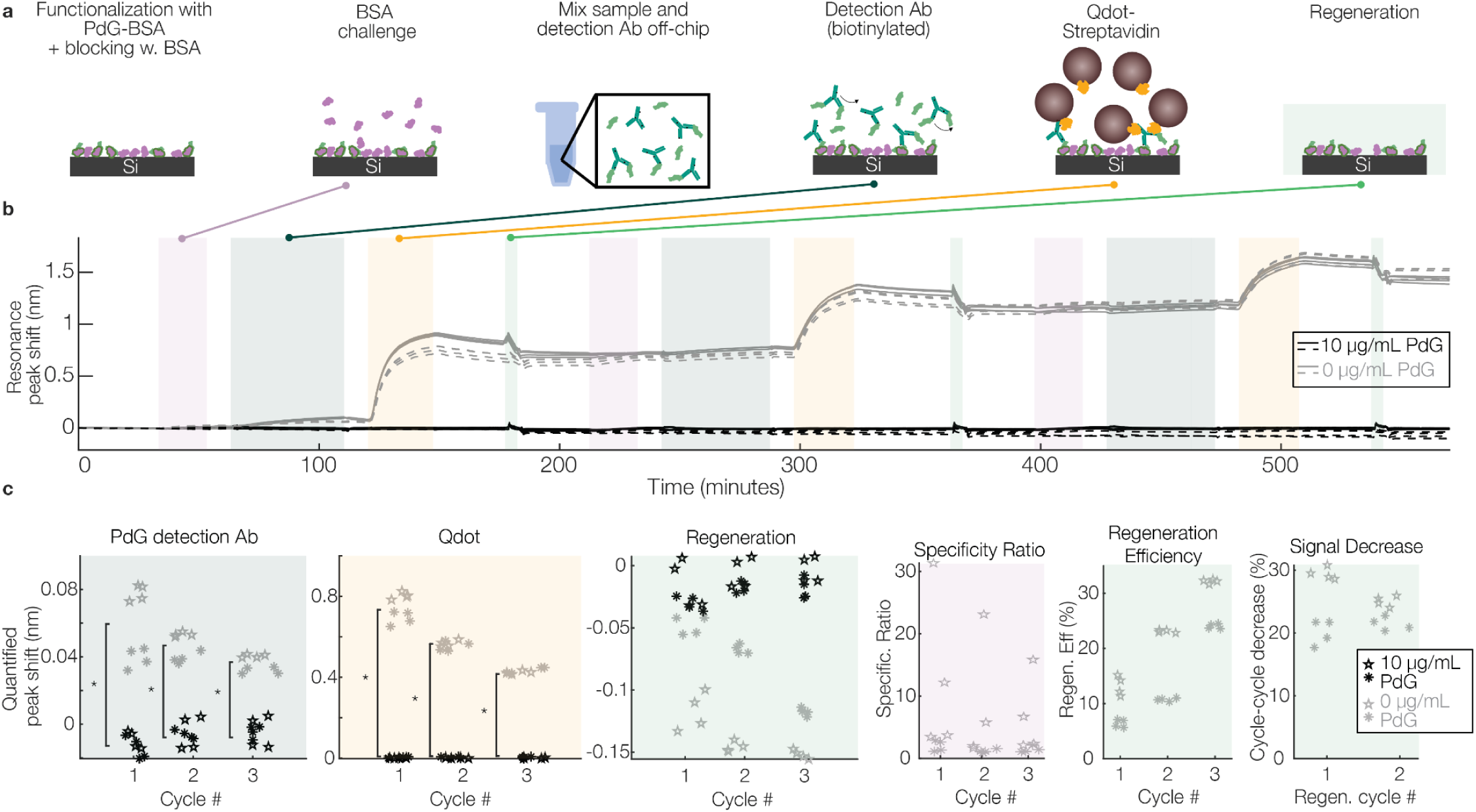
Detecting 10 μg/mL PdG in artificial urine using silicon photonic resonator sensors. (a) Cross-sectional schematics illustrating the stages of the antigen-down competitive binding assay for PdG detection, including specificity challenge, sample delivery (artificial urine control or artificial urine containing 10 μg/mL PdG), detection antibody binding, Qdot-mediated amplification, and sensor regeneration using pH 2.2 glycine-HCl. (b) sensorgrams depicting the sensor resonance peak shift signal throughout 3 cycles of the PdG detection assay. (c) quantified resonance peak shift sensor signal during the PdG sample/detection antibody, amplification, and regeneration stages for the assays, as well as computed specificity ratio (absolute value of PdG detection antibody binding signal for control channel normalized to 1 mg/mL BSA binding signal) and regeneration metrics (regeneration efficiency and cycle-cycle decrease in Qdot signal). Each datapoint represents the quantified signal from one sensor (N = 4 sensors per assay), and marker styles denote assay replicates. (*p < 0.001, Mann-Whitney U-test). Section 3 of the Supplementary Information presents the quantified intrinsic bulk refractive index sensitivity (S_bulk_) of each sensor measured in this binding assay.

Figure 3(b) presents the reference-subtracted sensorgram signals for PdG assays detecting PdG and negative control in artificial urine. As expected for this competitive assay format, there is clear signal from the PdG detection antibody binding to the sensors exposed to the negative control solution, and minimal binding signal from the sensors exposed to 10 μg/mL PdG. There are only modest regeneration shifts in all cases. Figure 3(c) presents the quantification of these reference-subtracted signals. As described in Section 1 of the Supplementary Information for FSH detection, we demonstrated specific detection in two ways: (1) by comparing the signal from PdG-containing and negative-control samples, and (2) by computing a specificity ratio, which compares the specific detection shift during the sample/PdG detection antibody binding stage (Δλ*sample*) to the nonspecific shift resulting from delivery of 1 mg/mL BSA during the nonspecific challenge stage (Δλ*BSA*). For both the primary (sample/PdG detection antibody binding) and amplification (Qdot) stages, there are significant differences between the PdG-containing and negative control channels, demonstrating specific detection of PdG in artificial urine. The CVs in the reference-subtracted binding signal for the negative control channel were 36%, 17%, and 13% for the primary PdG detection antibody binding and 8%, 4%, and 3% for the amplification stage for detection cycles 1-3, respectively (n = 8 sensors total, with n = 4 sensors in each of two duplicate assays). We observe specificity ratios of 7 ± 10, 5 ± 8, and 4 ± 5 (mean ± standard deviation of Δλ*_sample_*/Δλ*_BSA_* across all sensors) for the three binding rounds of the negative control sample, again showing higher PdG detection antibody signal than BSA binding signal.

We also evaluated regeneration performance by assessing 3 metrics of interest: the regeneration peak shift signal; the regeneration efficiency (defined as the regeneration shift divided by the sum of the shifts at the BSA, FSH, detection antibody, and Qdot assay stages, represented as a percentage: 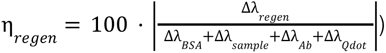, where Δλ*_regen_* represents theshift in resonance wavelength induced by the regeneration step (typically a negative (blue) shift), and Δλ*_BSA_*, Δλ*_sample_*, Δλ*_Ab_*, Δλ*_Qdot_* represent the shifts in resonance wavelength measured in response to the nonspecific BSA challenge, FSH sample, detection antibody, and quantum dot assay stages, respectively (which are all typically positive (red) shifts as material is added to the surface of the waveguide). We also evaluated the percentage by which the Qdot signal decreases between binding cycles. The regeneration efficiency gives a comparison of how much material is removed from the surface vs. how much was added during the binding assay; however, 100% regeneration efficiency does not necessarily mean that all bound material was removed because removal of the functionalization layers, blocking, and etching of the waveguides can all cause negative (blue) shifts in the resonance wavelength independent of the bound assay reagent removal. Denaturation or changes in protein conformation of the functionalization and blocking layers could also lead to changes in the thickness or refractive index of these layers, resulting in a positive or negative resonance shift. The percentage by which the Qdot signal decreases between binding cycles gives an indication of degradation in assay performance from cycle to cycle (e.g., due to functionalization denaturation or incomplete removal of bound material from the previous assay).

The negligible binding signal from the PdG-containing sample suggests that this (relatively high 10 μg/mL) concentration of PdG was above the upper limit of detection for this assay. For this kind of competitive binding assay where the detected signal is due to the balance between the detection antibody and PdG concentrations in solution^86^, increasing the range of detection will likely require increasing the PdG detection antibody concentration that is mixed with the sample, or the selection of a different detection antibody. As a starting point, showing a significant difference between 0 and 10 μg/mL PdG in this single-assay format was sufficient to move forward with development of the multiplexed assay and evaluation of the range of detection in that multiplexed assay format.

The modest regeneration shifts for the PdG assay resulted in regeneration efficiencies of only 10% ± 4%, 17% ± 7%, and 28% ± 4% for the three binding rounds of the negative control sample: considerably lower than the regeneration efficiencies for the FSH assay. The cycle-cycle decrease in Qdot signal was 25% ± 5% for the first regeneration cycle (from binding cycles 1-2) and 23% ± 2% for the second regeneration cycle. These results suggest that, like for the FSH assay, the regeneration conditions used for this assay may be too mild, incompletely removing bound material from the surface of the waveguides and further highlighting the opportunity to optimize these conditions in future work.

Overall, these results show that detection of two hormones separately is possible using silicon photonic resonator sensors. Both assays were designed and demonstrated using similar functionalization and regeneration conditions, including the use of polydopamine for sensor functionalization and use of reference sensors functionalized with BSA. This consideration affords improved likelihood of compatibility of the multiplexed assay.

### 2.3 A multiplexed SiP biosensor chip supports simultaneous detection of FSH and PdG in artificial urine

Having demonstrated the detection performance of the two individual assays, we next sought to demonstrate multiplexed detection of both FSH and PdG in the same sample. Figure 4(a) illustrates the workflow for this multiplexed assay, which is very similar to the PdG assay but with the addition of an FSH detection antibody assay stage. The sensors were first functionalized to detect FSH and PdG using FSH capture antibody and PdG-BSA as described in section 2.2 and Section 1 of the Supplementary Information, using the pipette-based spotting process illustrated in Figure 1(b). The chip was subsequently blocked, coated with immunoassay stabilizer, dried, and integrated with microfluidics. The BSA nonspecific challenge solution was the first analyte delivered to the sensors. Like in the PdG assay, the sample solution (either 25 ng/mL FSH and 10 μg/mL PdG as a hormone-containing sample or 0 ng/mL FSH and 0 μg/mL PdG as a negative control, both in a 1:1 mix of running buffer and artificial urine) was mixed with biotinylated PdG detection antibody (1000 ng/mL) off-chip and delivered through the microfluidics. Biotinylated FSH detection antibody (2 μg/mL) and Qdot solutions (5 nM) were subsequently delivered in sequence prior to regeneration with pH 2.2 glycine-HCl solution.

**Figure 4.**
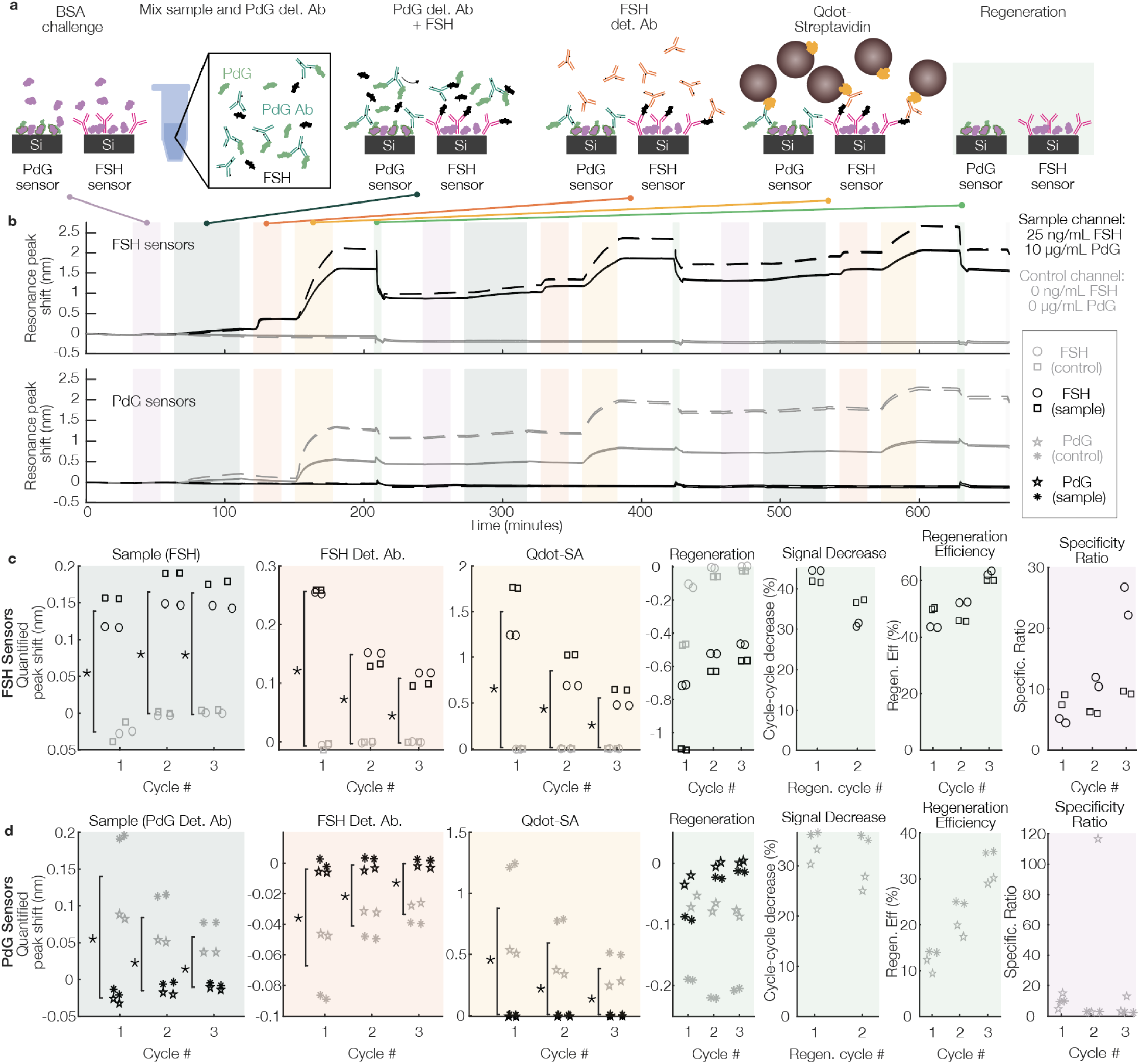
Multiplexed hormone detection using silicon photonic resonator sensors. (a) Cross-sectional schematics illustrating the stages of the multiplexed binding assay for PdG and FSH detection, including specificity challenge, sample delivery (artificial urine control or artificial urine containing 25 ng/mL FSH and 10 μg/mL PdG, mixed with PdG detection antibody), FSH detection antibody binding, Qdot-mediated amplification, and sensor regeneration using pH 2.2 glycine-HCl. (b) Reference-subtracted sensorgrams depicting the sensor signal throughout 3 cycles of the multiplexed detection assay for the FSH sensors (top) and PdG sensors (bottom). Line styles denote assay replicates. (c-d) Quantified reference-subtracted resonance peak shift sensor for the (c) FSH and (d) PdG sensors during the sample/PdG detection antibody, FSH detection antibody, amplification, and regeneration assay stages, as well as regeneration shift. Computed specificity ratio and regeneration metrics (regeneration efficiency and cycle-cycle decrease in Qdot signal) are also shown. Each datapoint represents the quantified signal from one sensor (N = 2 sensors for each target per duplicate assay for each condition), and marker styles denote assay replicates. Sensor peak shift signal for FSH- and PdG-containing samples is significantly different from that for negative control samples for the sample detection/PdG detection antibody, FSH detection antibody, and Qdot amplification assay stages for both the FSH and PdG sensors (*p < 0.05, Mann-Whitney U-test). Section 3 of the Supplementary Information presents the quantified intrinsic bulk refractive index sensitivity (S_bulk_) of each sensor measured in this binding assay.

Figure 4(b) presents the resulting reference-subtracted sensorgram signals for the FSH (top) and PdG (bottom) sensors during the 3 detection cycles. In this multiplexed assay format, we once again observe significant differences in the quantified reference-subtracted peak shift signal between the sensors exposed to hormone-containing and negative control samples during the sample, FSH detection antibody, and Qdot amplification assay stages as shown in Figure 4(c-d). The measured specificity ratios for the FSH sensors (hormone-containing sample) are 7 ± 11, 9 ± 3, and 17 ± 9, while those for the PdG sensors (negative control sample) are 10 ± 4, 31 ± 56, and 5 ± 5 for the three binding rounds. There is large variability in the specificity ratio due to the near-zero and slightly negative nonspecific binding signals in some cases.

We observe higher signal variability across these duplicate trials than in the single-analyte assays, with one trial yielding much higher signal than the previous single-analyte trials in both the sample and Qdot binding stages. Curiously, although the FSH detection antibody binding signal from the FSH sensors shows lower variability between the two replicates, the first-cycle average binding signal (0.256 ± 0.003 nm) during the detection antibody stage for both trials was considerably higher than that previously measured during the FSH-only assays (0.14 ± 0.02 in Section 1 of the Supplementary Information).

Nevertheless, these results demonstrate specific multiplexed detection of both PdG and FSH in artificial urine, with detection performance similar to that of the single-hormone assays. Notably, the signal from the FSH sensors exposed to the negative control sample as well as that from the PdG sensors exposed to FSH remains near zero, suggesting minimal cross-reactivity of the FSH sensors with the PdG detection antibody present in the negative control sample, or of the PdG sensors with the FSH present in the hormone-containing sample.

It is interesting to note that there are significant differences in the quantified signal from the sensors exposed to hormone-containing sample vs. the negative control during the FSH detection antibody binding stage for both the FSH and PdG sensors, despite the fact that there should be no expected binding interaction between the FSH detection antibody and PdG sensors. For the PdG sensors, we hypothesize that the signal measured here (more negative for the negative control channel that showed binding signal, and close to zero for the hormone-containing channel) results from the natural off-rate of the PdG detection antibody as it dissociates from the PdG-BSA on the sensor surface during the 20-min FSH detection antibody binding stage. This dissociation would be expected to reduce the amplified binding signal due to the reduction in surface-bound PdG detection antibody remaining during the amplification assay stage, and it could be reduced by selecting a different PdG antibody with lower off-rate, or by reducing the length of the FSH detection antibody binding stage (which is possible in this case due to the fast kinetics of the observed FSH detection antibody binding, with signal saturating well before the end of the assay stage). It may also be possible to leverage this signal for detection.

Regeneration performance was similar for the multiplexed assay format compared to the single-hormone assays, with regeneration efficiencies for the three detection cycles of 47% ± 4, 49% ± 4%, and 61.5% ± 1.6% for the FSH sensors (hormone-containing sample), and 12% ± 2%, 22% ± 4%, and 33% ± 4% for the PdG sensors (negative control). The cycle-cycle decrease in Qdot binding signal after regeneration rounds 1-2 was 43.1% ± 1.6% and 34% ± 3% for the FSH sensors (hormone-containing sample), and 34% ± 3% and 31% ± 5% for the PdG sensors (negative-control sample). These pilot measurement results demonstrate proof-of-principle for multiplexed SiP hormone detection from the same sample, and we next sought to assess the system’s quantitative performance and measurement range.

### 2.4 Multiplexed SiP assay shows promise for on-chip hormone quantification

To evaluate the quantitative performance of our multiplexed assay, sensors were exposed to a binding assay with hormone samples in artificial urine with concentrations from 0-50 ng/mL FSH and 0-20 μg/mL PdG. Figure 5 presents the results of this evaluation. For these measurements (Figure 5(a)), we used the same multiplexed assay described in Figure 4 but with only a single binding cycle and no nonspecific challenge stage, to mitigate regeneration-related variability effects (as regeneration procedures will be refined in future studies) and better recapitulate the workflow for a typical single-use test. Figure 5(b) presents representative reference-subtracted sensorgrams illustrating the concentration-dependent signal for each analyte, while Figure 5(c-d) presents (i) the quantified reference-subtracted resonance peak shift vs. concentration for the sample, FSH detection antibody, and quantum dot amplification assay stages, (ii) the results of a quantum dot-stage calibration analysis using a 4-parameter logistic regression, (iii) the quantified CV of the detection signal at each assay stage, and (iv) the relative uncertainty of quantification (width of a 95% confidence interval on quantified hormone concentrations normalized to the quantified concentration) for calibration fits for each stage of the assay. For both assays, we observe concentration-dependent signal that fits well to a 4-parameter logistic function. We also performed calibration curve fitting for the FSH detection antibody and primary sample stages of the binding assay and compared the use of a 5-parameter logistic calibration function, and these results are presented in section 4 of the Supplementary Information. Building upon the FSH amplification assessment reported in Figure 2, section 5 of the Supplementary Information reports the concentration-dependent amplification factors observed for FSH detection in our multiplexed assay. Similar to the relationship reported previously for another particle-mediated amplification approach^65^, we observe increasing amplification factor for decreasing FSH concentration, with Qdot amplification factors of >100 for FSH concentrations ≤ 3 ng/mL.

**Figure 5.**
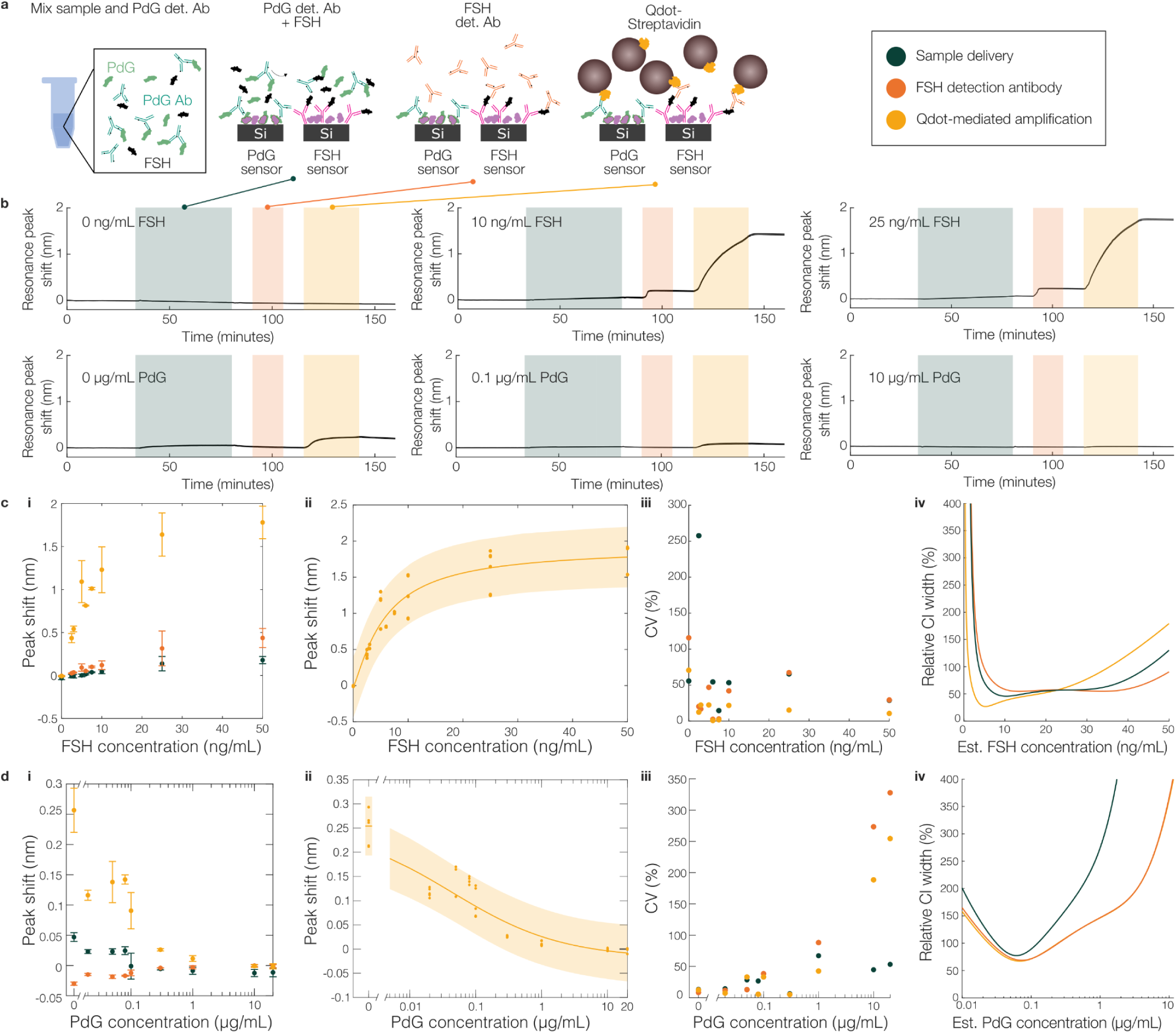
Quantification of multiplexed hormone detection using silicon photonic resonator sensors. (a) Cross-sectional schematics illustrating the stages of the multiplexed binding assay for PdG and FSH detection, including sample delivery, FSH detection antibody binding, and Qdot-mediated amplification. (b) Reference-subtracted sensorgrams depicting the FSH (top) and PdG (bottom) sensor signal for three representative sample concentrations detected using the multiplexed detection assay (signal from n = 2 sensors/plot). (c-d) Quantification performance of SiP sensors for both (c) FSH and (d) PdG detection. (i) Average reference-subtracted peak shift of each assay stage (sample delivery, FSH detection binding, Qdot-mediated amplification) at each calibration concentration). Error bars represent one standard deviation of n = 4, 6, or 8 sensors) (ii) Reference-subtracted peak shift values (datapoints), fitted calibration curves (lines) and 95% confidence intervals (shaded overlay) for the Qdot-mediated amplification stage. FSH curve fit parameters: a = -0.01734 ± 0.16034 nm; b = 1.214 ± 0.4727; c = 5.82 ± 1.972 ng/mL; d = 1.91 ± 0.287 nm; R^2^ = 0.8994. PdG curve fit parameters: a = 0.2542 ± 0.023 nm; b = 0.5177 ± 0.2357; c = 0.04346 ± 0.02647 μg/mL; d = -0.01909 ± 0.03004 nm; R^2^ = 0.9027. The FSH data (cii) are plotted on a linear x-axis scale, while the PdG data (dii) are plotted on a logarithmic x-axis from 0.01 to 10 µg/mL and a linear x-axis at a concentration of 0 µg/mL. (iii) Coefficient of variation (CV) of binding shifts of each assay stage at each calibration concentration, with PdG plotted using the same broken x-axis as previously described. (iv) Relative confidence interval (CI) widths of fits performed on the binding data from each assay stage, calculated as the upper bound of the estimated concentration minus the lower bound, as a percentage of the estimated concentration. More details on the relative CI widths can be found in section 9 of the Supplementary Information. Section 3 of the Supplementary Information presents the quantified intrinsic bulk refractive index sensitivity (S_bulk_) of each sensor measured in this binding assay.

Using the calibration functions, we can assess the limits of detection as previously described^87^. The limit of blank (LoB) was calculated from the calibration functions and zero-concentration measurements as LoB = μ_Blank_ + 1.645 ᐧ σ_Blank_, where μ_Blank_ and σ_Blank_ represent the mean and standard deviation, respectively, of a set of measurements of a blank (zero-concentration sample). The LoB for the FSH assay was calculated as 0.129 ng/mL, while that for the PdG assay was calculated as 1.75 ng/mL. The limit of detection (LoD) was calculated as LoD = LoB + 1.645 ᐧ σ_Low-concentration_, where σ_Low-concentration_ represents the standard deviation of a set of measurements of a low-concentration sample. The LoD of the FSH assay was calculated as 0.594 ng/mL, while that for the PdG assay was calculated as 19.1 ng/mL. The FSH LoD may be overestimated here because the lowest tested FSH concentration was 2.5 ng/mL (with standard deviation 0.263 ng/mL). Lower concentrations (approaching the LoD) would likely have lower standard deviation. In contrast, the LoD and LoB for the PdG assay are likely underestimated because only 2 of 8 measurements of 0 ng/mL PdG were quantifiable and used in the LoB and LoD calculations, due to the inter-assay variability (the remaining 6 measurements had Qdot amplification shift signals above the calibration fit value for ‘a’ (0.2542 nm), resulting in a non-quantifiable (imaginary) inverse 4-parameter logistic value).

Quantification is limited by the inter-assay variability in binding signal, which results in relatively high quantification uncertainty (with the relative width of a 95% confidence interval on quantified data having a minimum of 26.82% at 5.3 ng/mL for the FSH assay, and 68.99% at 68 ng/mL for the PdG assay). The relative uncertainty for the FSH assay remains below 50% for FSH concentrations from 2.6 to 18.3 ng/mL and remains below 100% for FSH concentrations from 1.3 to 35.5 ng/mL, while that for the PdG assay remains below 100% for PdG concentrations from 25 to 233 ng/mL. The presence of the second analyte does not appear to have an effect on detected signal and quantified PdG or FSH concentration, as shown in section 6 of the Supplementary Information. This variability likely stems from the immunoassay rather than the intrinsic sensor sensitivity, because correcting the quantum dot amplification-stage signal by the intrinsic S_bulk_ for each sensor does not significantly improve the average CV, as shown in section 7 of the Supplementary Information.

For the multiplexed quantification assays presented in Figure 5, we used a combination of protein and polymer blocking, as this was previously found to yield improved blocking of interferometric sensors for tuberculosis detection in urine specimens compared to either protein-based or polymer-based blocking alone^88^. Although the FSH assay signal remained similar, we observed poorer detection signal for the PdG assay in Figure 5 compared to protein blocking alone (Figures 3-4). We hypothesize that this may be due to the smaller size of the bioreceptors used for the antigen-down competitive binding assay (∼66 kDa BSA-PdG compared to ∼150 kDa IgG in the sandwich assay), which may lead to additional steric hindrance effects from the surface-attached PEG and effective reduction in the concentration of surface binding sites. Future studies could assess PLL-g-PEG formulations with lower PEG molecular weight than the 5 kDa product used here, which may offer an additional means to mitigate this challenge.

The range tested here aligns well with the clinical range for FSH in urine (∼1-70 mIU/mL^46,48^ or 0.339-24 ng/mL using the 2023 WHO standard of 1 mIU = 0.3390 ng^50^). The quantifiable range for PdG, in contrast, is lower than the clinical range for PdG in urine (∼1-47 μg/mL in menstruating women^46,49^). Diluting the sample, increasing the PdG detection antibody concentration, or selecting an alternate antibody could each be strategies to increase the quantifiable range for PdG to higher concentrations to align with the clinical range in future work. Improving the inter-assay replicability will be a critical step to improve the quantitative performance of this multiplexed immunoassay.

Although the average CVs in this initial multiplexed demonstration (11 ± 8% for the FSH assay across all sensors that measured concentrations from 2.5-50 ng/mL, and 19 ± 15% for the PdG assay across all sensors that measured concentrations from 0-1000 ng/mL) do not yet match the performance of a mature commercial PdG lateral flow assay (average CV of 5.05%)^47^, they nevertheless establish reproducible dose–response behavior and confirm the feasibility of simultaneous hormone quantification on a silicon photonic platform. Importantly, the assay’s current variability reflects early-stage surface chemistry and flow-handling limitations rather than fundamental limits of the technology, and therefore represents a clear avenue for targeted optimization. Traditional LFAs are challenging to multiplex beyond 2-3 analytes^18^ due to factors including the physical limitations of fitting multiple test lines and controls on a test strip, influence of multiple test lines on the sample flow rate and assay time, complexity of readout, and cross-reactivity^89^. These multiplexing challenges have been highlighted as a challenge for hormone LFAs^22^. Sensitivity has also been a historical challenge for LFAs^18^. Although strategies have been proposed to improve both multiplexing^20,89^ and sensitivity^90^, these remain research efforts with tradeoffs in many cases. A recent review has reported that LoDs for commercial LFAs for urinary FSH detection range from 25-40 mIU/mL (8.475-13.56 ng/mL)^22^ – poorer than the LoD reported for our initial proof-of-principle by a factor of >14.

Future work will focus on building upon this promising initial demonstration by improving replicability, for example by exploring alternate immunoassay stabilizer protocols and blocking strategies, incorporating an initial regeneration step, quantifying the replicability of different stages of our functionalization strategy (e.g., PDA coating, bioreceptor loading, blocking), and performing orthogonal testing of sample solutions.

### 2.5 Inkjet spotting extends functionalization precision and throughput for highly multiplexed hormone biomarker detection

To assess the feasibility of highly multiplexed hormone marker detection on our SiP platform, we performed multiplexed FSH and PdG assays on SiP sensors functionalized via piezoelectric inkjet printing-mediated capture agent spotting (Figure 6). Unlike manual pipette spotting, which has millimeter-scale resolution, inkjet printing can be used to pattern tens of different receptors on a single sensor chip for detection of tens of hormone targets, effectively leveraging the multiplexability of SiP MRR arrays. Piezoelectric inkjet printing employs piezoelectrically actuated glass capillaries to dispense droplets with volumes on the order of 1–100 pL at specified locations on the substrate with xy spatial accuracies on the order of 10 µm^30^. Piezoelectric inkjet printing has previously been used to pattern protein-^91–93^, peptide-^94^, polymer-^95^, and glycoconjugate-based^27^ receptor chemistries on SiP sensors. This technique’s non-contact nature eliminates the risk of damage to waveguides during patterning, while its high resolution (∼10–100 µm) allows for resonators to be individually addressed during functionalization for detection of different targets on adjacent resonators.

**Figure 6.**
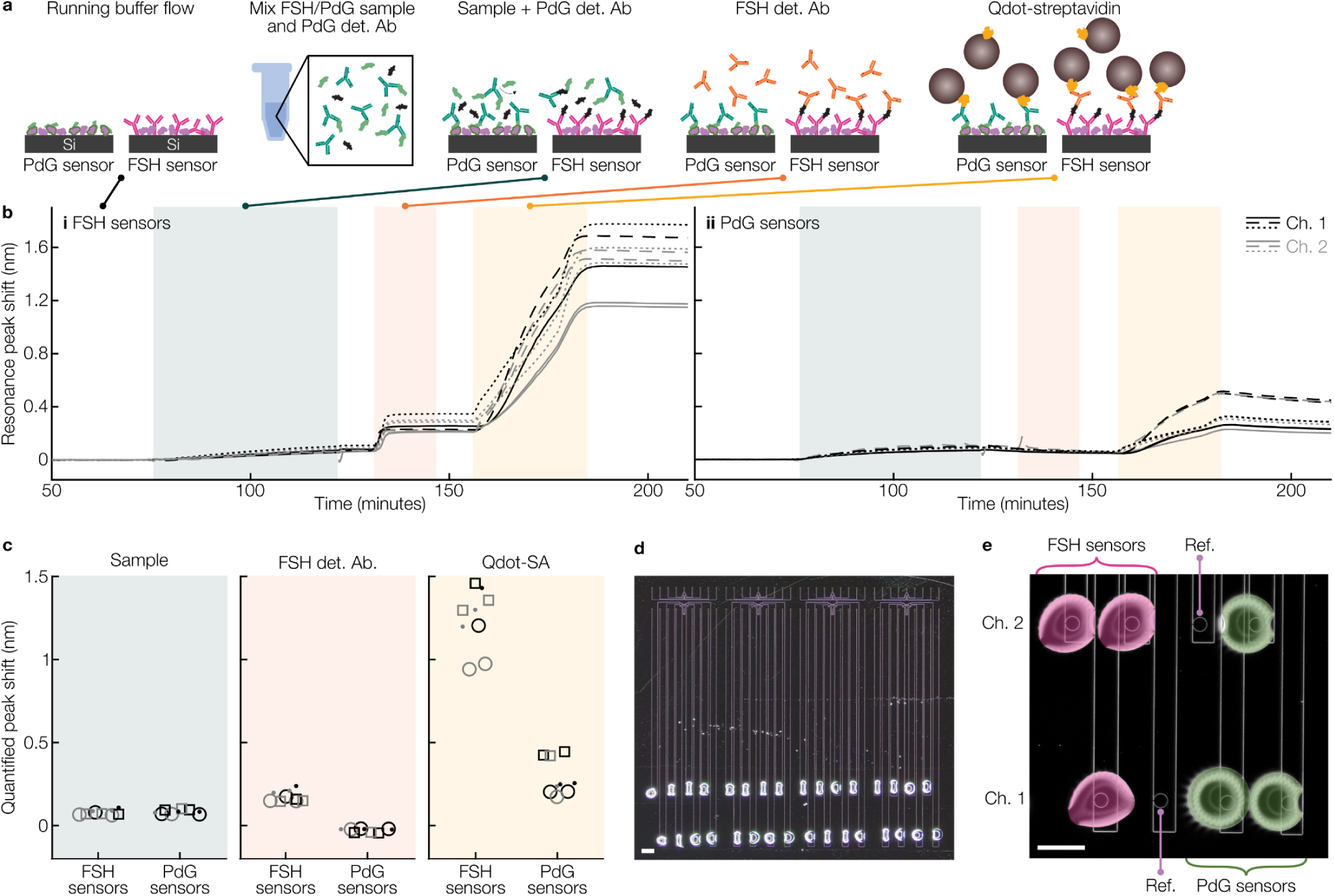
Proof-of-principle towards highly multiplexed hormone detection using inkjet-spotted silicon photonic resonator sensors. (a) Cross-sectional schematics illustrating the stages of the multiplexed binding assay for FSH and PdG detection, including sample delivery (artificial urine containing 25 ng/mL FSH and 0.05 μg/mL PdG, mixed with PdG detection antibody), FSH detection antibody binding, and Qdot-mediated amplification. (b) Reference-subtracted sensorgrams depicting the sensor signal throughout the multiplexed detection assay for the FSH sensors (left) and PdG sensors (right). (c) Quantified reference-subtracted resonance peak shift for the FSH and PdG sensors during the sample/PdG detection antibody, FSH detection antibody, and amplification assay stages. Each datapoint represents the quantified signal from one sensor. Marker colours represent different fluidic channels and marker styles denote assay replicates (N = 3 assay replicates, each with 3 sensor replicates per target). (d) Micrograph of an inkjet-spotted SiP chip (scale bar represents 400 µm). All 8 resonators in each of the 4 replicate sensor groups were spotted with a 25 µg/mL antibody solution containing 10% glycerol and 0.005% Triton X-100 in PBS to test the feasibility of accurate protein ink dispensing on adjacent resonators. (e) Annotated micrograph showing zoomed-in view of inkjet-printed capture agent spots, aligned with microring resonator sensors, on one device group of the SiP chip used for trial 2 (scale bar represents 200 µm). False colouring is used to differentiate the FSH capture antibody and PdG-BSA spots.

For these multiplexed assays, we used sensor chips designed with 4 groups of 8 simultaneously-addressable SWG MRRs, split equally across 2 microfluidic channels. This chip design (Supplementary Information Section 8.1) features close inter-resonator spacing within each 8-resonator group (240–288 µm between adjacent MRRs), requiring precise capture agent spotting to achieve multiplexed detection. These chips were first coated with PDA, then spotted with nL-scale spots (5–7 × 100 pL-scale droplets per resonator) of capture agent solutions using a custom-built piezoelectric inkjet printer (Supplementary Information Section 8.5). As with pipette spotting, capture agent solutions were prepared in PBS with 0.005% Triton X-100 and 10% glycerol ^96^. The Triton X-100 surfactant was included to reduce inkjet nozzle fouling, prevent the coffee ring effect, and encourage uniform receptor loading across the spotted region. The humectant glycerol was included to slow droplet drying. In each 8-resonator group, 3 resonators were spotted with FSH capture antibody, 3 were spotted with PdG-BSA, and the remaining 2 were left bare and used as reference sensors (Figure 6(d-e)). The chip was incubated with the spotting solutions for 1 hour. Based on order-of-magnitude mass transport calculations modelling this system, we expect that the capture agent concentrations used here were in large excess of what is required to achieve saturation of the spotted regions of the sensor surface within the 1-hour incubation time ^44,72^. The chip was then blocked, coated with immunoassay stabilizer, dried, and integrated with microfluidics, as for the experiments reported in Sections 2.3-2.4. Single-cycle assays were performed, as in Section 2.4, using FSH and PdG concentrations of 25 ng/mL and 0.05 µg/mL, respectively. Figure 6(a-b) shows the reference-subtracted sensorgrams for the (i) FSH and (ii) PdG sensors in N = 3 replicate assays performed on inkjet-spotted chips. Figure 6(c) presents the quantified reference-subtracted resonance peak shifts for the (i) sample, (ii) FSH detection antibody, and (iii) quantum dot amplification assay stages on the FSH and PdG sensors. Brightfield microscopy was used to verify deposition of discrete spots of the two bioreceptor “inks” on the targeted resonators, as demonstrated in Figure 6(e).

We see signals at each assay stage (Figure 6(b)) that are similar to those observed in the single-target assays and multiplexed assays performed on pipette-spotted chips, indicating specific detection of the two analytes. Across three replicate assays, the average signals for the FSH sensors are 0.078 ± 0.015 nm, 0.175 ± 0.033 nm, and 1.240 ± 0.183 nm, and those for the PdG sensors are 0.083 ± 0.012 nm, -0.028 ± 0.011 nm, and 0.290 ± 0.108 nm for the sample, FSH detection antibody, and amplification assay stages, respectively. For two of the assay replicates, the reference sensors exhibited negligible binding signals for all assay stages, as expected. However, for one assay replicate, one reference sensor exhibited positive binding signals for the FSH detection antibody (0.036 nm) and amplification (0.477 nm) assay stages. While no droplets were visible on this resonator during microscope examination after inkjet spotting, these results suggest that this resonator was contaminated with satellite droplets of FSH capture antibody solution during spotting. Reference sensor contamination could be mitigated by spotting these sensors with blocking solution prior to printing other reagents. Motivated by these results, future work will focus on further optimizing droplet actuation waveforms and ink formulations and testing nozzle coatings to achieve more reliable satellite-free inkjet printing^96–98^, as well as leveraging the capability for highly multiplexed functionalization afforded by inkjet spotting to develop assays to simultaneously detect a larger number of hormone targets.

Similarly to the multiplexed assays employing pipette-based functionalization (section 2.3), we observe considerable inter-assay variability in binding signals, with the FSH and PdG sensors exhibiting inter-assay CVs of 15% and 37%, respectively, during the amplification assay stage, supporting the plan for future development to improve assay replicability. Compared to assays performed on pipette-spotted chips using the same FSH and PdG concentrations, amplification signals were significantly lower for the inkjet-spotted FSH sensors and higher for the inkjet-spotted PdG sensors (p < 0.05, Mann-Whitney U-test). Given the opposite trends observed for the two analytes, combined with the relatively low number of assay replicates, it is possible that these differences in amplification signal do not reflect differences in assay sensitivity associated with spotting modality, but instead, can be attributed to other experimental factors contributing to inter-assay variability. Overall, these results demonstrate the feasibility of hormone marker detection on inkjet-spotted SiP MRR sensors, highlighting a path toward future development and optimization of highly multiplexed hormone assays on this platform.

## 3. Discussion

We have demonstrated, for the first time, multiplexed detection of a pilot set of two hormone markers relevant to women’s health in artificial urine using silicon photonic biosensors. In our multiplexed immunoassay, the sandwich assay used to detect FSH and the antigen-down competitive binding assay used to detect PdG are both amplified using a commercially available quantum dot conjugate, demonstrating amplification factors of 18.7 ± 1.9 at 25 ng/mL FSH, and increasing with decreasing FSH concentration. We have shown regeneration of the sensors’ surface functionalization in the multiplexed assay using a low-pH buffer solution; however, further optimization is needed to improve the cycle-to-cycle replicability after regeneration and ensure that all bound material is removed from the surface bioreceptors. This optimization, which is expected to be bioreceptor-dependent, could include tuning the regeneration time, pH, and ion concentrations, as well as investigating high pH (e.g., NaOH-based) regeneration solutions^99,100^. The multiplexed assay shows promise for hormone quantification, detecting FSH levels within the clinical range and PdG levels in the sub-clinical range. Optimization of detection antibody and amplification solution concentrations, as well as further dilution of the sample, could improve alignment of the detection capability with the clinical range in urine. Quantification performance is limited by the replicability of the detection signal; systematic evaluation of immunoassay stabilizers, blocking conditions, and other factors influencing inter-assay variability will be the focus of future optimization efforts^44^.

Despite these limitations, we demonstrate that two distinct assay formats can detect two classes of hormone biomarkers from the same sample on a single millimeter-scale chip – highlighting the strong potential for multiplexed hormone quantification using silicon photonic biosensors to meet important clinical needs. Beyond further optimization of these two pilot assays, future work could include expanding the multiplexing capability of the system to additional targets (e.g., E3G, LH, E1C, estradiol, estrone) using multiplexed inkjet functionalization of tens of sensors on the same chip^27,30^. Towards low-cost at-home quantification systems, future work could also include hormone measurement using photonic sensor architectures optimized for miniaturization of the readout system and laser^31–33^. Ultimately, with continued development and validation, putting this type of multiplexed quantification system in the hands of patients and women’s health researchers will likely afford new opportunities to study hormone dynamics and support data-driven insights in diagnoses and personalized medicine.

## 4. Methods

This section briefly describes our experimental methods. Additional details and detailed fluidic protocols for each assay are included in Sections 8-9 of the Supplementary Information.

### 4.1 Sensor design and readout

SiP sensing chips were designed to accommodate 2-plex measurements alongside a group of reference sensors to facilitate reference-subtraction to reduce the effects of nonspecific binding, temperature fluctuations, and bulk refractive index changes in the fluid. The chips were designed to measure 2 samples simultaneously (via 2 parallel microfluidic channels) and to permit simple functionalization via pipette spotting. The microring resonator (MRR) sensors were designed as previously described by our group^43,44,71^ using sub-wavelength grating waveguides with the following design parameters: ring radius *R* = 30 µm, coupling gaps *g_c_* = [500, 550] nm, grating period *Λ* = 250 nm, duty cycles *δ* = [0.65, 0.7], waveguide width *w* = 500 nm, and waveguide thickness *t* = 220 nm. The photonic integrated circuit included 12 sensors read out simultaneously (2 sensors in each of the 3 functionalization spots, in each of the 2 microfluidic channels). Section 3 of the Supplementary Information presents the quantified intrinsic bulk refractive index sensitivity (S_bulk_) of each sensor measured in the FSH, PdG, and multiplexed binding assays presented here.

A second SiP sensor chip layout, previously described by our group^43,44^, was used for multiplexed sensing experiments employing inkjet spotting-based functionalization. The compact inter-resonator spacing on this layout (240–288 µm between adjacent MRRs) precludes multiplexed functionalization via manual pipette spotting, demonstrating how the added precision and accuracy of an inkjet dispense system can enable a higher degree of multiplexing in a smaller photonic IC area. In this design, 8 MRRs, split equally across 2 fluidic channels can be simultaneously read out. The MRRs employ sub-wavelength grating waveguides with the same parameters as the sensor chips designed for pipette spotting.

Photonic sensor chips were fabricated on silicon-on-insulator (SOI) wafers through the Applied Nanotools Inc. (ANT, Edmonton, AB, Canada) NanoSOI Fabrication Service Silicon Device Layer process, which uses 100 keV electron beam lithography and reactive ion etching^101^. The chips were integrated with poly(dimethylsiloxane) (PDMS) microfluidics (channel width *w* = 300–400 µm and height *h* = 300 µm) as previously described^43,44,71^. Schematics of the photonic integrated circuit and microfluidics designs are included in section 8.1 of the Supplementary Information.

The optical transmission spectra of the sensor circuits was read out using a custom optical testing setup (Maple Leaf Photonics, Seattle, WA, USA), a 12-channel lidless fiber array (VGA-12-127-8-A-14.4-5.0-1.03-P-1550-8/125-3A-1-1-0.5-GL-NoLid-Horizontal, OZ Optics, Ottawa, ON, Canada), a C-band swept-tunable laser (Agilent 81682A, Agilent Technologies, Santa Clara, CA, USA), and optical detectors (Agilent 8164A and Keysight N7744C, Keysight Technologies, Santa Rosa, CA, USA), as previously described^43,44,71^.

### 4.2 Sensor functionalization

#### 4.2.1. Polydopamine coating

MRR sensors were functionalized with FSH capture antibodies (FSH sensors) or PDG-BSA conjugate (PdG sensors) using a covalent polydopamine (PDA)-mediated attachment protocol that our team has previously described for other detection assays^44^. Briefly, photonic sensor chips were solvent-cleaned via sequential immersion in acetone, isopropanol, and ASTM Type I ultrapure water and dried with nitrogen. A PDA film of ∼1.8 nm was subsequently deposited using a one-pot aqueous reaction, in which 1-3 sensor chips were placed in a glass crystallization dish with 40 mL of freshly prepared dopamine hydrochloride (Sigma H8502-5G, lot BCCJ9540) solution (2 mg/mL in Tris buffer (0.5 M, pH 8.6, Thermo Scientific AAJ62287AP, lots P24K523, M16L509)) for 30 minutes. The solution was uncovered and continuously stirred at ∼100 RPM to facilitate oxygen transport. The chips were then rinsed in fresh Tris buffer and ultrapure water, and dried with nitrogen. After PDA-coating, chips were stored in the dark under ambient conditions for 1-6 days prior to functionalization.

#### 4.2.2. Manual spotting

Functionalization was performed by manually spotting each of the 3 groups of resonators on each chip with 1-2 μL of functionalization solution. For all bioreceptor solutions, the spotting buffer contained 10% glycerol (Fisher Scientific G33-4, lot 116960) and 0.005% Triton X-100 (Sigma Aldrich T8787-100ML, lot SLCJ6163) in pH 7.4 phosphate-buffered saline (1× PBS, Gibco 10010-023). FSH sensors were spotted with 1 mg/mL FSH capture antibody (BiosPacific mouse monoclonal IgG1 A18054501P, lot A7942). PdG sensors were spotted with 1 mg/mL PdG-BSA (BiosPacific V56131314, lot V0143). Reference sensors were spotted with 1 mg/mL BSA (Sigma A706-50G, lot 0000296702). After spotting, the chip was incubated at room temperature in a humid chamber (35 mm petri dish with a wetted cleanroom wipe lining the bottom) for 60 minutes. The chip was then rinsed with a wash buffer (0.05% Tween 20 (Fisher Scientific BP337-500, lot 194435) in PBS) immediately prior to blocking.

#### 4.2.3. Inkjet spotting

Functionalization was performed by dispensing ∼100 pL-scale droplets of functionalization solution on individual resonators using a custom-built piezoelectric inkjet printer (described in section 8.5 of the Supplementary Information). The spotting solution formulations were the same as for manual spotting (described in section 4.3.2). The three leftmost resonators (1–3) in each 8-resonator cluster were spotted with FSH capture antibody, the two middle resonators (4, 5) were left unspotted to act as references, and the three rightmost resonators (6-8) were spotted with PdG-BSA. This yielded 1 FSH sensor, 1 reference sensor, and 2 PdG sensors in fluidic ch. 1 and 2 FSH sensors, 1 reference sensor, and 1 PdG sensor in fluidic ch. 2. The actuation waveworms used to trigger droplet ejection were manually tuned immediately before printing. For both spotting solutions, a bipolar trapezoidal waveform with a high voltage of 2.40–3.30 V, high dwell time of 25.0–28.5 µs, low voltage of -2.00 V, low dwell time of 3.0 µs, and rise/fall time of 3.0 µs was generated by an arbitrary waveform generator (Keysight 33500B) and amplified 20× by a linear amplifier (PiezoDrive PD200), prior to being delivered to the dispensing nozzles (80 µm orifice, MicroFab Technologies Inc. MJ-AL-00-080). A total of 5–7 × 100 pL-scale drops of spotting solution were printed on each resonator. Spotted chips were visualized under an optical microscope to verify suitable alignment of capture agent spots with the targeted resonators. After spotting, the chip was incubated at room temperature in a humid chamber (plastic box with a wetted cleanroom wipe lining the bottom) for 60 minutes. The chip was then thoroughly rinsed with a wash buffer immediately prior to blocking.

#### 4.2.4. Blocking

For single-hormone assays and pilot multiplexed detection assays, the chips were blocked using a 60-min incubation in 20 mg/mL BSA in PBS followed by a wash buffer rinse, 15-min incubation in immunoassay stabilizer (Sigma S0950-1L, PCode 1003667371, Source MKCS8259), and nitrogen dry. For multiplexed quantification assays and multiplexed detection assays on inkjet-spotted chips, we used a combination of protein and polymer blocking, as previously found to yield improved blocking of interferometric sensors for tuberculosis detection in urine specimens compared to either protein-based or polymer-based blocking alone^88^. The chips were blocked using a two-step process. First, 100-200 μL of 0.25 mg/mL solution of poly-L-lysine grafted with polyethylene glycol (PLL-g-PEG, SuSoS Surface Technologies PLL(20)-g[3.5]- PEG(5), lot NB03-53) in HEPES buffer (10 mM HEPES and 150 mM NaCl in ultrapure water, adjusted to pH 7.4), and incubated for 60 minutes in the humid chamber. The chip was then briefly rinsed with wash buffer and immersed in the 20 mg/mL BSA blocking solution for a further 60 minutes, rinsed with wash buffer again, incubated in immunoassay stabilizer for 15 minutes, and dried on the benchtop at ambient conditions for 30 minutes. All chips for this work were used immediately after sensor functionalization.

### 4.3 Hormone-detection assays

The photonic chip was integrated with PDMS microfluidics immediately after functionalization and drying, using gasket treatments and surfactant-based channel prewetting to reduce the likelihood of bubble formation during the assay as previously described^44^. Briefly, cured PDMS gaskets were degassed overnight in a desiccator and treated with air plasma immediately prior to integration with the photonic chip. All 10 fluidic lines in each channel of our two-channel modular Fluigent LineUp™ series fluid control system (configuration previously described in detail^44^) were primed with fluid prior to connecting the microfluidic device, and the fluidic control system was used to deliver 0.3 mM Triton X-100 in PBS at a slow flow rate of μL/min to wet the channel crevices, reducing the likelihood of formation of Harvey nuclei where channel crevices and defects are incompletely wetted (which serve as recurrent bubble-nucleation sites during long assays^44,102^). After ∼5 minutes of flow of the pre-wetting solution, we switched to flowing assay running buffer (ARB; 0.1 mg/mL BSA and 0.05% Tween 20 in PBS) at a flow rate of 30 µL/min. The photonic chip was optically coupled to the readout system, the automated fluidic delivery protocol was started, and the spectra from 12 resonator devices was read out continuously during the assay (approximately 25 s per sweep).

Hormone-detection assays involved sequential delivery of: (1) nonspecific binding challenge solutions (1 mg/mL BSA in ARB, for single-hormone and pilot multiplexed detection assays); (2) sample solutions (FSH and/or PdG in 50% artificial urine and ARB; PdG and multiplexed detection assays also included 1-5 µg/mL PdG detection antibody in the sample solution); (3) 2 µg/mL FSH detection antibody in ARB for FSH and multiplexed detection assays; (4) 0.005-0.01 µM Qdot-streptavidin conjugate in ARB; (5) sensor regeneration buffer (10 mM glycine, 160 mM NaCl, adjusted to pH 2.2 with HCl). ARB was flowed for 10 minutes between each assay step to rinse away unbound reagents and permit quantification, and ARB was flowed for 30 minutes at the beginning and end of each assay cycle to quantify stability and permit baseline correction.

Multiple binding cycles were tested for single-hormone and pilot multiplexed detection assays. Following each binding assay, the bulk refractive index sensitivity (S_bulk_) of each sensor was characterized by sequentially delivering a sequence of three NaCl solutions that served as refractive index standards. Two ramps of the standard solutions were consecutively delivered (each consisting of stepping up the concentration/refractive index then stepping back down to water), and S_bulk_ was reported as the average of the duplicate measurements. Details for all reagent solutions and flow protocols for each assay type are provided in sections 8.2-8.3 of the Supplementary Information.

### 4.4 Data analysis

The sequentially acquired optical spectra for each experiment were analyzed as previously described to extract the resonance peak shift vs. time for each optical sensor^44^. Briefly, the resonance peaks for each optical spectrum were identified and Lorentzian-fitted to extract the central resonance wavelength and other Lorentzian fit parameters for the ∼3 peaks in each sweep range for each sensor using a custom Python software GUI. The vectors of Lorentzian fit parameters for each resonance peak in each sweep were compared with those from sequentially acquired sweeps (using either cosine similarity or Euclidean distance measures) to track the shift in the resonance wavelength of each resonance peak vs. time in the assay. The average shift across all quantified resonance peaks in each spectrum (typically 3-4 peaks per spectrum) was quantified as the resonance peak shift vs. time. The use of Lorentzian fitting reduces the impact of intensity noise and permits higher-resolution identification of the central wavelength, while the use of cosine similarity or Euclidean distance for all resonance peaks (vs. tracking only the central wavelength from one resonance peak) facilitates quantification that is more robust to noise, spectral nonidealities, and large resonance shifts.

After quantifying the resonance peak shift vs. time (sensorgram data) for each monitored sensor, reference-subtraction was performed. For single-hormone assays and pilot multiplexed detection assays, the shifts for the 2 reference sensors in each microfluidic channel were averaged and subtracted from the shifts of the remaining 4 sensors in that channel using a custom MATLAB script. For multiplexed detection assays on inkjet-spotted chips, shifts for the single reference sensor in each channel were subtracted from the shifts of the remaining 3 sensors in that channel. In the second assay replicate, however, analyte binding was observed on the ch. 1 reference sensor, likely due to contamination of the sensor by a satellite droplet of functionalization solution during inkjet printing. As such, data from the contaminated resonator were omitted and reference-subtraction for that trial was performed using shift data for the ch. 2 reference sensor only. Reference-subtraction effectively reduces the effects of bulk refractive index changes, surfactant effects, temperature effects, and nonspecific binding, as shown in section 8.4 of the Supplementary Information; however, some differences remain in the sensorgram data (e.g., during sensor regeneration), potentially due to differences in conformational changes between the reference and sensor biofunctionalized surfaces. Some differences in nonspecific binding performance are also expected due to the difference between the reference probe (BSA in this work) and functionalized surfaces (capture antibody or PdG-BSA). Previous work has found that optimization of the negative control probe selection for each bioreceptor can improve these kinds of effects^103^.

The reference-subtracted sensorgram data were baseline-corrected using a custom MATLAB script as previously described^44^ to mitigate the effects of any sensor drift and functionalization dissociation on the subsequent quantification of peak shifts and baseline drift slopes for each assay stage. The sensorgram peak shift (*y*) vs. time (*t*) data for an initial ∼15-min ARB equilibration period in each binding cycle was nonlinear least-squares fitted to a modified bimolecular dissociation model^104^ incorporating additional fit coefficients for equilibrium (*t* = ∞) peak shift (*d*) and time offset (*c*) in addition to the off-rate of the binding interaction (*c*) and the maximum signal from the bound fraction (*a*):

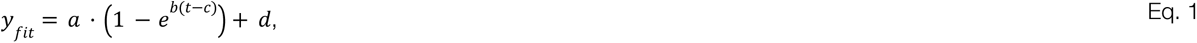

This fit function was subtracted from the sensorgram data for the binding cycle, and the resulting reference-subtracted and baseline-corrected data were postprocessed using a custom semi-automated MATLAB script to extract the quantified reference-subtracted peak shifts for each assay stage, as previously described^44^. Plots of the reference-subtracted and baseline-corrected data were also overlaid with coloured boxes to visualize the stages of the assay using another custom MATLAB script, using log files from the automated fluid control system. The start and end time points for these overlay regions were defined based on the log files as well as a delay factor that took into account the integrated measured flow rates (from the fluid control system log files) and the estimated internal volume between the fluidic switch and the microfluidic gasket (approximately ∼150 µL for our system). Additional details on our data analysis workflows are reported in our previous works and their corresponding supplementary information^43,44^.

### 4.5 Hormone quantification

The hormone concentration-series data presented in section 8.5 were assessed to investigate the sensors’ quantitative performance by performing calibration curve fitting and extracting the relative uncertainty of quantified concentrations from the calibration^105–107^. We used a 4-parameter logistic model (commonly used for immunoassay calibration^86^) to calibrate both the FSH and PdG sensor data, and performed the curve fitting, uncertainty quantification, and concentration quantification from sensor peak shift signal using custom MATLAB scripts. Details regarding the procedures for calibration and the tested models are provided in section 9 of the Supplementary Information.

## Data and code availability

Data supporting this article have been included as part of the Supplementary Information. Additional data and code will be provided upon request.

## Supporting information

Electronic Supplementary Information

## Acknowledgements

The authors are grateful to work and live on land that is the traditional, ancestral, and unceded territory of the Coast Salish Peoples, including the territories of the Musqueam, Squamish, and Tsleil-Waututh First Nations. The authors sincerely appreciate contributions, feedback, and helpful discussions from all members of our interdisciplinary biosensor collaboration team, as well as from the Dream Photonics Inc. team, whose efforts helped enable this work. We gratefully acknowledge immensely helpful discussions with Michael Williams from the UBC Antibody and Biologics Core Facility (AbLab™) during the course of this assay development, as well as protein conjugation services by AbLab™. The authors are also grateful for SEM imaging by Sheri Jahan Chowdhury, data acquisition and analysis software contributions from Avineet Randhawa, Mohammed Al-Qadasi, Yifei Liu, and Piramon Tisapramotkul, and piezoelectric inkjet dispenser development contributions from Abdel-Rahman AL Faouri and Spencer Witt. The authors acknowledge project and fellowship support from the Schmidt Science Polymaths Program, Optica Foundation Challenge, School of Biomedical Engineering Trainee Research Awards program, Mitacs, UBC Expansion Funding for Canada’s Digital Technology Supercluster (CDTS) projects, Natural Sciences and Engineering Research Council of Canada (NSERC), Killam Trusts, Silicon Electronics-Photonics Integrated Circuits Fabrication (SIEPICfab) consortium, CMC Microsystems, and Canada’s National Design Network. We also acknowledge undergraduate research funding supplements from the UBC Work Learn International Undergraduate Research Awards, Faculty of Medicine Multidisciplinary Research Program in Medicine, Centre for Blood Research Undergraduate Summer Research Program, NSERC Undergraduate Student Research Awards, and School of Biomedical Engineering Synergy Research Program.

## Author contributions

**Conceptualization**, S.M.G., L.C., S.S., and K.C.C.; **Data curation**, S.M.G., M. Wang, L.S.P., and M. Wei; **Formal analysis**, S.M.G., M. Wang, L.S.P., and M. Wei; **Funding acquisition**, S.M.G., L.C., S.S., and K.C.C.; **Investigation**, S.M.G., M. Wang, L.S.P., M. Wei, K.N., and S.C.; **Methodology**, S.M.G., M. Wang, L.S.P., M. Wei, K.N., B. C.-K., K.H., S.C., J.S., K.W., and N.T.; **Project administration**, S.M.G., L.C., S.S., and K.C.C.; **Software**, S.M.G., M. Wang, J.S., K.W., and Z.K.; **Resources**, S.M.G., L.C., S.S., and K.C.C.; **Supervision**, S.M.G., L.C., S.S., and K.C.C.; **Validation**, S.M.G., M. Wang, L.S.P., and M. Wei; **Visualization**, S.M.G., M. Wang, and L.S.P.; **Writing - original draft**, S.M.G., M. Wang, and L.S.P.; **Writing - review & editing**, S.M.G., M. Wang, L.S.P., M. Wei, K.N., K.H., S.C., L.C., S.S., and K.C.C.

