## Supplementary material for "Multiplexed hormone immunoassays for women’s health on a regenerable silicon photonic chip": Electronic Supplementary Information

### Multiplexed measurement of hormone biomarkers relevant to women's health using silicon photonic resonators

#### 1. Specific detection of 25 ng/mL FSH in artificial urine using a sandwich immunoassay

Seeking to demonstrate the regeneration performance and specificity of our sandwich assay using Qdot-mediated amplification, we next ran replicate experiments comparing detection of a clinically relevant 25 ng/mL concentration of FSH in a 1:1 mixture of artificial urine in immunoassay running buffer with that of a negative control (0 ng/mL FSH in the same 1:1 mix). We demonstrated 3 binding cycles of detection, regenerating the sensor in between binding cycles using a pH 2.2 glycine-HCl regeneration solution. The results of this experiment are shown in Figure S1.

Figure S1(b) depicts the reference-subtracted sensorgram traces for two assays, each with an FSH-containing and negative-control channel. Resonance peak shift signal is clearly visible for the FSH-containing channel only, at the detection antibody and Qdot amplification assay stages. Sensors in both channels show blue-shifts in the reference-subtracted resonance wavelength during regeneration, corresponding (as expected) to material being removed from the waveguide surface.

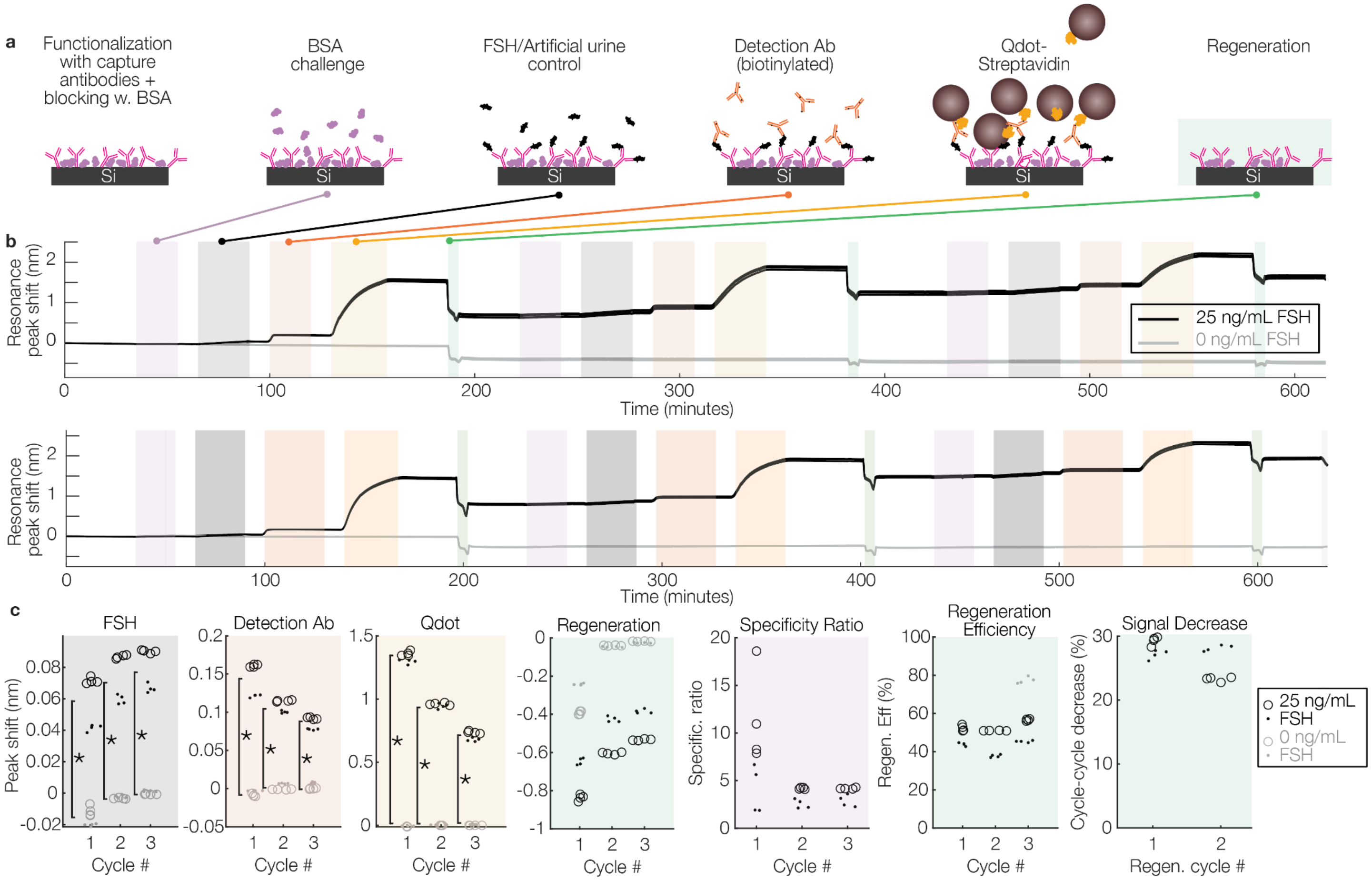

**Figure S1. Detecting 25 ng/mL FSH in artificial urine using silicon photonic resonator sensors.** (a) Cross-sectional schematics illustrating the stages of the sandwich assay for FSH detection, including specificity challenge, sample delivery (artificial urine control or artificial urine containing 25 ng/mL FSH), detection antibody binding, Qdot-mediated amplification, and sensor regeneration using pH 2.2 glycine-HCl. (b) Reference-subtracted sensorgrams depicting the sensor signal throughout 3 cycles of the detection assay. (c) Quantified reference-subtracted resonance peak shift sensor signal during the FSH, detection antibody, amplification, and regeneration assay stages, as well as computed specificity ratio (absolute value of 25 ng/mL FSH binding signal normalized to 1 mg/mL BSA binding signal) and regeneration metrics (regeneration efficiency and cycle-cycle decrease in Qdot signal). Each datapoint represents the quantified signal from one sensor (n = 4 sensors per assay with duplicate assays), and marker styles denote assay replicates. Sensor peak shift signal for FSH-containing samples is significantly different from that for negative control samples for the native FSH detection, detection antibody, and Qdot amplification assay stages (*$p < 0.001$, Mann-Whitney U-test). Section 3 of the Supplementary Information presents the quantified intrinsic bulk refractive index sensitivity ($S_{bulk}$) of each sensor measured in this binding assay.

Figure S1(c) presents the quantification of the binding and regeneration signals, as well as metrics of specificity and regeneration performance. Specific detection of FSH is demonstrated in two ways: firstly by comparing the reference-subtracted resonance peak shift signals in the FSH-containing and control channels, and secondly by computing the specificity ratio, which compares the specific detection shift from 25 ng/mL native/primary FSH binding to the nonspecific shift resulting from delivery of 1 mg/mL BSA (a 40,000-fold higher protein concentration). Across two assay replicates (n = 8 sensors/condition), we observed significantly higher binding shifts from the FSH-containing channels than from the control channels at the primary FSH, detection antibody, and Qdot binding stages in all three detection cycles, demonstrating specific detection of FSH in artificial urine both with and without signal amplification. For the FSH-containing samples, we measure CVs of 29%, 20%, and 16% for the primary FSH binding stage signal, 15%, 7%, and 9% for the FSH detection antibody signal, and 3%, 2%, and 5% for the Qdot stage signal for binding cycles 1-3, respectively (n = 8 sensors total, with n = 4 sensors in each of two duplicate assays for each condition). We observe average specificity ratios of 8 ± 5, 3.4 ± 0.9, and 3.5 ± 0.8 in detection cycles 1, 2, and 3, respectively, highlighting how, despite the 40,000-fold higher protein concentration, specific signal remains higher than nonspecific signal for all detection cycles for primary detection of FSH in artificial urine. The sandwich assay and signal amplification impart additional specificity to the assay, due to the higher detection signals and lower chance of nonspecific signal during these later assay stages (which contain assay reagents in buffer rather than a sample that could contain confounding proteins). Section 2 of the Supplementary Information presents a pilot comparison of FSH detection performance in matrices of increasing complexity (simple buffer, artificial urine, and artificial urine containing fetal bovine serum (FBS)), and demonstrates specific detection of 25 ng/mL FSH from the FBS-containing sample, using the signal from the FSH detection antibody and quantum dot assay stages.

We also evaluated regeneration performance by assessing 3 metrics of interest: the regeneration peak shift signal; the regeneration efficiency (defined as the regeneration shift divided by the sum of the shifts at the BSA, FSH, detection antibody, and Qdot assay stages, represented as a percentage: $\eta_{regen} = 100 \cdot \left|\frac{\Delta\lambda_{regen}}{\Delta\lambda_{BSA}+\Delta\lambda_{sample}+\Delta\lambda_{Ab}+\Delta\lambda_{Qdot}}\right|$), where $\Delta\lambda_{regen}$ represents the shift in resonance wavelength induced by the regeneration step (typically a negative (blue) shift), and $\Delta\lambda_{BSA}$, $\Delta\lambda_{sample}$, $\Delta\lambda_{Ab}$, $\Delta\lambda_{Qdot}$ represent the shifts in resonance wavelength measured in response to the nonspecific BSA challenge, FSH sample, detection antibody, and quantum dot assay stages (typically positive (red) shifts). We also evaluated the percentage by which the Qdot signal decreases between binding cycles. The regeneration efficiency gives a comparison of how much material is removed from the surface vs. how much was added during the binding assay; however, 100% regeneration efficiency does not necessarily mean that all bound material was removed because removal of the functionalization layers, blocking, and etching of the waveguides can all cause negative (blue) shifts in the resonance wavelength separate from the bound assay reagent removal. Denaturation or changes in protein conformation of the functionalization and blocking layers could also lead to changes in the thickness or refractive index of these layers, resulting in a positive or negative resonance shift. The percentage by which the Qdot signal decreases between binding cycles gives an indication of degradation in assay performance from cycle to cycle (e.g., due to functionalization denaturation or incomplete removal of bound material from the previous assay).

As expected, we observe larger-magnitude (more negative) regeneration shifts for the sensors exposed to 25 ng/mL FSH than for those exposed to the control channel, consistent with the bound assay reagents being removed from the resonator surface. We observe larger-magnitude regeneration shifts during the first regeneration cycle than those for the second or third, for all sensors. We hypothesize that this is due to the removal of loosely attached functionalization or blocking reagents from the surface of the sensors; an initial regeneration cycle (prior to the first binding assay) could be used to remove these materials and improve the regeneration consistency from cycle to cycle[1–3]. The regeneration efficiency for the sensors exposed to 25 ng/mL FSH was 48% ± 5% for the first cycle, 44% ± 7% for the second, and 51% ± 6% for the third. The cycle-cycle decrease in Qdot signal was 28.2% ± 1.3% for the first regeneration cycle (from binding cycles 1-2) and 26% ± 3% for the second regeneration cycle. These results suggest that the regeneration conditions used for this assay yield incomplete regeneration of the sensor surface, with remaining bound FSH, detection antibody, and/or Qdots after each binding cycle reducing the number of binding sites available on the surface for subsequent assays. This kind of signal decrease from cycle to cycle could potentially be calibrated for, and optimization of the regeneration conditions by tuning the regeneration buffer and regeneration time can be investigated in future work. Optimal regeneration conditions are often assay-specific, depending on the particular bioreceptor used for each assay, so careful optimization will be required for multiplexed assays[4,5].

Overall, the results highlighted in Figure S1 demonstrate successful specific detection of FSH in artificial urine, as well as regeneration of the sensor surface for multiple binding cycles, though future optimization of the regeneration conditions will be required to improve sensor performance for multi-cycle assays.

#### 2. FSH detection in complex media

The artificial urine recipe used in this work did not contain proteins[6], so we would expect higher-complexity human urine specimens to present more challenges with nonspecific binding. We thus sought to compare FSH detection performance of our on-chip sandwich immunoassay in matrices of increasing complexity. As a pilot investigation of the detection of FSH in matrices of increasing complexity, we compared FSH detection in (a) simple buffer, (b) artificial urine (AU), and (c) artificial urine containing 10% fetal bovine serum (FBS). FBS provides a high concentration of a range of serum proteins and has been previously added to artificial urine for in vitro cell-based assay applications[7]. Figure S2 presents the results of this investigation, comparing quantified reference-subtracted sensor signal at the FSH sample, detection antibody, and quantum dot assay stages for 5 conditions:

1. 25 ng/mL FSH in PBS buffer with 0.1 mg/mL BSA and 0.05% Tween 20 (assay running buffer or ARB)
2. 25 ng/mL FSH in 50% artificial urine (preparation details in Supplementary Section 3.21) in ARB
3. 25 ng/mL FSH in 50% artificial urine and 10% FBS in ARB
4. 0 ng/mL FSH (negative control) in 50% artificial urine in ARB
5. 0 ng/mL FSH (negative control) in 50% artificial urine and 10% FBS in ARB

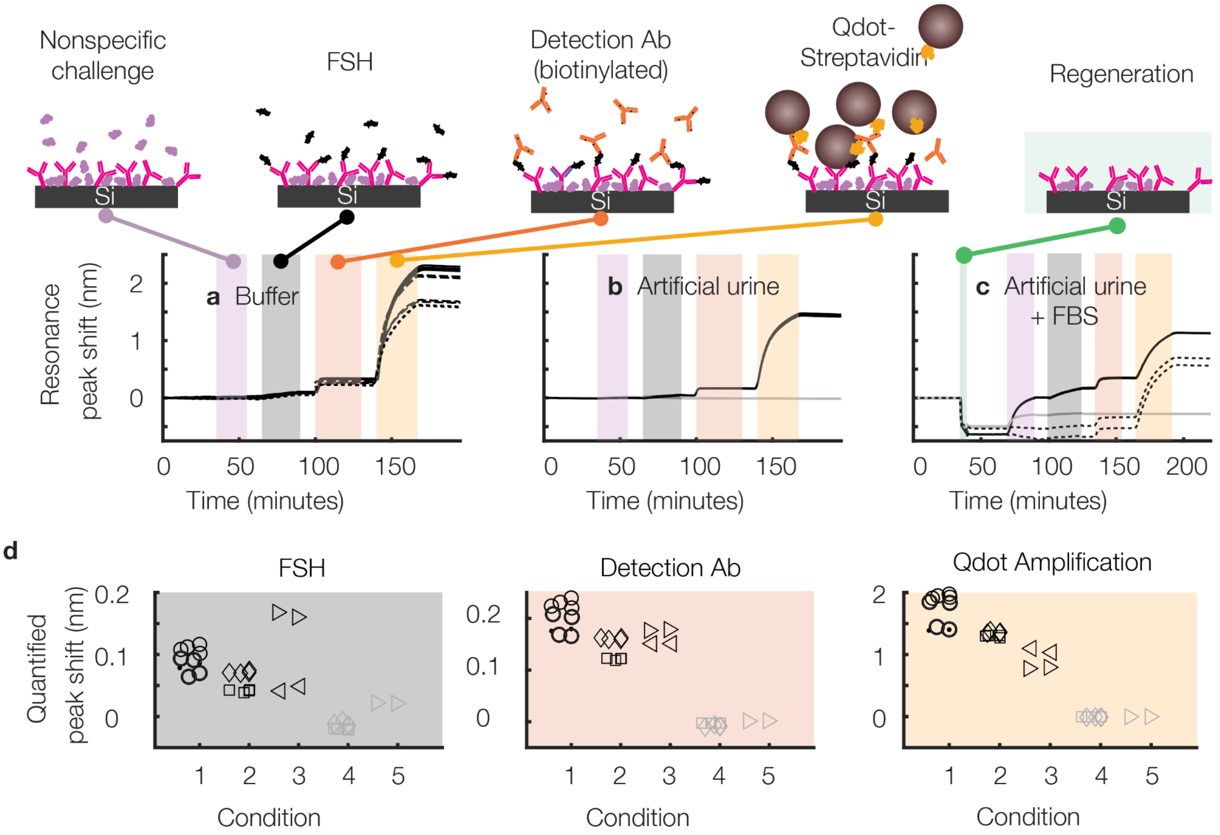

**Figure S2: Detecting FSH in matrices of increasing complexity**. (a-c) Reference-subtracted sensorgram plots depicting sensor signal for each stage of the FSH detection assay, for (a) simple PBS buffer with 0.1 mg/mL BSA and 0.05% Tween-20, (b) artificial urine (AU), and (c) AU spiked with 10% fetal bovine serum (FBS). Line styles denote channel replicates; black denotes samples containing 25 ng/mL FSH; grey denotes negative controls (0 ng/mL FSH). (d) Quantification of SiP sensor resonance peak shift signal for native FSH detection, detection antibody binding, and amplification for each tested condition (1. simple buffer, 2. AU, 3. AU+FBS, 4. AU negative control, 5. AU+FBS negative control). Marker types denote channel replicates; grey denotes negative control, markers denote replicates (N = 2–10 sensors/condition). The nonspecific challenge solution for the AU+FBS conditions included both 1 mg/mL BSA as well as 10% FBS.

The sandwich assay confers specific detection of FSH using both the detection antibody and quantum dot amplification assay stage signals for both the artificial urine matrix and the artificial urine matrix that includes additional confounding proteins from the FBS. However, the additional nonspecific binding from the sample during the primary (direct FSH binding) assay stage likely confounds specific detection using the primary signal only.

### 3. Intrinsic sensitivity measurements

Following each immunoassay, a series of salt solutions (serving as refractive index standards) were delivered to the sensors to assess their intrinsic sensitivity to bulk refractive index changes, as previously described. This section reports the intrinsic sensitivity data for the biosensors used in this work. The intrinsic sensitivity data for each sensor in all of the assays analyzed as part of the work presented in Figures 3-5 and Supplementary Figure S1 are reported in Tables S1-S4. Across all analyzed assays, the average $S_{bulk}$ was 400 ± 40 nm/RIU (inter-assay CV 10.5%). These values are in line with expectations for this type of SWG waveguide resonator. Our previous analysis of the intrinsic $S_{bulk}$ sensitivity of similar resonators with SWG duty cycle of 0.7 and a coupling gap of 500 nm reported $S_{bulk}$ of 439 ± 18 nm/RIU[3,8]. These previous experiments were run on bare, pre-functionalization silicon photonic chips. The slightly lower sensitivity and higher variability reported here is likely due in part to the design of these assays, in which the intrinsic sensitivity analysis was run after the binding assay (with functionalization and potentially residual proteins and Qdots remaining on the sensor surface after regeneration). The presence of these layers (forming a thin cladding layer of approximately ~10-30 nm) likely imparts a slight reduction in the waveguide's susceptibility to the surrounding fluid. Further contributing to the variability, three of these experiments (PdG assay trial 1, multiplexed quantification assays 4-5) used SWG waveguide ring resonators with an SWG duty cycle of 0.65; these are expected to have a slightly higher $S_{bulk}$ than the remainder of the assays, which used an SWG duty cycle of 0.7. Of the n = 17 assays using the SWG duty cycle of 0.7, we measure an average $S_{bulk}$ of 385 ± 26 nm/RIU (inter-assay CV 6.6%).

Table S1. Intrinsic sensitivity of the sensors used for FSH detection, for the data shown in Supplementary Figure S1. All trials used resonators with SWG duty cycle of 0.7.

| Trial | Average $S_{bulk}$ (nm/RIU) | Stdev $S_{bulk}$ (nm/RIU) | $S_{bulk}$ Intra-assay CV (%) | Channel | FSH conc. (ng/mL) | Sensor | $S_{bulk}$ (nm/RIU) |
|---|---|---|---|---|---|---|---|
| 1 | 377.50 | 4.71 | 1.25 | 1 | 0 | 1 | 376.72 |
| | | | | 2 | 25 | 2 | 379.43 |
| | | | | 1 | 0 | 3 | 376.53 |
| | | | | 2 | 25 | 4 | 382.34 |
| | | | | 1 | 0 | 5 | 370.92 |
| | | | | 2 | 25 | 6 | 378.96 |
| | | | | 1 | 0 | 7 | 376.60 |
| | | | | 2 | 25 | 8 | 387.34 |
| | | | | 1 | 0 | 9 | 375.08 |
| | | | | 2 | 25 | 10 | 380.36 |
| | | | | 1 | 0 | 11 | 370.08 |
| | | | | 2 | 25 | 12 | 375.63 |
| 2 | 409.77 | 6.19 | 1.51 | 1 | 0 | 1 | 393.47 |
| | | | | 2 | 25 | 2 | 407.04 |
| | | | | 1 | 0 | 3 | 412.93 |
| | | | | 2 | 25 | 4 | 405.84 |
| | | | | 1 | 0 | 5 | 415.61 |
| | | | | 2 | 25 | 6 | 407.51 |
| | | | | 1 | 0 | 7 | 417.28 |
| | | | | 2 | 25 | 8 | 412.08 |
| | | | | 1 | 0 | 9 | 413.13 |
| | | | | 2 | 25 | 10 | 412.95 |
| | | | | 1 | 0 | 11 | 410.34 |
| | | | | 2 | 25 | 12 | 409.11 |

Table S2. Intrinsic sensitivity of the sensors used for PdG detection, for the data shown in Figure 3. *Used resonators with SWG duty cycle of 0.65. All other trials used resonators with SWG duty cycle of 0.7.

| Trial | Average $S_{bulk}$ (nm/RIU) | Stdev $S_{bulk}$ (nm/RIU) | $S_{bulk}$ Intra-assay CV (%) | Channel | PdG conc. (µg/mL) | Sensor | $S_{bulk}$ (nm/RIU) |
|---|---|---|---|---|---|---|---|
| 1* | 412.70 | 3.41 | 0.83 | 2 | 0 | 1 | 411.93 |
| | | | | 1 | 10 | 2 | 413.09 |
| | | | | 1 | 10 | 3 | 411.47 |
| | | | | 1 | 10 | 4 | 415.20 |
| | | | | 2 | 0 | 5 | 416.79 |
| | | | | 2 | 0 | 6 | 419.17 |
| | | | | 1 | 10 | 7 | 407.93 |
| | | | | 2 | 0 | 8 | 406.69 |
| | | | | 1 | 10 | 9 | 412.13 |

| | | | | | | | |
|---|---|---|---|---|---|---|---|
| | | | | 2 | 0 | 10 | 413.35 |
| | | | | 1 | 10 | 11 | 411.58 |
| | | | | 2 | 0 | 12 | 413.11 |
| 2 | 340.51 | 5.26 | 1.55 | 1 | 0 | 1 | 334.48 |
| | | | | 2 | 10 | 2 | 345.83 |
| | | | | 1 | 0 | 3 | 337.12 |
| | | | | 2 | 10 | 4 | 347.95 |
| | | | | 1 | 0 | 5 | 336.94 |
| | | | | 2 | 10 | 6 | 341.88 |
| | | | | 1 | 0 | 7 | 346.16 |
| | | | | 2 | 10 | 8 | 344.61 |
| | | | | 1 | 0 | 9 | 335.30 |
| | | | | 2 | 10 | 10 | 343.86 |
| | | | | 1 | 0 | 11 | 332.08 |
| | | | | 2 | 10 | 12 | 339.96 |

Table S3. Intrinsic sensitivity of the sensors used for pilot multiplexed detection, for the data shown in Figure 4. All trials used resonators with SWG duty cycle of 0.7.

| | | | | | FSH Sensors | | | PdG Sensors | | |
|---|---|---|---|---|---|---|---|---|---|---|
| **Trial** | **Average $S_{bulk}$ (nm/RIU)** | **Stdev Sbulk (nm/RIU)** | **$S_{bulk}$ Intra-assay CV (%)** | **Channel** | **FSH conc. (ng/mL)** | **Sensor** | **$S_{bulk}$ (nm/RIU)** | **PdG conc. (µg/mL)** | **Sensor** | **$S_{bulk}$ (nm/RIU)** |
| 1 | 401.98 | 13.42 | 3.34 | 1 | 25 | 1 | 419.95 | 10 | 9 | 397.22 |
| | | | | 1 | 25 | 3 | 418.50 | 10 | 11 | 391.96 |
| | | | | 2 | 0 | 2 | 384.24 | 0 | 10 | 413.32 |
| | | | | 2 | 0 | 4 | 393.03 | 0 | 12 | 397.63 |
| 2 | 378.23 | 10.08 | 2.67 | 1 | 25 | 1 | 395.07 | 10 | 9 | 374.65 |
| | | | | 1 | 25 | 3 | 392.33 | 10 | 11 | 370.25 |
| | | | | 2 | 0 | 2 | 374.14 | 0 | 10 | 372.89 |
| | | | | 2 | 0 | 4 | 378.64 | 0 | 12 | 367.87 |

Table S4. Intrinsic sensitivity of the sensors used for multiplexed hormone quantification, for the data shown in Figure 5. *Used resonators with SWG duty cycle of 0.65. All other trials used resonators with SWG duty cycle of 0.7.

| | | | | | **FSH Sensors** | | | **PdG Sensors** | | |
|---|---|---|---|---|---|---|---|---|---|---|
| **Trial** | **Average $S_{bulk}$ (nm/RIU)** | **Stdev Sbulk (nm/RIU)** | **$S_{bulk}$ Intra-assay CV (%)** | **Channel** | **FSH conc. (ng/mL)** | **Sensor** | **$S_{bulk}$ (nm/RIU)** | **PdG conc. (µg/mL)** | **Sensor** | **$S_{bulk}$ (nm/RIU)** |
| 1 | 370.90 | 13.69 | 3.69 | 1 | 0 | 1 | 362.46 | 0 | 9 | 362.57 |
| | | | | 1 | 0 | 3 | 345.34 | 0 | 11 | 368.71 |
| | | | | 2 | 50 | 2 | 382.05 | 20 | 10 | 378.65 |
| | | | | 2 | 50 | 4 | 384.61 | 20 | 12 | 382.79 |
| 2 | 349.73 | 50.21 | 14.36 | 1 | 25 | 1 | 381.80 | 10 | 9 | 349.08 |
| | | | | 1 | 25 | 3 | 381.95 | 10 | 11 | 344.80 |
| | | | | 2 | 5 | 2 | 395.77 | 0.1 | 10 | 274.00 |
| | | | | 2 | 5 | 4 | 395.59 | 0.1 | 12 | 274.89 |
| 3 | 417.12 | 12.23 | 2.93 | 1 | 50 | 1 | 397.38 | 20 | 9 | 421.13 |
| | | | | 1 | 50 | 3 | 400.78 | 20 | 11 | 414.40 |
| | | | | 2 | 5 | 2 | 429.12 | 0.1 | 10 | 423.59 |
| | | | | 2 | 5 | 4 | 430.17 | 0.1 | 12 | 420.40 |
| 4* | 461.19 | 13.24 | 2.87 | 1 | 25 | 2 | 472.60 | 10 | 10 | 440.94 |
| | | | | 1 | 25 | 4 | 473.03 | 10 | 12 | 440.93 |
| | | | | 2 | 10 | 1 | 467.60 | 1 | 9 | 461.47 |
| | | | | 2 | 10 | 3 | 470.89 | 1 | 11 | 462.10 |
| 5* | 523.02 | 14.56 | 2.78 | 1 | 5 | 2 | 521.92 | 0.1 | 10 | 526.87 |
| | | | | 1 | 5 | 4 | 518.31 | 0.1 | 12 | 526.16 |
| | | | | 2 | 0 | 1 | 502.87 | 0 | 9 | 542.31 |
| | | | | 2 | 0 | 3 | 504.73 | 0 | 11 | 541.00 |
| 6 | 367.57 | 17.17 | 4.67 | 1 | 50 | 1 | 391.82 | 20 | 9 | 366.04 |
| | | | | 1 | 50 | 3 | 394.80 | 20 | 11 | 359.90 |
| | | | | 2 | 25 | 2 | 350.51 | 10 | 10 | 366.60 |
| | | | | 2 | 25 | 4 | 348.85 | 10 | 12 | 362.07 |
| 7 | 429.32 | 22.43 | 5.23 | 1 | 10 | 1 | 400.17 | 1 | 9 | 424.75 |

| | | | | | | | | | | |
|---|---|---|---|---|---|---|---|---|---|---|
| | | | | 1 | 10 | 3 | 402.54 | 1 | 11 | 424.05 |
| | | | | 2 | 0 | 2 | 430.62 | 0 | 10 | 465.54 |
| | | | | 2 | 0 | 4 | 433.54 | 0 | 12 | 453.32 |
| 8 | 403.11 | 32.00 | 7.94 | 1 | 25 | 1 | 427.60 | 10 | 9 | 365.92 |
| | | | | 1 | 25 | 3 | 425.45 | 10 | 11 | 361.39 |
| | | | | 2 | 10 | 2 | 434.19 | 1 | 10 | 388.19 |
| | | | | 2 | 10 | 4 | 439.63 | 1 | 12 | 382.53 |
| 9 | 367.09 | 8.94 | 2.44 | 1 | 50 | 1 | 362.99 | 20 | 9 | 358.09 |
| | | | | 1 | 50 | 3 | 366.48 | 20 | 11 | 355.67 |
| | | | | 2 | 25 | 2 | 367.76 | 10 | 10 | 379.11 |
| | | | | 2 | 25 | 4 | 365.93 | 10 | 12 | 380.69 |
| 10 | 410.40 | 13.60 | 3.31 | 1 | 7.5 | 1 | 403.17 | 0.02 | 9 | 395.65 |
| | | | | 1 | 7.5 | 3 | 406.98 | 0.02 | 11 | 392.98 |
| | | | | 2 | 2.5 | 2 | 430.25 | 0.3 | 10 | 414.37 |
| | | | | 2 | 2.5 | 4 | 427.63 | 0.3 | 12 | 412.18 |
| 11 | 410.21 | 22.52 | 5.49 | 1 | 2.5 | 1 | 440.05 | 0.3 | 9 | 404.19 |
| | | | | 1 | 2.5 | 3 | 438.09 | 0.3 | 11 | 378.17 |
| | | | | 2 | 7.5 | 2 | 413.67 | 0.02 | 10 | 396.96 |
| | | | | 2 | 7.5 | 4 | 422.34 | 0.02 | 12 | 388.18 |
| 12 | 371.10 | 18.04 | 4.86 | 1 | 3 | 1 | 377.09 | 0.05 | 9 | 396.21 |
| | | | | 1 | 3 | 3 | 375.66 | 0.05 | 11 | 395.60 |
| | | | | 2 | 2.5 | 2 | 359.81 | 0 | 10 | 354.24 |
| | | | | 2 | 2.5 | 4 | 360.78 | 0 | 12 | 349.36 |
| 13 | 391.75 | 4.58 | 1.17 | 1 | 0 | 1 | 393.58 | 0.02 | 9 | 395.33 |
| | | | | 1 | 0 | 3 | 394.52 | 0.02 | 11 | 392.60 |
| | | | | 2 | 3 | 2 | 392.06 | 0.05 | 10 | 385.20 |
| | | | | 2 | 3 | 4 | 396.52 | 0.05 | 12 | 384.22 |
| 14 | 360.29 | 16.21 | 4.50 | 1 | 6 | 1 | 380.58 | 0.08 | 9 | 367.45 |
| | | | | 1 | 6 | 3 | 382.45 | 0.08 | 11 | 365.98 |
| | | | | 2 | 6 | 2 | 341.46 | 0.08 | 10 | 352.69 |
| | | | | 2 | 6 | 4 | 342.32 | 0.08 | 12 | 349.41 |

#### 4. Calibration fit comparison

This section explores the selection process for the calibration model, including fitting curves on binding shift data from different assay stages, using calibration data from different ranges of concentrations, and using two different calibration function equations. In the final reported calibration curve, data from the Qdot-mediated amplification assay stage was used as it had the largest binding shifts, lowest CVs and relative CI widths, and were meaningful for both analytes. Figure S3 demonstrates calibration curves fitted using a 4-parameter logistic (4PL) on the sample delivery and FSH detection binding shifts. Qdot stage binding shifts from two different ranges of concentrations were fitted with both a 4-parameter and 5-parameter logistic (5PL) curves, and these models were quantitatively compared. The results of the comparison of these fits are presented in Figures S4-S5 and Tables S5-S6 below.

The final model that was chosen for both analytes (presented in Figure 5 of the manuscript) was the 4PL curve fitted to data from the complete range of concentrations. The models fitted on the calibration dataset with the complete range of concentrations had higher $R^2$ values and smaller relative CI widths than those fitted on the calibration dataset with the smaller range of concentrations. For both FSH and PdG, the 5PL models provided a slightly higher $R^2$ value, but weren't appropriate for the data. When fitted to the FSH data, the prediction function produced values with large imaginary components unless the slope parameter was fixed at 2. When fitted to the PdG data, the uncertainty on several of the predicted coefficients was very large. The 4PL models were chosen for both analytes as they have a similar $R^2$ to the 5PL model, a lower minimum relative CI width, and did not have the same issues as encountered with the 5PL models.

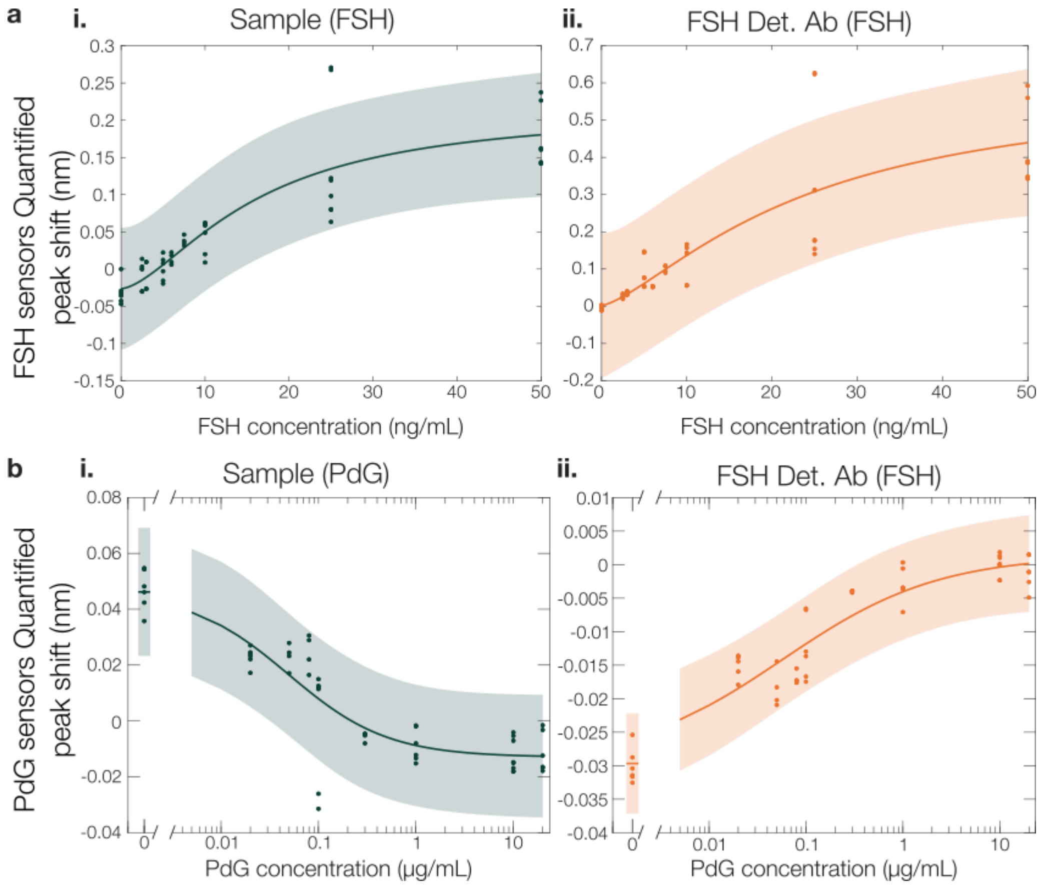

**Figure S3: Calibration curve fits for primary sample and secondary FSH detection binding assay stages**. Calibration curves for (a) FSH and (b) PdG sensors, fitted to peak shifts from (i) sample delivery and (ii) FSH detection antibody stages. (a(i)) FSH sensors' primary sample binding curve fit parameters: a = -0.02632 ± 0.02707 nm; b = 1.601 ± 1.0783; c = 16.07 ± 10.982 ng/mL; d = 0.214 ± 0.0985 nm; $R^2$ = 0.7882. (a(ii)) FSH sensors' FSH detection antibody curve fit parameters: a = 0.002244 ± 0.065234 nm; b = 1.411 ± 1.1941; c = 23.75 ± 29.801 ng/mL; d = 0.5924 ± 0.4571 nm; $R^2$ = 0.7358. (b(i)) PdG sensors' primary sample binding curve fit parameters: a = 0.04618 ± 0.00862 nm; b = 0.8552 ± 0.4762; c = 0.04943 ± 0.02954 ng/mL; d = -0.01297 ± 0.0985 nm; $R^2$ = 0.7903. (b(ii)) PdG sensors' FSH detection antibody binding curve fit parameters: a = -0.02967 ± 0.00282 nm; b = 0.5413 ± 0.2461; c = 0.05779 ± 0.03727 ng/mL; d = 0.001432 ± 0.003604 nm; $R^2$ = 0.8915.

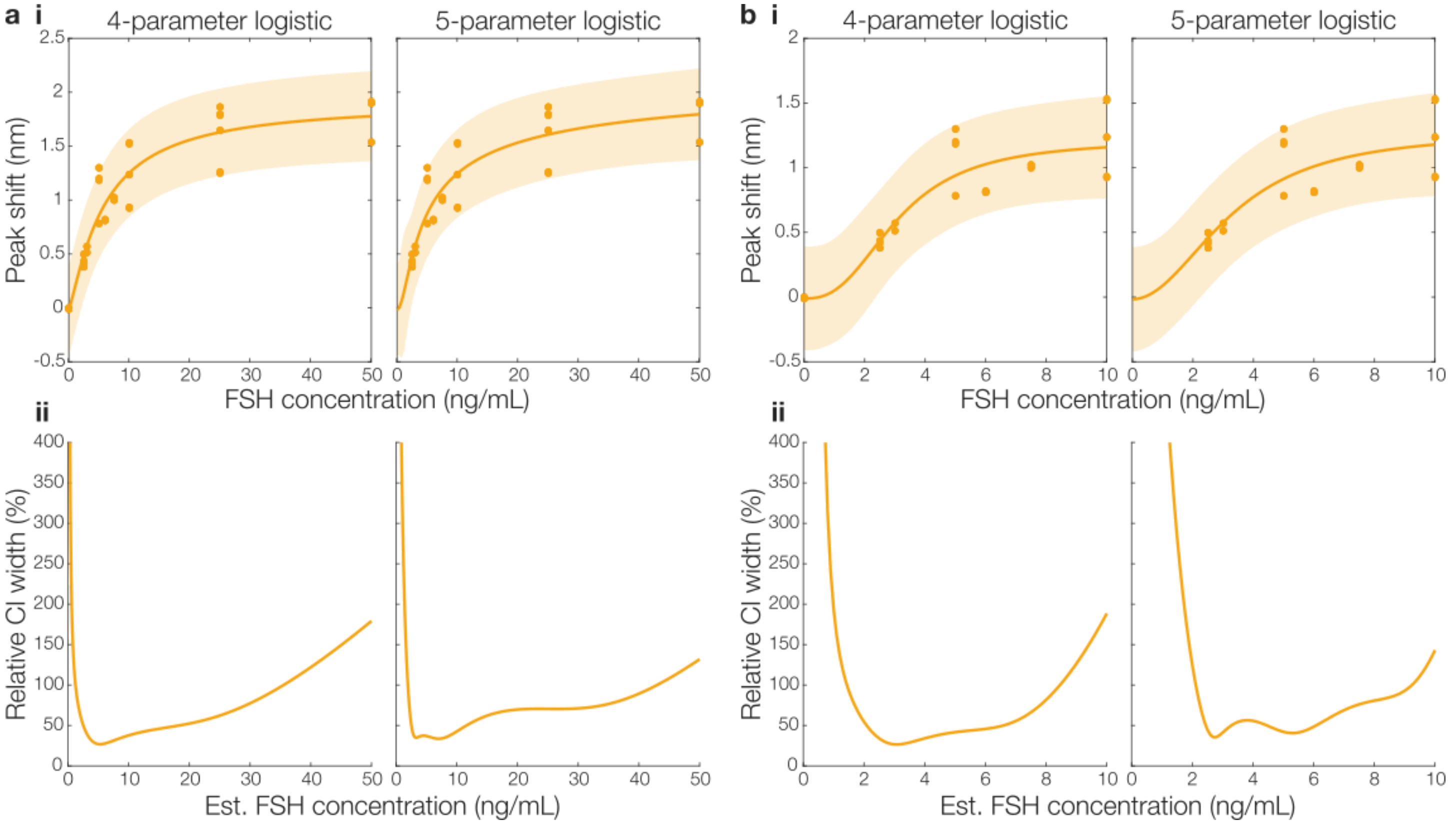

**Figure S4: Comparison of calibration fit equations and data ranges for FSH**. Two fit equations (4PL and 5PL) as well as two sets of calibration data were used to create and compare calibration curves. (a) The first set of calibration data used to fit the calibration model included peak shifts from a concentration range of 0-50 ng/mL. (b) Another set of calibration data was created using a shorter range of concentrations, 0-10 ng/mL to more accurately represent the quantitative range of the assay. (i) Calibration curve fits as well as (ii) relative CI widths are presented for both 4PL and 5PL models. Upper bound of the slope parameter b for both 5PL fits was constrained at 2 to prevent the emergence of imaginary numbers in predicted values. Lower bound of inflection point parameter c constrained at 0 for all fits.

Table S5. Fit parameters and performance characteristics of each fit on FSH sensor data.

| Conc. range | 0-50 ng/mL | | 0-10 ng/mL | |
|---|---|---|---|---|
| Fit | 4PL | 5PL | 4PL | 5PL |
| a (nm) | -0.01734 ± 0.16034 | -0.0144 ± 0.1592 | -0.01094 ± 0.15146 | -0.01652 ± 0.15198 |
| b | 1.214 ± 0.4727 | 2 | 2.594 ± 1.7879 | 2 |
| c (ng/m) | 5.82 ± 1.972 | 1.92 ± 1.4383 | 3.096 ± 0.799 | 3.811 ± 6.334 |
| d (nm) | 1.91 ± 0.287 | 2.272 ± 1.01 | 1.213 ± 0.2579 | 1.27 ± 0.5495 |
| g | – | 0.2405 ± 0.27448 | – | 1.281 ± 3.906 |
| $R^2$ | 0.8994 | 0.9009 | 0.8530 | 0.8515 |
| Min. relative CI width (%) | 26.8366 | 33.6060 | 26.5022 | 35.3956 |

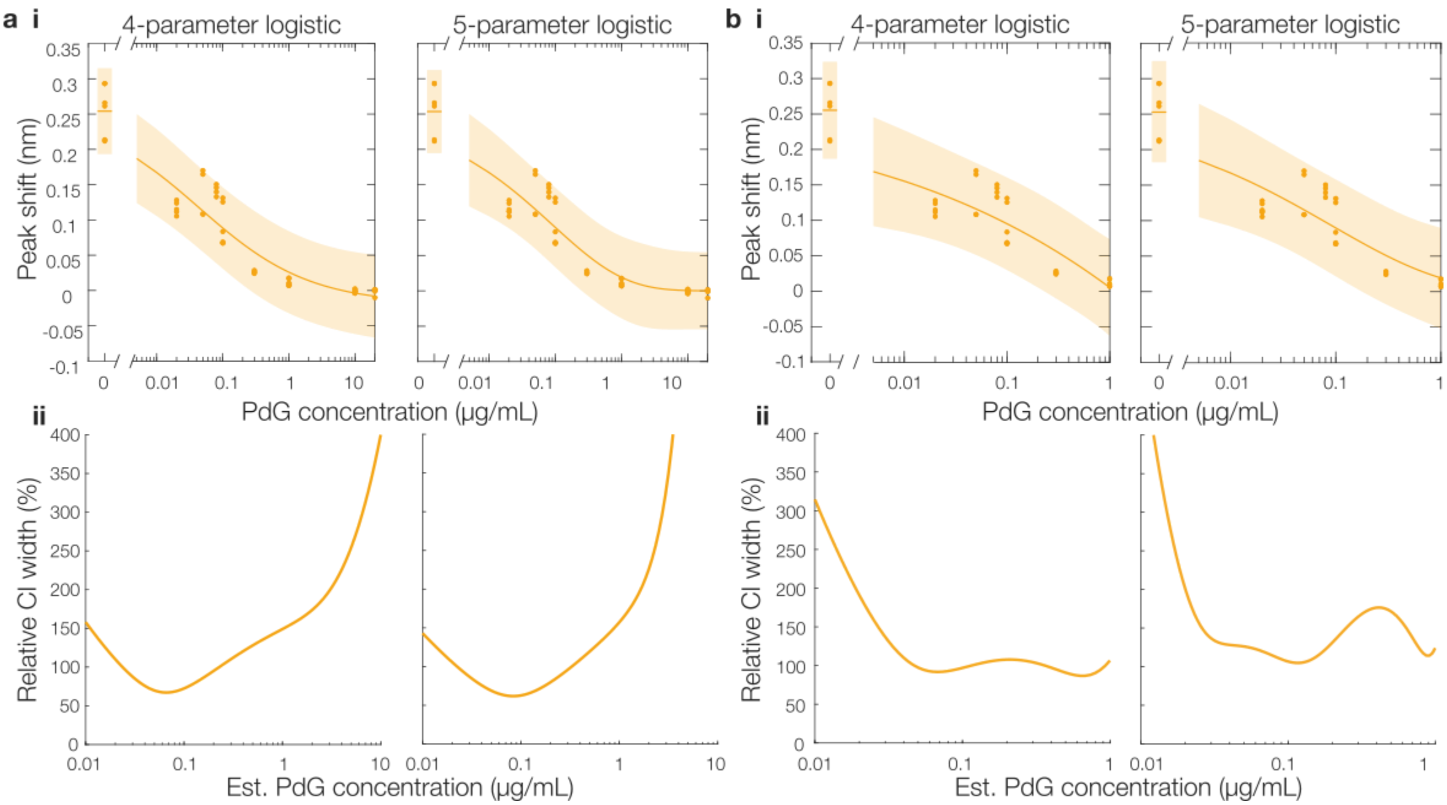

**Figure S5: Comparison of calibration fit equations and data ranges for PdG**. Two fit equations (4PL and 5PL) as well as two sets of calibration data were used to create and compare calibration curves. (a) The first set of calibration data used to fit the calibration model included peak shifts from a concentration range of 0-20 µg/mL. (b) Another set of calibration data was created using a shorter range of concentrations, 0-1 µg/mL to more accurately represent the quantitative range of the assay. (i) Calibration curve fits as well as (ii) relative CI widths are presented for both 4PL and 5PL models. Lower bound of inflection point parameter c constrained at 0 for all fits.

Table S6. Fit parameters and performance characteristics of each fit on PdG sensor data.

| Conc. range | 0-20 µg/mL | | 0-1 µg/mL | |
|---|---|---|---|---|
| Fit | 4PL | 5PL | 4PL | 5PL |
| a (nm) | 0.2542 ± 0.023 | 0.2535 ± 0.0224 | 0.2553 ± 0.0259 | 0.2555 ± 0.0265 |
| b | 0.5177 ± 0.2357 | 0.3948 ± 0.45613 | 0.2226 ± 0.4887 | 0.2803 ± 5.4343 |
| c (µg/mL) | 0.04346 ± 0.02647 | 866.6 ± 172566.6 | 1696 ± 172496 | 0.176 ± 160.576 |
| d (nm) | -0.01909 ± 0.03004 | -0.004098 ± 0.02322 | -1.304 ± 24.976 | -1.795 ± 1525.205 |
| g | – | 36.64 ± 2697.64 | – | 0.1332 ± 128.3332 |
| $R^2$ | 0.9027 | 0.9097 | 0.8708 | 0.87698 |
| Min. relative CI width (%) | 68.9884 | 70.7767 | 87.6712 | 104.8103 |

#### 5. Amplification performance vs. FSH concentration

Building upon the single-concentration data presented in Figure 2 of the mainbody manuscript, we assessed the Qdot amplification performance as a function of FSH concentration. The results of this assessment are plotted in Figure S6 and tabulated in Table S7. Similar to the relationship reported previously for another particle-mediated amplification approach[9], we observe increasing amplification factor for decreasing FSH concentration, with amplification factors of >100 for FSH concentrations ≤ 3 ng/mL.

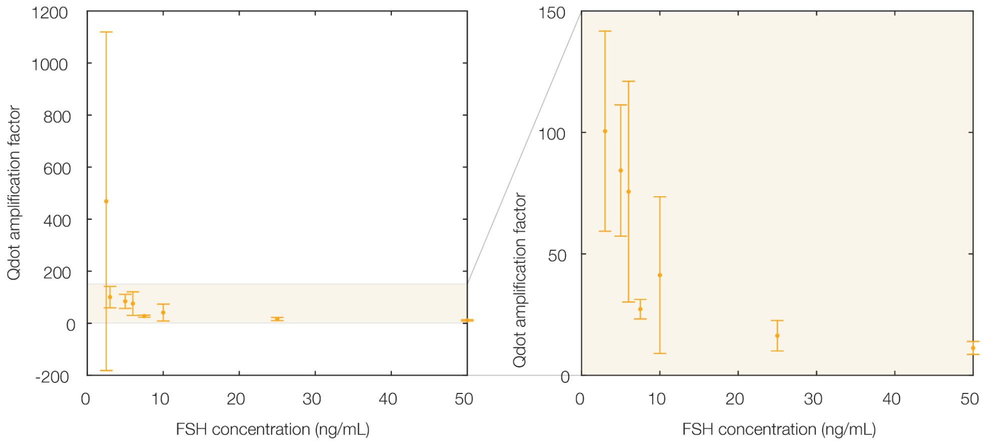

**Figure S6. Quantum dot-mediated amplification shows increasing amplification factor (Qdot signal normalized to FSH-binding signal) with decreasing FSH concentration.** Both the full range of amplification factors for FSH concentrations from 2.5-50 ng/mL (left) and a zoom-in showing only 3-50 ng/mL (right) are shown. amplification factors ranging from ~11 at 50 ng/mL FSH to >100 at 2.5-3 ng/mL FSH. Error bars represent one standard deviation of the amplification factors for n = 3-10 sensors. Error bars tend to be large due to the small primary FSH binding signal at low concentrations. Sensors that recorded unquantifiable primary FSH binding signal (e.g., small-magnitude negative values, potentially due to negative baseline drift resulting from unbinding of loosely attached functionalization or blocking reagents) were excluded from the analysis. Number of sensors included in the analysis and excluded due to negative FSH binding signal are reported in Table S7 below.

**Table S7. Quantum dot-mediated amplification shows increasing amplification factor (Qdot signal normalized to FSH-binding signal) with decreasing FSH concentration.** The mean, standard deviation, and number of sensors assessed to calculate these values are each reported. The number of sensors excluded from each analysis due to unquantifiable (negative) primary FSH binding signal are also reported.

| FSH concentration (ng/mL) | Average amplification factor | Standard deviation of amplification factor | Number of sensors assessed | Number of sensors excluded due to negative FSH signal |
|---|---|---|---|---|
| 2.5 | 469 | 651 | 3 | 3 |
| 3 | 101 | 41 | 6 | 2 |
| 5 | 84 | 27 | 3 | 3 |
| 6 | 76 | 45 | 4 | 0 |
| 7.5 | 27 | 4 | 4 | 0 |
| 10 | 41 | 32 | 6 | 0 |
| 25 | 16 | 6.3 | 10 | 0 |
| 50 | 11 | 2.6 | 8 | 0 |

#### 6. Comparison of multiplexed detection with and without the second analyte

In order to evaluate the impact that the presence of each analyte may have on the detection of the other analyte in the multiplexed assay, two assays were run with only a single analyte in the sample (FSH or PdG only) and Qdot-mediated amplification stage binding shifts were compared to those from assays with the second analyte present. The results of this comparison are presented in Figure S7. A two-sided Mann-Whitney U test was performed in MATLAB to compare the shifts and the results suggest no significant difference between the two conditions at the 5% significance level (p = 0.1333 for FSH and p = 0.8 for PdG), though this may be limited by the small sample size (n = 2 for single analyte in sample, n = 4 for both analytes in sample).

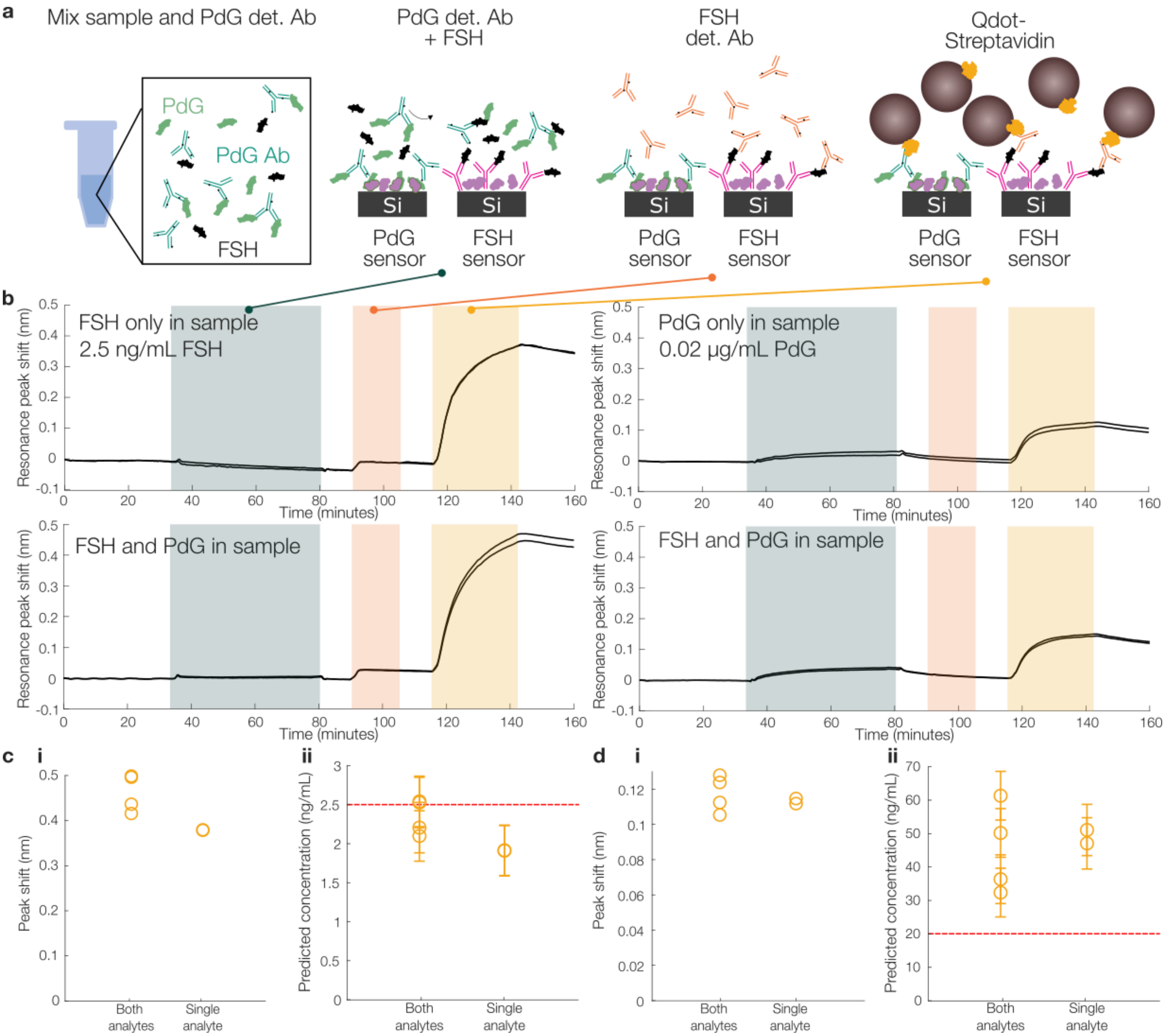

**Figure S7: Comparison of multiplexed detection with and without the second analyte**. (a) Overview of assay binding stages. (b) Reference-subtracted sensorgrams showing first binding cycle peak shifts for the primary sample binding, FSH detection antibody, and Qdot-mediated amplification stages for FSH (left) and PdG (right) sensors. (c-d) Beeswarm plots comparing peak shifts for the Qdot-mediated amplification stage when both analytes are present in sample vs. only one is present, for (c(i)) FSH and (d(i)) PdG sensors. Predicted concentrations from these peak shifts are plotted for both conditions (both analytes vs. single analyte in sample), for (c(ii)) FSH and (d(ii)) PdG sensors. Error bars represent the square root of the uncertainty of the prediction (variance $\nu^2$) provided by the model, calculated by propagation of error of all fitted parameters as described in section 9 of the Supplementary Information. Dashed line indicates the actual concentration. A Mann-Whitney U test revealed no significant difference between the two groups at the 5% significance level ($p = 0.1333$ for FSH and $p = 0.8$ for PdG).

### 7. Correction of quantification data by intrinsic sensitivity

As described in section 2 of the Supplementary Information, two of the trials used for our calibration analysis used resonators with a lower-duty cycle SWG waveguide, which resulted in a higher $S_{bulk}$. We sought to investigate whether correcting the resonance shift data by the $S_{bulk}$ for each sensor might improve the observed inter-assay replicability. To test this, we corrected each resonance shift by multiplying by that sensor's measured $S_{bulk}$ normalized to the average $S_{bulk}$ for all of the sensors used for the calibration analysis ($\Delta\lambda_{corr} = \Delta\lambda \cdot \frac{S_{bulk}}{\overline{S}_{bulk}}$). A comparison of the uncorrected and $S_{bulk}$-corrected quantum dot shifts for the FSH and PdG assays is presented in Figure SX.

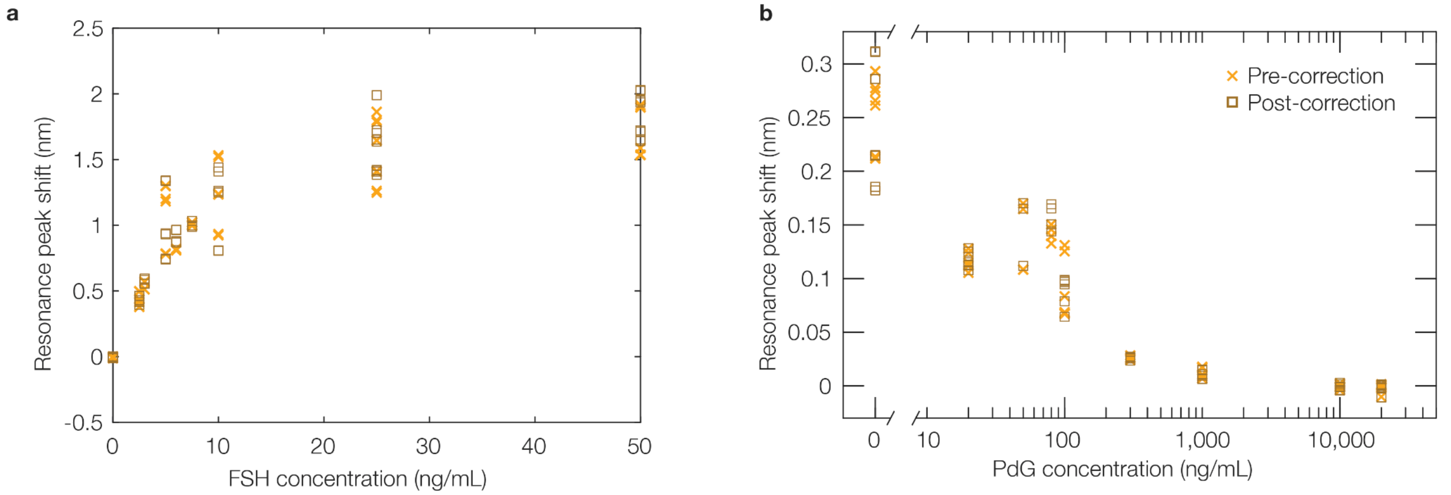

**Figure S8. Correcting the quantum dot amplification stage resonance peak shift signals in our multiplexed assay calibration and quantification data by the intrinsic sensitivity of each resonator ($S_{bulk}$).** (a-b) comparison of the resonance peak shift signals prior to correction (yellow 'x's) and after correction (brown squares) by the individual sensor's $S_{bulk}$, for the (a) FSH and (b) PdG sensor calibration data.

Overall, correction by the $S_{bulk}$ does not have a large effect on the results. Quantification of the mean, standard deviation, and CV of the Qdot signal for each measured concentration, both before and after $S_{bulk}$ correction, is presented in Table SX.Although the CVs of some concentrations are reduced by this correction, others slightly increase. These data suggest that inter-chip variability of the sensors' $S_{bulk}$ is not the dominant contributing factor to the variability observed in the immunoassay signal. Nevertheless, correcting each sensor's shift by its bulk refractive index sensitivity may be a strategy to improve variability in cases where the effects of fabrication variability are larger.

One limitation of this correction is that the sensors were exposed to 4 additional binding and regeneration cycles with different regeneration solutions after the initial binding cycle that was used to generate the calibration curve but prior to the bulk RI sensitivity measurement (data for these intermediate binding cycles are not reported here, as they are outside the scope of this manuscript). This exposure may have led to variability in the $S_{bulk}$ (for example, due to surface etching effects) that negatively impacted the efficacy of $S_{bulk}$ correction.

**Table S8. Impact of per-sensor signal correction based on the sensor's intrinsic $S_{bulk}$ on variability in observed sensor signal at each tested hormone concentration**. The $S_{bulk}$ correction does not appear to yield a significant improvement in the CV of the Qdot shifts ($p = 0.95$ for the FSH sensors; $p = 0.58$ for the PdG sensors).

| Analyte | Concentration (ng/mL) | Mean Qdot shift (nm) | Corrected mean Qdot shift (nm) | Standard deviation of Qdot shift (nm) | Standard deviation of corrected Qdot shift (nm) | Qdot shift CV (%) | Corrected Qdot shift CV (%) |
|---|---|---|---|---|---|---|---|
| FSH | 0 | -0.0069 | -0.0065 | 0.0049 | 0.0047 | 70.64 | 71.58 |
| | 2.5 | 0.4343 | 0.4322 | 0.0537 | 0.0265 | 12.37 | 6.13 |
| | 3 | 0.5411 | 0.5724 | 0.0330 | 0.0201 | 6.11 | 3.51 |
| | 5 | 1.0912 | 1.0056 | 0.2439 | 0.2724 | 22.35 | 27.09 |
| | 6 | 0.8133 | 0.9196 | 0.0060 | 0.0522 | 0.74 | 5.68 |
| | 7.5 | 1.0110 | 1.0023 | 0.0132 | 0.0210 | 1.30 | 2.09 |
| | 10 | 1.2301 | 1.1625 | 0.2673 | 0.2858 | 21.73 | 24.58 |
| | 25 | 1.5929 | 1.6325 | 0.2418 | 0.2292 | 15.18 | 14.04 |
| | 50 | 1.7321 | 1.8332 | 0.1832 | 0.1671 | 10.58 | 9.11 |
| | | | | | **Average:** | **11.3** | **11.5** |
| | | | | | **Standard deviation:** | **8.3** | **9.6** |
| PdG | 0 | 0.2614 | 0.2490 | 0.0321 | 0.0553 | 12.29 | 22.21 |
| | 20 | 0.1160 | 0.1170 | 0.0083 | 0.0080 | 7.20 | 6.83 |

| Analyte | Concentration (ng/mL) | Mean Qdot shift (nm) | Corrected mean Qdot shift (nm) | Standard deviation of Qdot shift (nm) | Standard deviation of corrected Qdot shift (nm) | Qdot shift CV (%) | Corrected Qdot shift CV (%) |
|---|---|---|---|---|---|---|---|
| | 50 | 0.1379 | 0.1398 | 0.0341 | 0.0324 | 24.76 | 23.18 |
| | 80 | 0.1421 | 0.1573 | 0.0077 | 0.0120 | 5.39 | 7.63 |
| | 100 | 0.0906 | 0.0885 | 0.0299 | 0.0140 | 33.03 | 15.78 |
| | 300 | 0.0261 | 0.0257 | 0.0019 | 0.0016 | 7.11 | 6.05 |
| | 1000 | 0.0114 | 0.0105 | 0.0048 | 0.0038 | 42.52 | 36.27 |
| | 10000 | -0.0010 | -0.0010 | 0.0019 | 0.0020 | 188.40 | 204.97 |
| | 20000 | -0.0015 | -0.0016 | 0.0037 | 0.0039 | 254.43 | 245.19 |
| | | | | | **Average:** | **18.9** | **16.9** |
| | | | | | **Standard deviation:** | **14.7** | **11.2** |

The full pre- and post-correction data for these experiments is presented in Table S9.

**Table S9. Per-sensor data for sensor resonance peak shift signal before and after correction for variable $S_{bulk}$, for each of the hormone measurement trials used for the sensor calibration reported in Figure 6.**

| | FSH | | | | | | | | | PdG | | | | | | | | |
|---|---|---|---|---|---|---|---|---|---|---|---|---|---|---|---|---|---|---|
| Trial | FSH conc. (ng/mL) | Sensor | $S_{bulk}$ (nm/RIU) | Sample shift (nm) | Corr. sample shift (nm) | FSH det. Ab shift (nm) | Corr. FSH det. Ab shift (nm) | Qdot shift (nm) | Corr. Qdot shift (nm) | PdG conc. (µg/mL) | Sensor | $S_{bulk}$ (nm/RIU) | Sample shift (nm) | Corr. sample shift (nm) | FSH det. Ab shift (nm) | Corr. FSH det. Ab shift (nm) | Qdot shift (nm) | Corr. Qdot shift (nm) |
| 1 | 0 | 1 | 362.46 | -1.90E-05 | -2.14E-05 | 3.55E-03 | 4.00E-03 | 0.0015 | 0.0017 | 0 | 9 | 362.57 | 5.47E-02 | 5.99E-02 | -2.54E-02 | -2.78E-02 | 0.2613 | 0.2860 |
| | 0 | 3 | 345.34 | -3.25E-02 | -3.83E-02 | 1.13E-03 | 1.33E-03 | -0.0078 | -0.0092 | 0 | 11 | 368.71 | 5.42E-02 | 5.84E-02 | -2.88E-02 | -3.10E-02 | 0.2658 | 0.2861 |
| | 50 | 2 | 382.05 | 2.37E-01 | 2.53E-01 | 5.92E-01 | 6.33E-01 | 1.8958 | 2.0239 | 20 | 10 | 378.65 | -1.67E-02 | -1.75E-02 | -4.90E-03 | -5.14E-03 | -0.0102 | -0.0107 |
| | 50 | 4 | 384.61 | 2.27E-01 | 2.40E-01 | 5.60E-01 | 5.93E-01 | 1.9148 | 2.0306 | 20 | 12 | 382.79 | -1.24E-02 | -1.29E-02 | -2.59E-03 | -2.68E-03 | -0.0012 | -0.0013 |
| 2 | 25 | 1 | 381.80 | 2.71E-01 | 2.89E-01 | 6.27E-01 | 6.69E-01 | 1.8637 | 1.9909 | 10 | 9 | 349.08 | -4.12E-03 | -4.69E-03 | 1.32E-03 | 1.50E-03 | 0.0012 | 0.0014 |
| | 25 | 3 | 381.95 | 2.68E-01 | 2.86E-01 | 6.23E-01 | 6.65E-01 | 1.8627 | 1.9892 | 10 | 11 | 344.80 | -5.35E-03 | -6.16E-03 | 1.86E-03 | 2.14E-03 | 0.0024 | 0.0028 |
| | 5 | 2 | 395.77 | -1.56E-02 | -1.60E-02 | 1.47E-01 | 1.52E-01 | 1.2973 | 1.3369 | 0.1 | 10 | 274.00 | -3.15E-02 | -4.56E-02 | -1.67E-02 | -2.42E-02 | 0.0676 | 0.0979 |
| | 5 | 4 | 395.59 | -1.95E-02 | -2.01E-02 | 1.44E-01 | 1.49E-01 | 1.3015 | 1.3419 | 0.1 | 12 | 274.89 | -2.61E-02 | -3.77E-02 | -1.75E-02 | -2.52E-02 | 0.0670 | 0.0967 |
| 3 | 50 | 1 | 397.38 | 1.44E-01 | 1.47E-01 | 3.90E-01 | 4.00E-01 | 1.8978 | 1.9479 | 20 | 9 | 421.13 | -1.55E-03 | -1.46E-03 | -1.07E-03 | -1.00E-03 | 0.0012 | 0.0011 |
| | 50 | 3 | 400.78 | 1.42E-01 | 1.44E-01 | 3.85E-01 | 3.92E-01 | 1.9013 | 1.9349 | 20 | 11 | 414.40 | -3.22E-03 | -3.08E-03 | -1.14E-03 | -1.09E-03 | 0.0015 | 0.0015 |
| | 5 | 2 | 429.12 | -2.95E-03 | -2.80E-03 | 5.21E-02 | 4.95E-02 | 0.7853 | 0.7464 | 0.1 | 10 | 423.59 | 1.21E-02 | 1.13E-02 | -6.56E-03 | -6.14E-03 | 0.0687 | 0.0644 |
| | 5 | 4 | 430.17 | 7.66E-03 | 7.26E-03 | 5.44E-02 | 5.16E-02 | 0.7801 | 0.7397 | 0.1 | 12 | 420.40 | 1.49E-02 | 1.41E-02 | -6.75E-03 | -6.37E-03 | 0.0835 | 0.0788 |
| 4 | 25 | 2 | 472.60 | 9.82E-02 | 8.47E-02 | 1.54E-01 | 1.33E-01 | 1.6473 | 1.4217 | 10 | 10 | 440.94 | -7.09E-03 | -6.38E-03 | 1.02E-04 | 9.18E-05 | -0.0009 | -0.0008 |
| | 25 | 4 | 473.03 | 6.34E-02 | 5.47E-02 | 1.40E-01 | 1.21E-01 | 1.6434 | 1.4170 | 10 | 12 | 440.93 | -1.80E-02 | -1.62E-02 | -1.26E-04 | -1.13E-04 | -0.0014 | -0.0013 |
| | 10 | 1 | 467.60 | 8.86E-03 | 7.72E-03 | 5.52E-02 | 4.82E-02 | 0.9231 | 0.8052 | 1 | 9 | 461.47 | -1.66E-03 | -1.43E-03 | 3.35E-04 | 2.88E-04 | 0.0178 | 0.0153 |
| | 10 | 3 | 470.89 | 2.00E-02 | 1.73E-02 | 5.69E-02 | 4.93E-02 | 0.9335 | 0.8086 | 1 | 11 | 462.10 | -1.97E-03 | -1.69E-03 | -5.81E-04 | -4.99E-04 | 0.0170 | 0.0146 |
| 5 | 5 | 2 | 521.92 | 1.23E-02 | 9.58E-03 | 7.58E-02 | 5.92E-02 | 1.2009 | 0.9385 | 0.1 | 10 | 526.87 | 1.25E-02 | 9.44E-03 | -1.30E-02 | -9.77E-03 | 0.1311 | 0.0988 |
| | 5 | 4 | 518.31 | 2.22E-02 | 1.75E-02 | 7.71E-02 | 6.07E-02 | 1.1822 | 0.9303 | 0.1 | 12 | 526.16 | 1.14E-02 | 8.59E-03 | -1.37E-02 | -1.03E-02 | 0.1254 | 0.0946 |
| | 0 | 1 | 502.87 | -3.58E-02 | -2.90E-02 | -6.51E-03 | -5.28E-03 | -0.0093 | -0.0075 | 0 | 9 | 542.31 | 4.82E-02 | 3.52E-02 | -3.04E-02 | -2.23E-02 | 0.2931 | 0.2145 |
| | 0 | 3 | 504.73 | -2.92E-02 | -2.36E-02 | -5.11E-03 | -4.13E-03 | -0.0071 | -0.0057 | 0 | 11 | 541.00 | 4.24E-02 | 3.11E-02 | -3.14E-02 | -2.31E-02 | 0.2933 | 0.2152 |
| 6 | 50 | 1 | 391.82 | 1.13E-01 | 1.18E-01 | 2.85E-01 | 2.96E-01 | 1.5878 | 1.6528 | 20 | 9 | 366.04 | -1.87E-02 | -2.02E-02 | 1.68E-04 | 1.82E-04 | -0.0014 | -0.0015 |
| | 50 | 3 | 394.80 | 1.14E-01 | 1.18E-01 | 2.82E-01 | 2.91E-01 | 1.5877 | 1.6402 | 20 | 11 | 359.90 | -1.96E-02 | -2.16E-02 | 1.09E-03 | 1.20E-03 | -0.0019 | -0.0021 |
| | 25 | 2 | 350.51 | 6.03E-02 | 7.01E-02 | 1.57E-01 | 1.83E-01 | 1.4064 | 1.6366 | 10 | 10 | 366.60 | -2.16E-02 | -2.34E-02 | -4.21E-03 | -4.56E-03 | -0.0011 | -0.0012 |
| | 25 | 4 | 348.85 | 6.46E-02 | 7.55E-02 | 1.55E-01 | 1.81E-01 | 1.4129 | 1.6520 | 10 | 12 | 362.07 | -1.99E-02 | -2.18E-02 | -2.62E-03 | -2.87E-03 | -0.0006 | -0.0007 |
| 7 | 10 | 1 | 400.17 | 6.23E-02 | 6.35E-02 | 1.44E-01 | 1.47E-01 | 1.2385 | 1.2624 | 1 | 9 | 424.75 | -1.52E-02 | -1.42E-02 | -7.08E-03 | -6.62E-03 | 0.0069 | 0.0064 |
| | 10 | 3 | 402.54 | 4.88E-02 | 4.95E-02 | 1.41E-01 | 1.43E-01 | 1.2336 | 1.2499 | 1 | 11 | 424.05 | -8.04E-03 | -7.53E-03 | -3.46E-03 | -3.23E-03 | 0.0070 | 0.0066 |
| | 0 | 2 | 430.62 | -4.27E-02 | -4.04E-02 | -1.18E-02 | -1.12E-02 | -0.0129 | -0.0122 | 0 | 10 | 465.54 | 4.61E-02 | 3.93E-02 | -3.16E-02 | -2.70E-02 | 0.2136 | 0.1821 |
| | 0 | 4 | 433.54 | -4.69E-02 | -4.41E-02 | -1.08E-02 | -1.02E-02 | -0.0124 | -0.0117 | 0 | 12 | 453.32 | 3.58E-02 | 3.13E-02 | -3.25E-02 | -2.85E-02 | 0.2118 | 0.1854 |

| | FSH | | | | | | | | | PdG | | | | | | | | |
|---|---|---|---|---|---|---|---|---|---|---|---|---|---|---|---|---|---|---|
| Trial | FSH conc. (ng/mL) | Sensor | $S_{bulk}$ (nm/RIU) | Sample shift (nm) | Corr. sample shift (nm) | FSH det. Ab shift (nm) | Corr. FSH det. Ab shift (nm) | Qdot shift (nm) | Corr. Qdot shift (nm) | PdG conc. (µg/mL) | Sensor | $S_{bulk}$ (nm/RIU) | Sample shift (nm) | Corr. sample shift (nm) | FSH det. Ab shift (nm) | Corr. FSH det. Ab shift (nm) | Qdot shift (nm) | Corr. Qdot shift (nm) |
| 8 | 25 | 1 | 427.60 | 1.19E-01 | 1.14E-01 | 3.11E-01 | 2.96E-01 | 1.7844 | 1.7021 | 10 | 9 | 365.92 | -1.69E-02 | -1.83E-02 | -2.35E-03 | -2.55E-03 | -0.0011 | -0.0012 |
| | 25 | 3 | 425.45 | 1.22E-01 | 1.17E-01 | 3.12E-01 | 3.00E-01 | 1.7983 | 1.7240 | 10 | 11 | 361.39 | -1.81E-02 | -1.99E-02 | -2.32E-03 | -2.54E-03 | -0.0009 | -0.0010 |
| | 10 | 2 | 434.19 | 6.05E-02 | 5.68E-02 | 1.66E-01 | 1.56E-01 | 1.5326 | 1.4397 | 1 | 10 | 388.19 | -1.34E-02 | -1.37E-02 | -3.44E-03 | -3.52E-03 | 0.0106 | 0.0108 |
| | 10 | 4 | 439.63 | 5.85E-02 | 5.43E-02 | 1.57E-01 | 1.46E-01 | 1.5191 | 1.4093 | 1 | 12 | 382.53 | -1.22E-02 | -1.27E-02 | -3.66E-03 | -3.79E-03 | 0.0091 | 0.0094 |
| 9 | 50 | 1 | 362.99 | 1.60E-01 | 1.80E-01 | 3.48E-01 | 3.91E-01 | 1.5334 | 1.7230 | 20 | 9 | 358.09 | -1.65E-02 | -1.83E-02 | 1.52E-03 | 1.68E-03 | 0.0003 | 0.0003 |
| | 50 | 3 | 366.48 | 1.62E-01 | 1.81E-01 | 3.42E-01 | 3.81E-01 | 1.5384 | 1.7122 | 20 | 11 | 355.67 | -1.79E-02 | -2.00E-02 | 1.46E-03 | 1.63E-03 | 0.0000 | 0.0000 |
| | 25 | 2 | 367.76 | 8.04E-02 | 8.92E-02 | 1.78E-01 | 1.97E-01 | 1.2480 | 1.3841 | 10 | 10 | 379.11 | -1.48E-02 | -1.55E-02 | -3.90E-05 | -4.08E-05 | -0.0040 | -0.0042 |
| | 25 | 4 | 365.93 | 7.97E-02 | 8.88E-02 | 1.76E-01 | 1.96E-01 | 1.2622 | 1.4069 | 10 | 12 | 380.69 | -1.51E-02 | -1.57E-02 | 1.03E-03 | 1.07E-03 | -0.0036 | -0.0038 |
| 10 | 7.5 | 1 | 403.17 | 4.63E-02 | 4.68E-02 | 9.37E-02 | 9.48E-02 | 1.0210 | 1.0330 | 0.02 | 9 | 395.65 | 2.32E-02 | 2.33E-02 | -1.44E-02 | -1.45E-02 | 0.1280 | 0.1283 |
| | 7.5 | 3 | 406.98 | 3.77E-02 | 3.77E-02 | 8.99E-02 | 9.01E-02 | 0.9963 | 0.9985 | 0.02 | 11 | 392.98 | 2.21E-02 | 2.23E-02 | -1.38E-02 | -1.40E-02 | 0.1239 | 0.1251 |
| | 2.5 | 2 | 430.25 | -1.51E-04 | -1.43E-04 | 2.37E-02 | 2.25E-02 | 0.4158 | 0.3942 | 0.3 | 10 | 414.37 | -5.22E-03 | -5.00E-03 | -4.18E-03 | -4.01E-03 | 0.0284 | 0.0272 |
| | 2.5 | 4 | 427.63 | 3.59E-04 | 3.42E-04 | 2.51E-02 | 2.40E-02 | 0.4365 | 0.4163 | 0.3 | 12 | 412.18 | -4.76E-03 | -4.59E-03 | -3.91E-03 | -3.76E-03 | 0.0265 | 0.0255 |
| 11 | 2.5 | 1 | 440.05 | 1.37E-02 | 1.27E-02 | 3.40E-02 | 3.15E-02 | 0.4989 | 0.4624 | 0.3 | 9 | 404.19 | -4.58E-03 | -4.49E-03 | -3.99E-03 | -3.92E-03 | 0.0241 | 0.0236 |
| | 2.5 | 3 | 438.09 | 3.25E-03 | 3.03E-03 | 3.10E-02 | 2.89E-02 | 0.4962 | 0.4619 | 0.3 | 11 | 378.17 | -8.02E-03 | -8.41E-03 | -3.89E-03 | -4.09E-03 | 0.0253 | 0.0265 |
| | 7.5 | 2 | 413.67 | 3.19E-02 | 3.15E-02 | 1.09E-01 | 1.08E-01 | 1.0035 | 0.9894 | 0.02 | 10 | 396.96 | 2.71E-02 | 2.71E-02 | -1.36E-02 | -1.36E-02 | 0.1125 | 0.1124 |
| | 7.5 | 4 | 422.34 | 3.53E-02 | 3.41E-02 | 1.07E-01 | 1.04E-01 | 1.0232 | 0.9881 | 0.02 | 12 | 388.18 | 2.45E-02 | 2.51E-02 | -1.36E-02 | -1.39E-02 | 0.1054 | 0.1077 |
| 12 | 3 | 1 | 377.09 | -2.70E-02 | -2.92E-02 | 3.14E-02 | 3.39E-02 | 0.5129 | 0.5548 | 0.05 | 9 | 396.21 | 2.45E-02 | 2.45E-02 | -2.09E-02 | -2.10E-02 | 0.1700 | 0.1703 |
| | 3 | 3 | 375.66 | -2.62E-02 | -2.85E-02 | 3.06E-02 | 3.33E-02 | 0.5121 | 0.5561 | 0.05 | 11 | 395.60 | 2.32E-02 | 2.33E-02 | -2.02E-02 | -2.03E-02 | 0.1647 | 0.1652 |
| | 2.5 | 2 | 359.81 | -3.03E-02 | -3.44E-02 | 2.41E-02 | 2.73E-02 | 0.3785 | 0.4291 | 0 | 10 | 354.24 | 4.53E-02 | 5.08E-02 | -3.22E-02 | -3.60E-02 | 0.2776 | 0.3110 |
| | 2.5 | 4 | 360.78 | -2.96E-02 | -3.35E-02 | 1.94E-02 | 2.20E-02 | 0.3797 | 0.4293 | 0 | 12 | 349.36 | 4.53E-02 | 5.14E-02 | -3.34E-02 | -3.79E-02 | 0.2746 | 0.3119 |
| 13 | 0 | 1 | 393.58 | -2.36E-02 | -2.45E-02 | -4.53E-03 | -4.69E-03 | -0.0049 | -0.0051 | 0.02 | 9 | 395.33 | 2.40E-02 | 2.41E-02 | -1.60E-02 | -1.60E-02 | 0.1147 | 0.1152 |
| | 0 | 3 | 394.52 | -1.28E-02 | -1.32E-02 | -2.64E-03 | -2.73E-03 | -0.0024 | -0.0025 | 0.02 | 11 | 392.60 | 1.72E-02 | 1.74E-02 | -1.79E-02 | -1.81E-02 | 0.1118 | 0.1130 |
| | 3 | 2 | 392.06 | 9.28E-03 | 9.65E-03 | 3.99E-02 | 4.15E-02 | 0.5716 | 0.5947 | 0.05 | 10 | 385.20 | 1.71E-02 | 1.76E-02 | -1.83E-02 | -1.89E-02 | 0.1085 | 0.1118 |
| | 3 | 4 | 396.52 | 9.66E-03 | 9.93E-03 | 4.02E-02 | 4.13E-02 | 0.5678 | 0.5840 | 0.05 | 12 | 384.22 | 2.79E-02 | 2.88E-02 | -1.44E-02 | -1.49E-02 | 0.1082 | 0.1118 |
| 14 | 6 | 1 | 380.58 | 1.84E-02 | 1.98E-02 | 5.39E-02 | 5.78E-02 | 0.8212 | 0.8801 | 0.08 | 9 | 367.45 | 2.89E-02 | 3.12E-02 | -1.73E-02 | -1.87E-02 | 0.1395 | 0.1506 |
| | 6 | 3 | 382.45 | 2.24E-02 | 2.39E-02 | 5.37E-02 | 5.73E-02 | 0.8149 | 0.8691 | 0.08 | 11 | 365.98 | 3.06E-02 | 3.32E-02 | -1.55E-02 | -1.68E-02 | 0.1327 | 0.1439 |
| | 6 | 2 | 341.46 | 5.97E-03 | 7.13E-03 | 5.11E-02 | 6.10E-02 | 0.8088 | 0.9662 | 0.08 | 10 | 352.69 | 2.20E-02 | 2.47E-02 | -1.76E-02 | -1.98E-02 | 0.1504 | 0.1692 |
| | 6 | 4 | 342.32 | 9.40E-03 | 1.12E-02 | 5.24E-02 | 6.24E-02 | 0.8084 | 0.9632 | 0.08 | 12 | 349.41 | 1.65E-02 | 1.87E-02 | -1.73E-02 | -1.96E-02 | 0.1457 | 0.1654 |

### 8. Details for materials and methods

#### 8.1 Photonic integrated circuit and microfluidic channel designs

Figure S9 presents details of the photonic integrated circuit and microfluidic channel designs, highlighting the overall photonic chip design, locations of the microfluidic channels integrated with the photonic chip, and sensor and waveguide architecture. Figure S10 presents details of the photonic integrated circuit used for inkjet spotting functionalization, using the same type of SWG ring resonators but different on-chip locations and I/O routing.

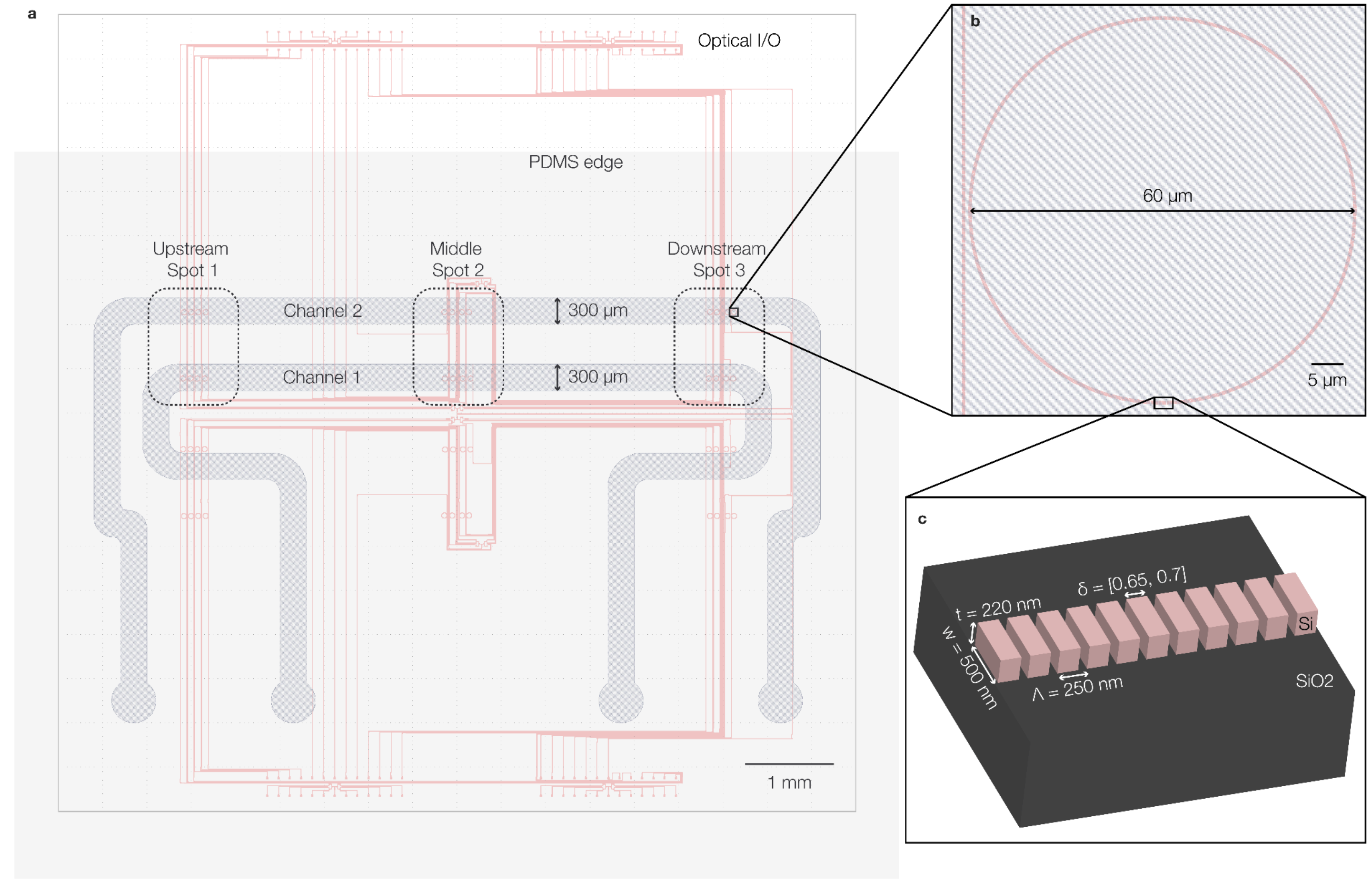

**Figure S9. Photonic integrated circuit and microfluidic channel details**. (a) Layout showing the silicon waveguides (pink), overlaid microfluidic channels (grey-blue), and photonic chip edge (grey outline), highlighting the 3 functionalization spot locations. The location of the top edge of the PDMS gasket is also shown. Each photonic chip contained 4 groups of 12 resonators; only one group was used on each chip. (b) Zoom-in showing one 60 µm diameter microring resonator (MRR) sensor. (c) Zoomed-in 3D rendering highlighting the sub-wavelength grating (SWG) waveguide architecture width $w$, thickness $t$, period $\Lambda$, and duty cycle $\delta$.

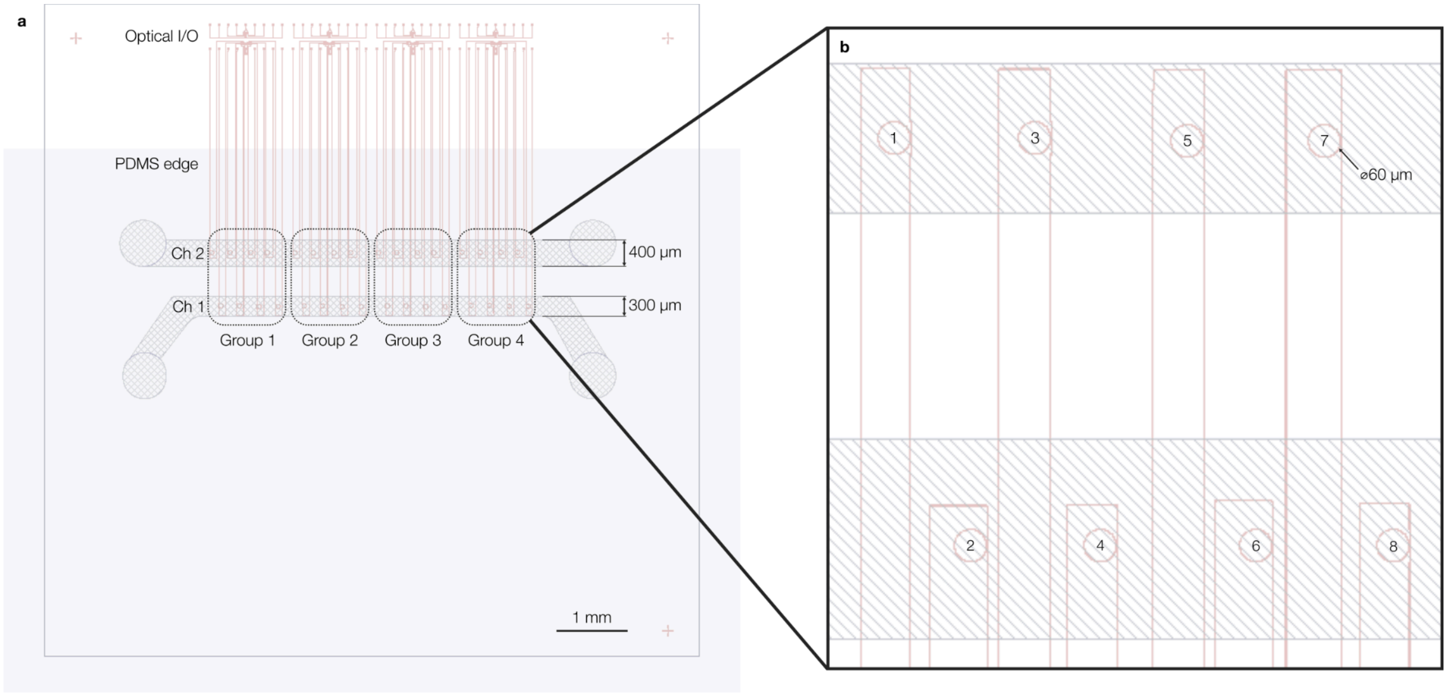

**Figure S10. Photonic integrated circuit and microfluidic channel details for sensor chips used in inkjet spotting experiments**. (a) Layout showing the silicon waveguides (pink), overlaid microfluidic channels (grey-blue), and photonic chip edge (grey outline). The location of the top edge of the PDMS gasket is also shown. Each photonic chip contained 4 groups of 8 resonators; only one group was used on each chip. (b) Zoom-in showing one 8-resonator group, with the resonators numbered from left-to-right.

#### 8.2 Assay solutions and fluidic control system reservoir setup

##### 8.2.1 Artificial urine preparation

For hormone marker detection in artificial urine, we used a previously published recipe for artificial urine[6]. This recipe is intended to reproduce key properties of human urine of healthy individuals, including concentrations of key compounds and spectral properties as measured by attenuated total reflection-Fourier transform infrared spectroscopy (ATR-FTIR). This previous work compared ATR-FTIR spectra of the artificial urine with those from first-morning, 8-hour fasted urine specimens from healthy volunteers, showing satisfactory agreement; however, some components were not included in the recipe (e.g., nucleic acids, amino acids and and proteins).

Artificial urine was prepared from ASTM Type I ultrapure water. The reagents used for artificial urine preparation as well as the concentrations of prepared stock solutions and the final concentration in artificial urine are provided in Table S10. For stock solution preparation instructions we also referenced a published protocol for artificial urine preparation[10].

**Table S10.** Constituents and reagent products used for artificial urine solution preparation. All stock solutions were prepared in ASTM Type I ultrapure water unless otherwise noted.

| Name | Vendor | Cat. number | Lot number | Shortened name | Concentration in artificial urine (mM) | Stock conc. (M) |
|---|---|---|---|---|---|---|
| Sodium sulfate | Sigma-Aldrich | 238597-500G | PCode: 1003706363 Source: MKCW0269 | Na2SO4 | 11.965 | 1.5 |
| Uric acid | Sigma-Aldrich | U2625-25G | PCode: 102771318 Source: BCCM0056 | UA | 1.487 | 0.025 (in 0.1 M NaOH) |
| Sodium Citrate Dihydrate | Fisher Chemical | S279-500 | 185194 | SCD | 2.45 | 0.175 |
| Creatinine | Sigma-Aldrich | C4255-10G | 384996 | Creatinine | 7.791 | 0.35 |
| Urea | Anachemia | 96238-300 | 310315 | Urea | 249.75 | 4 |
| Potassium chloride | EMD | PX1405-1 | XF23E | KCl | 30.953 | 2.25 |
| Sodium chloride | Fisher Scientific | S271-3 | 166221 | NaCl | 30.053 | 3 |
| Calcium chloride | Sigma-Aldrich | C5670-100G | SLCF4296 | CaCl2 | 1.663 | 1 |
| Ammonium chloride | Fisher Chemical | A661-500 | 241979 | NH4Cl | 23.667 | 2 |
| Potassium oxalate monohydrate | Sigma-Aldrich | 223425-500G | PCode: 1003687578, Source: MKCV4929 | POM | 0.19 | 0.1 |
| Magnesium sulfate heptahydrate | Sigma-Aldrich | 63138-250G | BCCC2138 | MSH | 4.389 | 0.5 |
| di-Sodium hydrogen phosphate dihydrate | Sigma-Aldrich | 1.06580-1000 | K46663780533 | di-SHPD | 4.667 | 0.25 |
| Sodium phosphate monobasic | Sigma-Aldrich | S0751-500G | SL8K1716V | SPM | 18.667 | 2 |

##### 8.2.2 FSH detection assays

**Table S11.** Fluidic control system reservoir setup and solution details for FSH detection assays for Qdot amplification demonstration (Figure 2).

| Reservoir | Assay solution | Priming solution | Solvent | Solution constituents, product and lot numbers |
|---|---|---|---|---|
| **1** | Ultrapure water RI | ⇇ | Ultrapure water | – |

| Reservoir | Assay solution | Priming solution | Solvent | Solution constituents, product and lot numbers |
|---|---|---|---|---|
| | standard | | | |
| **2** | 0.125 M NaCl RI standard | ⇇ | Ultrapure water | • 0.125 M NaCl (Thermo Fisher Scientific S271-3, lot 166221) |
| **3** | 0.250 M NaCl RI standard | ⇇ | Ultrapure water | • 0.250 M NaCl (Thermo Fisher Scientific S271-3, lot 166221) |
| **4** | Triton X-100 prewetting solution | ⇇ | PBS (Gibco 10010-023) | • 0.3 mM Triton X-100 (Sigma-Aldrich T8787-100ML, lot SLCJ6163) |
| **5** | Assay Running Buffer (ARB) | ⇇ | PBS | • 0.1 mg/mL BSA (Sigma-Aldrich A706-50G, lot 0000296702)<br>• 0.05% Tween 20 (Fisher Scientific BP337-500 lot 194435) |
| **6** | FSH detection Ab | ARB | ARB | • 2 µg/mL FSH detection antibody (BiosPacific A18056601P mouse IgG1, lot A8287, biotinylated in-house by the UBC Antibody and Biologics Core Facility) |
| **7** | Nonspecific challenge | ARB | ARB | • 1 mg/mL BSA (Sigma-Aldrich A706-50G, lot 0000296702) |
| **8** | FSH sample | ARB | ARB | • 25 ng/mL FSH (BiosPacific J18050015-LYO, lot TKA16) |
| **9a** | Qdot-streptavidin conjugate | ARB | ARB | • 0.01 µM Qdot-streptavidin conjugate (Thermo Scientific Q10163MP lot 2896575) |
| **9b** | Streptavidin-HRP | ARB | ARB | • 2 µg/mL streptavidin-HRP (Thermo Fisher Scientific Pierce High Sensitivity Streptavidin-HRP 21130, lot ZF394944) |
| **10** | pH 2.2 glycine-HCl regeneration solution | ⇇ | Ultrapure water | • 10 mM glycine (Sigma-Aldrich G8790-100G, lot SLCG2930)<br>• 160 mM NaCl (Thermo Fisher Scientific S271-3, lot 166221)<br>• Adjusted to pH 2.2 using 1 M HCl (Sigma-Aldrich 258148-2.5L-GL, PCode 4102988462, Source MKCS8144) |

⇇ These reservoirs were primed with the assay reagents.

**Table S12.** Fluidic control system reservoir setup and solution details for FSH detection assays (SI Section 1).

| Reservoir | Assay solution | Priming solution | Solvent | Solution constituents, product and lot numbers |
|---|---|---|---|---|
| **1** | Ultrapure water RI standard | ⇇ | Ultrapure water | – |
| **2** | 0.125 M NaCl RI standard | ⇇ | Ultrapure water | • 0.125 M NaCl (Thermo Fisher Scientific S271-3, lot 166221) |
| **3** | 0.250 M NaCl RI standard | ⇇ | Ultrapure water | • 0.250 M NaCl (Thermo Fisher Scientific S271-3, lot 166221) |
| **4** | Triton X-100 prewetting solution | ⇇ | PBS (Gibco 10010-023) | • 0.3 mM Triton X-100 (Sigma-Aldrich T8787-100ML, lot SLCJ6163) |
| **5** | Assay Running Buffer (ARB) | ⇇ | PBS | • 0.1 mg/mL BSA (Sigma-Aldrich A706-50G, lot 0000296702)<br>• 0.05% Tween 20 (Fisher Scientific BP337-500 lot 194435) |
| **6** | FSH detection Ab | ARB | ARB | • 2 µg/mL FSH detection antibody (BiosPacific A18056601P mouse monoclonal IgG1, lot A8287, biotinylated in-house by the UBC Antibody and Biologics Core Facility) |
| **7** | Nonspecific challenge | ARB | ARB | • 1 mg/mL BSA (Sigma-Aldrich A706-50G, lot 0000296702) |
| **8** | Sample (FSH or negative control) | ARB | ARB | • 50% artificial urine<br>• 0 ng/mL FSH (for negative control) or 25 ng/mL FSH (BiosPacific J18050015-LYO, lot TKA16) |
| **9** | Qdot-streptavidin conjugate | ARB | ARB | • 0.01 or 0.005 µM Qdot-streptavidin conjugate (Thermo Scientific Q10163MP lot 2896575) |
| **10** | pH 2.2 glycine-HCl regeneration solution | ⇇ | Ultrapure water | • 10 mM glycine (Sigma-Aldrich G8790-100G, lot SLCG2930)<br>• 160 mM NaCl (Thermo Fisher Scientific S271-3, lot 166221)<br>• Adjusted to pH 2.2 using 1 M HCl (Sigma-Aldrich |

| Reservoir | Assay solution | Priming solution | Solvent | Solution constituents, product and lot numbers |
|---|---|---|---|---|
| | | | | 258148-2.5L-GL, PCode 4102988462, Source MKCS8144) |

⇇ These reservoirs were primed with the assay reagents.

##### 8.2.3 PdG detection assays

**Table S13.** Fluidic control system reservoir setup and solution details for PdG detection assays (Figure 3).

| Reservoir | Assay solution | Priming solution | Solvent | Solution constituents, product and lot numbers |
|---|---|---|---|---|
| **1** | Ultrapure water RI standard | ⇇ | Ultrapure water | – |
| **2** | 0.125 M NaCl RI standard | ⇇ | Ultrapure water | • 0.125 M NaCl (Thermo Fisher Scientific S271-3, lot 166221) |
| **3** | 0.250 M NaCl RI standard | ⇇ | Ultrapure water | • 0.250 M NaCl (Thermo Fisher Scientific S271-3, lot 166221) |
| **4** | Triton X-100 prewetting solution | ⇇ | PBS (Gibco 10010-023) | • 0.3 mM Triton X-100 (Sigma-Aldrich T8787-100ML, lot SLCJ6163) |
| **5** | Assay Running Buffer (ARB) | ⇇ | PBS | • 0.1 mg/mL BSA (Sigma-Aldrich A706-50G, lot 0000296702)<br>• 0.05% Tween 20 (Fisher Scientific BP337-500, lot 194435) |
| **6** | Nonspecific challenge | ARB | ARB | • 1 mg/mL BSA (Sigma-Aldrich A706-50G, lot 0000296702) |
| **7** | Sample (PdG or negative control) | ARB | ARB | • 50% artificial urine<br>• 0 ng/mL PdG (for negative control) or 10,000 ng/mL PdG (BiosPacific J56131154, lot J0126)<br>• 1 µg/mL PdG detection antibody (BiosPacific mouse monoclonal IgG1 A56131314P, lot A8423, biotinylated in-house by the UBC Antibody and Biologics Core Facility) |
| **8** | Unused (prime with ARB) | ⇇ | ARB | – |
| **9** | Qdot-streptavidin conjugate | ARB | ARB | • 0.005 µM Qdot-streptavidin conjugate (Thermo Scientific Q10163MP lot 2896575) |
| **10** | pH 2.2 glycine-HCl regeneration solution | ⇇ | Ultrapure water | • 10 mM glycine (Sigma-Aldrich G8790-100G, lot SLCG2930)<br>• 160 mM NaCl (Thermo Fisher Scientific S271-3, lot 166221)<br>• Adjusted to pH 2.2 using 1 M HCl (Sigma-Aldrich 258148-2.5L-GL, PCode 4102988462, Source MKCS8144) |

⇇ These reservoirs were primed with the assay reagents.

##### 8.2.4 Pilot multiplexed detection assays

**Table S14.** Fluidic control system reservoir setup and solution details for pilot multiplexed detection assays (Figure 4).

| Reservoir | Assay solution | Priming solution | Solvent | Solution constituents, product and lot numbers |
|---|---|---|---|---|
| **1** | Ultrapure water RI standard | ⇇ | Ultrapure water | – |
| **2** | 0.125 M NaCl RI standard | ⇇ | Ultrapure water | • 0.125 M NaCl (Thermo Fisher Scientific S271-3, lot 166221) |
| **3** | 0.250 M NaCl RI standard | ⇇ | Ultrapure water | • 0.250 M NaCl (Thermo Fisher Scientific S271-3, lot 166221) |
| **4** | Triton X-100 prewetting solution | ⇇ | PBS (Gibco 10010-023) | • 0.3 mM Triton X-100 (Sigma-Aldrich T8787-100ML, lot SLCJ6163) |
| **5** | Assay Running Buffer (ARB) | ⇇ | PBS | • 0.1 mg/mL BSA (Sigma-Aldrich A706-50G, lot 0000296702)<br>• 0.05% Tween 20 (Fisher Scientific BP337-500, lot 194435) |
| **6** | Nonspecific challenge | ARB | ARB | • 1 mg/mL BSA (Sigma-Aldrich A706-50G, lot 0000296702) |

| | | | | |
|---|---|---|---|---|
| **7** | Sample (PdG/FSH or negative control) | ARB | ARB | • 50% artificial urine<br>• 0 ng/mL PdG and FSH (for negative control) or 10,000 ng/mL PdG (BiosPacific J56131154, lot J0126) and 25 ng/mL FSH (BiosPacific J18050015-LYO, lot TKA16)<br>• 1 µg/mL PdG detection antibody (BiosPacific mouse monoclonal IgG1 A56131314P, lot A8423, biotinylated in-house by the UBC Antibody and Biologics Core Facility) |
| **8** | FSH detection Ab | ARB | ARB | • 2 µg/mL FSH detection antibody (BiosPacific A18056601P mouse monoclonal IgG1, lot A8287, biotinylated in-house by the UBC Antibody and Biologics Core Facility) |
| **9** | Qdot-streptavidin conjugate | ARB | ARB | • 0.005 µM Qdot-streptavidin conjugate (Thermo Scientific Q10163MP lot 2896575) |
| **10** | pH 2.2 glycine-HCl regeneration solution | ⇇ | Ultrapure water | • 10 mM glycine (Sigma-Aldrich G8790-100G, lot SLCG2930)<br>• 160 mM NaCl (Thermo Fisher Scientific S271-3, lot 166221)<br>• Adjusted to pH 2.2 using 1 M HCl (Sigma-Aldrich 258148-2.5L-GL, PCode 4102988462, Source MKCS8144) |

⇇ These reservoirs were primed with the assay reagents.

##### 8.2.5 Multiplexed quantification assays

**Table S15.** Fluidic control system reservoir setup and solution details for multiplexed quantification assays (Figure 5).

| Reservoir | Assay solution | Priming solution | Solvent | Solution constituents, product and lot numbers |
|---|---|---|---|---|
| **1** | Ultrapure water RI standard | ⇇ | Ultrapure water | – |
| **2** | 0.125 M NaCl RI standard | ⇇ | Ultrapure water | • 0.125 M NaCl (Thermo Fisher Scientific S271-3, lot 166221) |
| **3** | 0.250 M NaCl RI standard | ⇇ | Ultrapure water | • 0.250 M NaCl (Thermo Fisher Scientific S271-3, lot 166221) |
| **4** | FSH detection Ab | ARB | ARB | • 2 µg/mL FSH detection antibody (BiosPacific A18056601P mouse monoclonal IgG1, lot A8287, biotinylated in-house by the UBC Antibody and Biologics Core Facility) |
| **5a (pre-wetting only)** | Triton X-100 prewetting solution | ⇇ | PBS (Gibco 10010-023) | • 0.3 mM Triton X-100 (Sigma-Aldrich T8787-100ML, lot SLCJ6163) |
| **5** | Assay Running Buffer (ARB) | ⇇ | PBS | • 0.1 mg/mL BSA (Sigma-Aldrich A706-50G, lot 0000296702)<br>• 0.05% Tween 20 (Fisher Scientific BP337-500, lot 194435) |
| **6** | Unused (prime with ARB) | ⇇ | ARB | – |
| **7** | Sample (PdG/FSH or negative control) | ARB | ARB | • 50% artificial urine<br>• 5 µg/mL PdG detection antibody (BiosPacific mouse monoclonal IgG1 A56131314P, lot A8423, biotinylated in-house by the UBC Antibody and Biologics Core Facility)<br>• PdG (BiosPacific J56131154, lot J0126) and FSH (BiosPacific J18050015-LYO, lot TKA16 or lot TKA17) at several sets of concentrations:<br>◦ 0 ng/mL FSH, 0 ng/mL PdG<br>◦ 2.5 ng/mL FSH, 300 ng/mL PdG<br>◦ 3 ng/mL FSH, 50 ng/mL PdG<br>◦ 5 ng/mL FSH, 100 ng/mL PdG<br>◦ 6 ng/mL FSH, 80 ng/mL PdG<br>◦ 7.5 ng/mL FSH, 20 ng/mL PdG<br>◦ 10 ng/mL FSH, 1,000 ng/mL PdG<br>◦ 25 ng/mL FSH, 10,000 ng/mL PdG<br>◦ 50 ng/mL FSH, 20,000 ng/mL PdG<br>◦ 2.5 ng/mL FSH, 0 ng/mL PdG |

| Reservoir | Assay solution | Priming solution | Solvent | Solution constituents, product and lot numbers |
|---|---|---|---|---|
| | | | | ○ 0 ng/mL FSH, 20 ng/mL PdG |
| **8** | Unused (prime with ARB) | ⇇ | ARB | – |
| **9** | Qdot-streptavidin conjugate | ARB | ARB | • 0.005 µM Qdot-streptavidin conjugate (Thermo Scientific Q10163MP lot 2896575) |
| **10** | pH 2.2 glycine-HCl regeneration solution | ⇇ | Ultrapure water | • 10 mM glycine (Sigma-Aldrich G8790-100G, lot SLCG2930)<br>• 160 mM NaCl (Thermo Fisher Scientific S271-3, lot 166221)<br>• Adjusted to pH 2.2 using 1 M HCl (Sigma-Aldrich 258148-2.5L-GL, PCode 4102988462, Source MKCS8144) |

⇇ These reservoirs were primed with the assay reagents.

##### 8.2.6 Multiplexed detection assays with inkjet functionalization

**Table S16.** Fluidic control system reservoir setup and solution details for multiplexed detection assays with inkjet functionalization (Figure 6).

| Reservoir | Assay solution | Priming solution | Solvent | Solution constituents, product and lot numbers |
|---|---|---|---|---|
| **1** | Ultrapure water RI standard | ⇇ | Ultrapure water | – |
| **2** | 0.125 M NaCl RI standard | ⇇ | Ultrapure water | • 0.125 M NaCl (Thermo Fisher Scientific S271-3, lot 166221) |
| **3** | 0.250 M NaCl RI standard | ⇇ | Ultrapure water | • 0.250 M NaCl (Thermo Fisher Scientific S271-3, lot 166221) |
| **4** | Triton X-100 prewetting solution | ⇇ | PBS (Gibco 10010-023) | • 0.3 mM Triton X-100 (Sigma-Aldrich T8787-100ML, lot SLCJ6163) |
| **5** | Unused (prime with ARB) | ⇇ | ARB | – |
| **6** | Assay Running Buffer (ARB) | ⇇ | PBS | • 0.1 mg/mL BSA (Sigma-Aldrich A706-50G, lot 0000296702)<br>• 0.05% Tween 20 (Fisher Scientific BP337-500, lot 194435) |
| **7** | Sample (PdG/FSH or negative control) | ARB | ARB | • 50% artificial urine<br>• 5 µg/mL PdG detection antibody (BiosPacific mouse monoclonal IgG1 A56131314P, lot A8423, biotinylated in-house by the UBC Antibody and Biologics Core Facility)<br>• 0.05 µg/mL PdG (BiosPacific J56131154, lot J0126)<br>• 25 ng/mL FSH (BiosPacific J18050015-LYO, lot TKA17) |
| **8** | FSH detection Ab | ARB | ARB | • 2 µg/mL FSH detection antibody (BiosPacific A18056601P mouse monoclonal IgG1, lot A8287, biotinylated in-house by the UBC Antibody and Biologics Core Facility) |
| **9** | Qdot-streptavidin conjugate | ARB | ARB | • 0.005 µM Qdot-streptavidin conjugate (Thermo Scientific Q10163MP lot 2896575) |
| **10** | Glycine-NaOH regeneration solution | ⇇ | Ultrapure water | • 10 mM glycine (Sigma-Aldrich G8790-100G, lot SLCG2930)<br>• 160 mM NaCl (Thermo Fisher Scientific S271-3, lot 166221)<br>• 10 mM NaOH (Fisher Chemical 1310-73-2, lot 205858)<br>• 0.05% Tween 20 (Fisher Scientific BP337-500, lot 194435) |

⇇ These reservoirs were primed with the assay reagents.

#### 8.3 Assay protocols

##### 8.3.1 FSH detection assays

**Table S17.** Automated fluidic protocol for FSH detection assays for Qdot amplification demonstration (Figure 2). Solution details are described in Section 8.2.2 of the Supplementary Information.

| Segment | Step | | Reservoir | Flow rate (µL/min) | Flow time (min) |
|---|---|---|---|---|---|
| Pre-wetting (non-automated) | Pre-1 | Triton X-100 pre-wetting sol'n | 4 | 1 | ~5 |
| | Pre-2 | PBS-BSA flow during setup & alignment | 5 | 30 | 60 |
| 0 (initial stability) | Initial stability | ARB stability | 5 | 30 | 30 |
| | 1.3 | Nonspecific challenge | 7 | 30 | 20 |
| | 1.4 | ARB rinse | 5 | 30 | 10 |
| | 1.5 | FSH | 8 | 30 | 25 |
| | 1.6 | ARB rinse | 5 | 30 | 10 |
| | 1.7 | Detection Ab (biotinylated α-FSH) | 6 | 30 | 30 |
| | 1.8 | ARB rinse | 5 | 30 | 10 |
| | 1.9 | Streptavidin-HRP or Qdot-streptavidin conjugate | 9 | 30 | 7 |
| | | Streptavidin-HRP or Qdot-streptavidin conjugate | 9 | 10 | 20 |
| 1 (detection round 1) | 1.10 | ARB stability | 5 | 30 | 30 |
| | 2.1 | Regeneration buffer | 10 | 30 | 5 |
| | 2.2 | ARB stability | 5 | 30 | 30 |
| | 2.3 | Nonspecific challenge | 7 | 30 | 20 |
| | 2.4 | ARB rinse | 5 | 30 | 10 |
| | 2.5 | FSH | 8 | 30 | 25 |
| | 2.6 | ARB rinse | 5 | 30 | 10 |
| | 2.7 | Detection Ab (biotinylated α-FSH) | 6 | 30 | 30 |
| | 2.8 | ARB rinse | 5 | 30 | 10 |
| | 2.9 | Streptavidin-HRP or Qdot-streptavidin conjugate | 9 | 30 | 5 |
| | | Streptavidin-HRP or Qdot-streptavidin conjugate | 9 | 10 | 20 |
| 2 (detection round 2) | 2.10 | ARB stability** Refill in last 10 mins | 5 | 30 | 40 |
| | 3.1 | Regeneration buffer | 10 | 30 | 5 |
| | 3.2 | ARB stability | 5 | 30 | 30 |
| | 3.3 | Nonspecific challenge | 7 | 30 | 20 |
| | 3.4 | ARB rinse | 5 | 30 | 10 |
| | 3.5 | FSH | 8 | 30 | 25 |
| | 3.6 | ARB rinse | 5 | 30 | 10 |
| | 3.7 | Detection Ab (biotinylated α-FSH) | 6 | 30 | 30 |
| | 3.8 | ARB rinse | 5 | 30 | 10 |
| | 3.9 | Streptavidin-HRP or Qdot-streptavidin conjugate | 9 | 30 | 5 |
| | | Streptavidin-HRP or Qdot-streptavidin conjugate | 9 | 10 | 20 |
| 3 (detection round 3) | 3.10 | ARB stability | 5 | 30 | 30 |
| | 4.1 | Regeneration buffer | 10 | 30 | 5 |
| Final regeneration | 4.2 | ARB stability | 5 | 30 | 30 |
| | 5.1 | ddW (extra to rinse out line) | 1 | 30 | 20 |
| | 5.2 | 0.125 M NaCl | 2 | 30 | 20 |

4 (bulk RI)

| Segment | Step | | Reservoir | Flow rate (µL/min) | Flow time (min) |
|---|---|---|---|---|---|
| | 5.3 | 0.250 M NaCl | 3 | 30 | 20 |
| | 5.4 | 0.125 M NaCl | 2 | 30 | 20 |
| | 5.5 | ddW | 1 | 30 | 20 |
| | 5.6 | 0.125 M NaCl | 2 | 30 | 20 |
| | 5.7 | 0.250 M NaCl | 3 | 30 | 20 |
| | 5.8 | 0.125 M NaCl | 2 | 30 | 20 |
| | 5.9 | ddW | 1 | 30 | 20 |
| | 5.10 | ddW overnight | 1 | 10 | 900 |

**Table S18.** Automated fluidic protocol for FSH detection assays (SI Section 1). Solution details are described in Section 8.2.2 of the Supplementary Information.

| Segment | Step | | Reservoir | Flow rate (µL/min) | Flow time (min) |
|---|---|---|---|---|---|
| Pre-wetting (non-automated) | Pre-1 | Triton X-100 pre-wetting sol'n | 4 | 1 | ~5 |
| | Pre-2 | ARB flow during setup & alignment | 5 | 30 | 60 |
| 0 (initial stability) | Initial stability | ARB stability | 5 | 30 | 30 |
| | 1.3 | Nonspecific challenge | 7 | 30 | 20 |
| | 1.4 | ARB rinse | 5 | 30 | 10 |
| | 1.5 | FSH or negative control | 8 | 30 | 25 |
| | 1.6 | ARB rinse | 5 | 30 | 10 |
| | 1.7 | Detection Ab (biotinylated α-FSH) | 6 | 30 | 20 or 30 |
| | 1.8 | ARB rinse | 5 | 30 | 10 |
| | 1.9 | Qdot-streptavidin conjugate | 9 | 30 | 7 |
| | | Qdot-streptavidin conjugate | 9 | 10 | 20 |
| 1 (detection round 1) | 1.10 | ARB stability | 5 | 30 | 30 |
| | 2.1 | Regeneration buffer | 10 | 30 | 5 |
| | 2.2 | ARB stability | 5 | 30 | 30 |
| | 2.3 | Nonspecific challenge | 7 | 30 | 20 |
| | 2.4 | ARB rinse | 5 | 30 | 10 |
| | 2.5 | FSH or negative control | 8 | 30 | 25 |
| | 2.6 | ARB rinse | 5 | 30 | 10 |
| | 2.7 | Detection Ab (biotinylated α-FSH) | 6 | 30 | 20 or 30 |
| | 2.8 | ARB rinse | 5 | 30 | 10 |
| | 2.9 | Qdot-streptavidin conjugate | 9 | 30 | 5 |
| | | Qdot-streptavidin conjugate | 9 | 10 | 20 |
| 2 (detection round 2) | 2.10 | ARB stability** Refill in last 10 mins | 5 | 30 | 40 |
| | 3.1 | Regeneration buffer | 10 | 30 | 5 |
| | 3.2 | ARB stability | 5 | 30 | 30 |
| | 3.3 | Nonspecific challenge | 7 | 30 | 20 |
| | 3.4 | ARB rinse | 5 | 30 | 10 |
| | 3.5 | FSH or negative control | 8 | 30 | 25 |
| | 3.6 | ARB rinse | 5 | 30 | 10 |

3 (detection round 3)

| | | | | | |
|---|---|---|---|---|---|
| | 3.7 | Detection Ab (biotinylated α-FSH) | 6 | 30 | 20 or 30 |
| | 3.8 | ARB rinse | 5 | 30 | 10 |
| | 3.9 | Qdot-streptavidin conjugate | 9 | 30 | 5 |
| | | Qdot-streptavidin conjugate | 9 | 10 | 20 |
| | 3.10 | ARB stability | 5 | 30 | 30 |
| Final regeneration | 4.1 | Regeneration buffer | 10 | 30 | 5 |
| | 4.2 | ARB stability | 5 | 30 | 30 |
| 4 (bulk RI) | 5.1 | ddW (extra to rinse out line) | 1 | 30 | 20 |
| | 5.2 | 0.125 M NaCl | 2 | 30 | 20 |
| | 5.3 | 0.250 M NaCl | 3 | 30 | 20 |
| | 5.4 | 0.125 M NaCl | 2 | 30 | 20 |
| | 5.5 | ddW | 1 | 30 | 20 |
| | 5.6 | 0.125 M NaCl | 2 | 30 | 20 |
| | 5.7 | 0.250 M NaCl | 3 | 30 | 20 |
| | 5.8 | 0.125 M NaCl | 2 | 30 | 20 |
| | 5.9 | ddW | 1 | 30 | 20 |
| | 5.10 | ddW overnight | 1 | 10 | 900 |

##### 8.3.2 PdG detection assays

**Table S19.** Automated fluidic protocol for PdG detection assays (Figure 3). Solution details are described in Section 8.2.3 of the Supplementary Information.

| Segment | Step | Reagent | Reservoir | Flow rate (µL/min) | Flow time (min) |
|---|---|---|---|---|---|
| Pre-wetting (non-automated) | Pre-1 | Triton X-100 pre-wetting sol'n | 4 | 2 | 30 |
| | Pre-2 | ARB rinse | 5 | 30 | 60 |
| 1 (detection round 1) | 1.01 | ARB stability | 5 | 30 | 30 |
| | 1.02 | Nonspecific challenge | 6 | 30 | 20 |
| | 1.03 | ARB rinse | 5 | 30 | 10 |
| | 1.04 | PdG sample + detection antibody | 7 | 30 | 7 |
| | 1.05 | PdG sample + detection antibody | 7 | 5 | 40 |
| | 1.06 | ARB rinse | 5 | 30 | 10 |
| | 1.07 | Qdot-streptavidin conjugate | 9 | 30 | 7 |
| | 1.08 | Qdot-streptavidin conjugate | 9 | 5 | 20 |
| | 1.09 | ARB stability | 5 | 30 | 30 |
| 2 (detection round 2) | 2.01 | Regeneration solution | 10 | 30 | 5 |
| | 2.02 | ARB stability | 5 | 30 | 30 |
| | 2.03 | Nonspecific challenge | 6 | 30 | 20 |
| | 2.04 | ARB rinse | 5 | 30 | 10 |

| Segment | Step | Reagent | Reservoir | Flow rate (µL/min) | Flow time (min) |
|---|---|---|---|---|---|
| | 2.05 | PdG sample + detection antibody | 7 | 30 | 5 |
| | 2.06 | PdG sample + detection antibody | 7 | 5 | 40 |
| | 2.07 | ARB rinse | 5 | 30 | 10 |
| | 2.08 | Qdot-streptavidin conjugate | 9 | 30 | 5 |
| | 2.09 | Qdot-streptavidin conjugate | 9 | 5 | 20 |
| | 2.10 | ARB stability | 5 | 30 | 30 |
| | 2.11 | ARB rinse during refill | 5 | 30 | 10 |
| 3 (detection round 3) | 3.01 | Regeneration solution | 10 | 30 | 5 |
| | 3.02 | ARB stability | 5 | 30 | 30 |
| | 3.03 | Nonspecific challenge | 6 | 30 | 20 |
| | 3.04 | ARB rinse | 5 | 30 | 10 |
| | 3.05 | PdG sample + detection antibody | 7 | 30 | 5 |
| | 3.06 | PdG sample + detection antibody | 7 | 5 | 40 |
| | 3.07 | ARB rinse | 5 | 30 | 10 |
| | 3.08 | Qdot-streptavidin conjugate | 9 | 30 | 5 |
| | 3.09 | Qdot-streptavidin conjugate | 9 | 5 | 20 |
| | 3.10 | ARB stability | 5 | 30 | 30 |
| 4 (final regen.) | 4.01 | Regeneration solution | 10 | 30 | 5 |
| | 4.02 | ARB stability | 5 | 30 | 30 |
| 5 (bulk RI) | 5.1 | ddW | 1 | 30 | 20 |
| | 5.2 | 0.125 M NaCl | 2 | 30 | 20 |
| | 5.3 | 0.250 M NaCl | 3 | 30 | 20 |
| | 5.4 | 0.125 M NaCl | 2 | 30 | 20 |
| | 5.5 | ddW | 1 | 30 | 20 |
| | 5.6 | 0.125 M NaCl | 2 | 30 | 20 |
| | 5.7 | 0.250 M NaCl | 3 | 30 | 20 |
| | 5.8 | 0.125 M NaCl | 2 | 30 | 20 |
| | 5.9 | ddW | 1 | 30 | 20 |
| | 5.10 | ddW | 1 | 10 | 900 |

##### 8.3.3 Pilot multiplexed detection assays

**Table S20.** Automated fluidic protocol for pilot multiplexed detection assays (Figure 4). Solution details are described in Section 8.2.4 of the Supplementary Information.

| Segment | Step | Reagent | Reservoir | Flow rate (μL/min) | Flow time (min) |
|---|---|---|---|---|---|
| Pre-wetting (non-automated) | Pre-1 | Triton X-100 pre-wetting sol'n | 4 | 2 | 30 |
| | Pre-2 | ARB rinse | 5 | 30 | 60 |
| 1 (detection round 1) | 1.01 | ARB stability | 5 | 30 | 30 |
| | 1.02 | Nonspecific challenge | 6 | 30 | 20 |
| | 1.03 | ARB rinse | 5 | 30 | 10 |
| | 1.04 | PdG/FSH sample + PdG detection antibody | 7 | 30 | 7 |
| | 1.05 | PdG/FSH sample + PdG detection antibody | 7 | 5 | 40 |
| | 1.06 | ARB rinse | 5 | 30 | 10 |
| | 1.07 | FSH detection antibody | 8 | 30 | 20 |
| | 1.08 | ARB rinse | 5 | 30 | 10 |
| | 1.09 | Qdot-streptavidin conjugate | 9 | 30 | 7 |
| | 1.1 | Qdot-streptavidin conjugate | 9 | 5 | 20 |
| | 1.11 | ARB stability | 5 | 30 | 30 |
| 2 (detection round 2) | 2.01 | Regeneration solution | 10 | 30 | 5 |
| | 2.02 | ARB stability | 5 | 30 | 30 |
| | 2.03 | Nonspecific challenge | 6 | 30 | 20 |
| | 2.04 | ARB rinse | 5 | 30 | 10 |
| | 2.05 | PdG/FSH sample + PdG detection antibody | 7 | 30 | 5 |
| | 2.06 | PdG/FSH sample + PdG detection antibody | 7 | 5 | 40 |
| | 2.07 | ARB rinse | 5 | 30 | 10 |
| | 2.08 | FSH detection antibody | 8 | 30 | 20 |
| | 2.09 | ARB rinse | 5 | 30 | 10 |
| | 2.1 | Qdot-streptavidin conjugate | 9 | 30 | 5 |
| | 2.11 | Qdot-streptavidin conjugate | 9 | 5 | 20 |
| | 2.12 | ARB stability | 5 | 30 | 30 |
| | 2.13 | ARB rinse during refill | 5 | 30 | 10 |
| 3 (detection round 3) | 3.01 | Regeneration solution | 10 | 30 | 5 |
| | 3.02 | ARB stability | 5 | 30 | 30 |
| | 3.03 | Nonspecific challenge | 6 | 30 | 20 |
| | 3.04 | ARB rinse | 5 | 30 | 10 |

| Segment | Step | Reagent | Reservoir | Flow rate (µL/min) | Flow time (min) |
|---|---|---|---|---|---|
| | 3.05 | PdG/FSH sample + PdG detection antibody | 7 | 30 | 5 |
| | 3.06 | PdG/FSH sample + PdG detection antibody | 7 | 5 | 40 |
| | 3.07 | ARB rinse | 5 | 30 | 10 |
| | 3.08 | FSH detection antibody | 8 | 30 | 20 |
| | 3.09 | ARB stability | 5 | 30 | 10 |
| | 3.1 | Qdot-streptavidin conjugate | 9 | 30 | 5 |
| | 3.11 | Qdot-streptavidin conjugate | 9 | 5 | 20 |
| | 3.12 | ARB stability | 5 | 30 | 30 |
| 4 (final regen.) | 4.01 | Regeneration solution | 10 | 30 | 5 |
| | 4.02 | ARB stability | 5 | 30 | 30 |
| 5 (bulk RI) | 5.1 | ddW | 1 | 30 | 20 |
| | 5.2 | 0.125 M NaCl | 2 | 30 | 20 |
| | 5.3 | 0.250 M NaCl | 3 | 30 | 20 |
| | 5.4 | 0.125 M NaCl | 2 | 30 | 20 |
| | 5.5 | ddW | 1 | 30 | 20 |
| | 5.6 | 0.125 M NaCl | 2 | 30 | 20 |
| | 5.7 | 0.250 M NaCl | 3 | 30 | 20 |
| | 5.8 | 0.125 M NaCl | 2 | 30 | 20 |
| | 5.9 | ddW | 1 | 30 | 20 |
| | 5.10 | ddW | 1 | 10 | 900 |

##### 8.3.4 Multiplexed quantification assays

**Table S21.** Automated fluidic protocol for multiplexed quantification assays (Figure 5). Solution details are described in Section 8.2.5 of the Supplementary Information.

| Segment | Step | Reagent | Reservoir | Flow rate (µL/min) | Flow time (min) |
|---|---|---|---|---|---|
| Pre-wetting (non-automated) | Pre-1 | Triton X-100 pre-wetting sol'n | 5a | 2 | 30 |
| | Pre-2 | ARB rinse | 5 | 30 | 60 |
| 1 (detection round 1) | 1.01 | ARB stability | 5 | 30 | 30 |
| | 1.02 | PdG/FSH sample + PdG detection antibody | 7 | 30 | 7 |
| | 1.03 | PdG/FSH sample + PdG detection antibody | 7 | 5 | 40 |
| | 1.04 | ARB rinse | 5 | 30 | 10 |
| | 1.05 | FSH detection antibody | 4 | 30 | 15 |
| | 1.06 | ARB rinse | 5 | 30 | 10 |
| | 1.07 | Qdot-streptavidin conjugate | 9 | 30 | 7 |
| | 1.08 | Qdot-streptavidin conjugate | 9 | 5 | 20 |

| Segment | Step | Reagent | Reservoir | Flow rate (µL/min) | Flow time (min) |
|---|---|---|---|---|---|
| | 1.09 | ARB stability | 5 | 30 | 30 |

##### 8.3.5 Multiplexed detection assays with inkjet functionalization

**Table S22.** Automated fluidic protocol for multiplexed detection assays with inkjet functionalization (Figure 6). Solution details are described in Section 8.2.6 of the Supplementary Information.

| Segment | Step | Reagent | Reservoir | Flow rate (µL/min) | Flow time (min) |
|---|---|---|---|---|---|
| Pre-wetting (non-automated) | Pre-1 | Triton X-100 pre-wetting sol'n | 4 | 2 | 30 |
| | Pre-2 | ARB rinse | 6 | 30 | 60 |
| 1 (detection round 1) | 1.01 | ARB stability | 6 | 30 | 30 |
| | 1.02 | Glycine-NaOH regeneration solution | 10 | 30 | 10 |
| | 1.03 | ARB stability | 6 | 30 | 30 |
| | 1.04 | PdG/FSH sample + PdG detection antibody | 7 | 30 | 7 |
| | 1.05 | PdG/FSH sample + PdG detection antibody | 7 | 5 | 40 |
| | 1.06 | ARB rinse | 6 | 30 | 10 |
| | 1.07 | FSH detection antibody | 8 | 30 | 15 |
| | 1.08 | ARB rinse | 6 | 30 | 10 |
| | 1.09 | Qdot-streptavidin conjugate | 9 | 30 | 7 |
| | 1.10 | Qdot-streptavidin conjugate | 9 | 5 | 20 |
| | 1.11 | ARB stability | 6 | 30 | 30 |

#### 8.4 Reference-subtraction example

For all of the data in manuscript Figures 1-6, the averaged resonance peak shift signal from reference sensors (functionalized with BSA) was subtracted from the signal from each sensor to yield the reference-subtracted sensor signal. Figure S11 illustrates the impact of this reference-subtraction process for a pilot multiplexed detection assay (a), comparing both (b) the raw sensor sensorgram signals and (c) the reference-subtracted sensorgram signals. By inspecting the sensorgrams during the nonspecific challenge portions of the binding cycles (purple overlay), it is clear that reference-subtraction reduces the impact of nonspecific binding: larger binding shifts are observable in the raw sensorgrams than in the reference-subtracted sensorgrams.

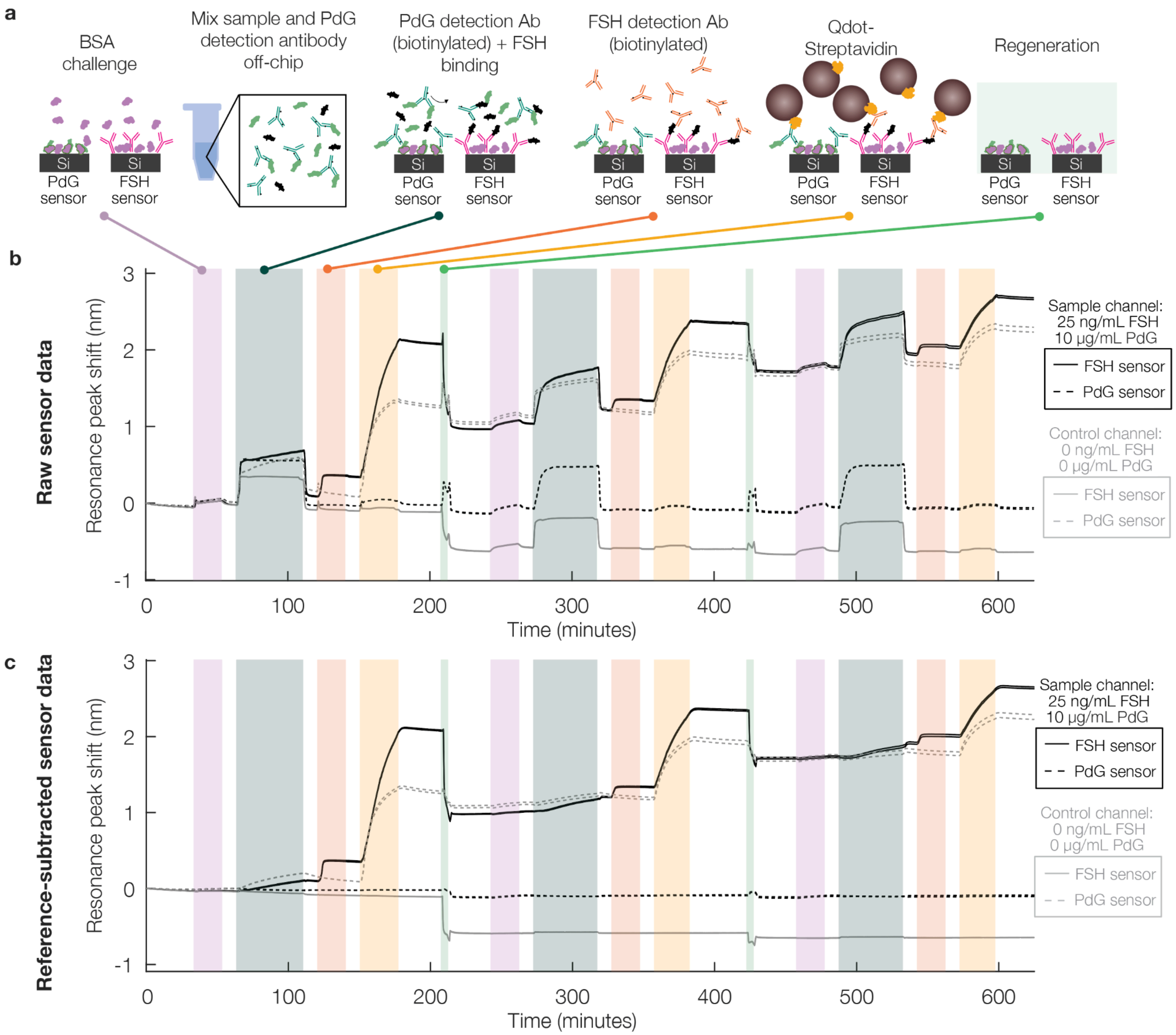

**Figure S11: Example of the impact of reference-subtraction in a 3-cycle multiplexed SiP hormone-detection assay**. (a) Schematic describing the assay stages in the multiplexed assay. (b) Raw sensorgram plots depicting the raw sensor resonance shift signal for each stage of the multiplexed detection assay. (c) Reference-subtracted sensorgram plots depicting the resonance shift signal for each sensor after subtraction of the reference sensor signal. Dashed lines denote PdG sensors while solid lines denote FSH sensors; black lines show the signal for sensors exposed to the hormone-containing sample while grey lines depict the signal for sensors exposed to the negative control sample.

The impact of reference-subtraction on reducing the impact of bulk refractive index changes is particularly apparent at the sample delivery assay stage of each binding cycle (blue-green overlay), where a notable change in bulk refractive index from the ARB is evident due to the presence of the artificial urine in the sample solution. This bulk refractive index shift is larger for the hormone-containing sample (black) than for the negative control (grey) because the PdG stock solution was prepared in ethanol, which has a higher refractive index than the aqueous buffer solutions. Reference-subtraction effectively removes this sharp shift up and down in the resonance peak position at the beginning and end of the sample delivery stage, permitting visualization of FSH and PdG detection antibody binding.

Overall, these results show that reference-subtraction using the data from BSA-functionalized reference sensors effectively reduces the effects of nonspecific binding and bulk refractive index changes from the sample fluid without introducing obvious artefacts in the data, for both FSH and PdG sensors.

#### 8.5 Piezoelectric inkjet printer

A custom-built piezoelectric inkjet dispense system was used for spotting-based functionalization of SiP sensor chips for multiplexed hormone detection. The substrate to be printed on (i.e., SiP chip) was positioned on a motorized XY linear stage (Zaber Technologies Inc. X-LSM050A-E03-KX13A and X-LSM100A-E03-PTB2). Piezoelectric inkjet nozzles (MicroFab Technologies Inc. MJ-AL-00-080) were mounted on a 3D-printed holder above the stage, which created ~5-mm Z-spacing between the nozzle orifice and substrate. An arbitrary waveform generator (Keysight 33500B) was used to define the printing actuation waveform, which was amplified 20× by a linear amplifier (PiezoDrive PD200), prior to reaching the nozzle. A top-view microscope camera (Aven Tools Mighty Scope) mounted above the stage facilitated system alignment and inspection of the printed droplets on the substrate. A high-speed side-view camera (Phantom VEO 710L), equipped with a 0.12–1.8× lens (PiShop 15-1) and placed opposite to a LED light source with a diffuser (Amazon B09M5KHBTC), enabled mid-air visualization of the jetted droplets and was used to monitor droplet jetting performance and satellite droplet formation during tuning of the actuation waveform. Custom PCBs and a microcontroller (Arduino Nano) were used to achieve communication between the system hardware and a custom control GUI (Python), operated on a PC. This GUI facilitated system alignment, actuation waveform tuning, and execution of automated printing runs in which droplets were printed with user-defined actuation waveforms at predefined locations on the substrate.

### 9. Hormone quantification calibration data analysis methods

For the hormone quantification assays, the binding shift data was obtained from only the first binding cycle of each assay, and reference subtraction and quantification was performed as described in Section 8.4 and in our previous work[3,8].

To quantify unknown concentrations of FSH and PdG, calibration data from both hormone assays were fitted with both a four-parameter (4PL) and five-parameter logistic (5PL) curve and compared. These models were selected as they offer strong performance for many immunoassays[11,12]. The inverse equation for each fit model was used for quantification, while the uncertainty on the quantified value was computed from the partial derivatives, variances, and covariances of each fit parameter as described previously for other fitted models [13]. The fit function equations, their parameters, and each of the partial derivatives are described in Table SX, with '$y$' representing the quantified sensor signal reading and '$x$' representing the analyte concentration. Partial derivatives for each parameter were computed in Symbolab. Two different datasets were used and compared during calibration to optimize the performance of the model: one containing the complete range of concentrations tested for each analyte (0-25 ng/mL of FSH, 0-10 µg/mL of PdG) and one with a shortened range of concentrations reflecting a more narrow analytic range for both analytes (0-10 ng/mL of FSH, 0-1 µg/mL of PdG).

Table S23. Details for the two sensor calibration fit models compared in section 4 of the Supplementary information, including the fit equation, inverse equation used for quantification, fit parameter details, and partial derivatives used in the uncertainty calculations. Degrees of freedom $df$ is the same as that used to estimate the residual standard deviation in the fit of the model [14]. $df = n - k$ where n is the number of fit points and k is the number of parameters being estimated [15].

| | **4-parameter logistic regression (4PL)** | **5-parameter logistic regression (5PL)** |
|---|---|---|
| **Equation** | $y = d + \frac{a-d}{1+(\frac{x}{c})^b}$ | $y = d + \frac{a-d}{(1+(\frac{x}{c})^b)^g}$ |
| **Inverse equation** | $x = c(\frac{a-y}{y-d})^{\frac{1}{b}}$ | $x = c((\frac{a-d}{y-d})^{\frac{1}{g}} - 1)^{\frac{1}{b}}$ |
| **Parameters** | a: the zero-dose asymptote (minimum (for FSH assay) or maximum (for PdG assay) peak shift that can be quantifiable)<br>b: the slope parameter (related to the steepness of the curve at point c)<br>c: inflection point, related to the EC50 | a: the zero-dose asymptote<br>b: the slope parameter (related to the steepness of the curve at point c)<br>c: inflection point, related to the EC50<br>d: infinite-dose asymptote<br>g: degree of asymmetry in the shape of the sigmoidal curve with respect to "EC50" |

| | | |
|---|---|---|
| | d: infinite-dose asymptote (maximum (for FSH assay) or minimum (for PdG assay) peak shift that can be quantifiable) | |
| **Partial derivatives** | $\frac{\partial}{\partial y} = \frac{c(-a+d)(a-y)^{(1-b)/b}}{b(y-d)^{(1+b)/b}}$ | $\frac{\partial}{\partial y} =- \frac{c((a-d)^{1/g}-(y-d)^{1/g})^{(1-b)/b}(a-d)^{1/g}}{bg(y-d)^{(bg+1)/bg}}$ |
| | $\frac{\partial}{\partial a} = \frac{c(a-y)^{(1-b)/b}}{b(y-d)^{1/b}}$ | $\frac{\partial}{\partial a} = \frac{c((a-d)^{1/g}-(y-d)^{1/g})^{(1-b)/b}(a-d)^{(-g+1)/g}}{bg(y-d)^{1/bg}}$ |
| | $\frac{\partial}{\partial b} =- \frac{cln(\frac{a-y}{y-d})(a-y)^{1/b}}{b^2(y-d)^{1/b}}$ | $\frac{\partial}{\partial b} =- \frac{cln((\frac{a-d}{y-d})^{1/g}-1)((\frac{a-d}{y-d})^{1/g}-1)^{1/b}}{b^2}$ |
| | $\frac{\partial}{\partial c} = (\frac{a-y}{y-d})^{\frac{1}{b}}$ | $\frac{\partial}{\partial c} = ((\frac{a-d}{y-d})^{1/g} - 1)^{1/b}$ |
| | $\frac{\partial}{\partial d} = \frac{c(a-y)^{1/b}}{b(y-d)^{(1+b)/b}}$ | $\frac{\partial}{\partial d} = \frac{c(a-y)((a-d)^{1/g}-(y-d)^{1/g})^{(1-b)/b}(-d+a)^{(-g+1)/g}}{gb(y-d)^{(gb+1)/gb}}$ |
| | | $\frac{\partial}{\partial g} - \frac{cln(\frac{a-d}{y-d})((a-d)^{1/g}-(y-d)^{1/g})^{(1-b)/b}(a-d)^{1/g}}{bg^2(y-d)^{1/bg}}$ |
| **Degrees of freedom** | $dof = n - 4$ | $dof = n - 5$ |

Calibration curve fitting was performed in MATLAB using custom scripts. The uncertainty of the prediction $v^2$ was calculated as the sum of the variance defined from propagation of error $u^2$ and the covariance terms between each parameter, as well as the standard deviation of the measurement $s_y$ [13].

$$u^2 = (\frac{\partial x}{\partial y})^2 s_y^{\ 2} + (\frac{\partial x}{\partial a})^2 s_a^{\ 2} + (\frac{\partial x}{\partial b})^2 s_b^{\ 2} + (\frac{\partial x}{\partial c})^2 s_c^{\ 2} + (\frac{\partial x}{\partial d})^2 s_d^{\ 2}$$

$$v^2 = u^2 + 2(\frac{\partial x}{\partial a})(\frac{\partial x}{\partial b})s_{ab} + 2(\frac{\partial x}{\partial a})(\frac{\partial x}{\partial c})s_{ac} + 2(\frac{\partial x}{\partial d})(\frac{\partial x}{\partial d})s_{ad} + 2(\frac{\partial x}{\partial b})(\frac{\partial x}{\partial c})s_{bc} + 2(\frac{\partial x}{\partial b})(\frac{\partial x}{\partial d})s_{bd} + 2(\frac{\partial x}{\partial c})(\frac{\partial x}{\partial d})s_{cd}$$

$$u^2 = (\frac{\partial x}{\partial y})^2 s_y^{\ 2} + (\frac{\partial x}{\partial a})^2 s_a^{\ 2} + (\frac{\partial x}{\partial b})^2 s_b^{\ 2} + (\frac{\partial x}{\partial c})^2 s_c^{\ 2} + (\frac{\partial x}{\partial d})^2 s_d^{\ 2} + (\frac{\partial x}{\partial g})^2 s_g^{\ 2}$$

$$v^2 = u^2 + 2(\frac{\partial x}{\partial a})(\frac{\partial x}{\partial b})s_{ab} + 2(\frac{\partial x}{\partial a})(\frac{\partial x}{\partial c})s_{ac} + 2(\frac{\partial x}{\partial d})(\frac{\partial x}{\partial d})s_{ad} + 2(\frac{\partial x}{\partial b})(\frac{\partial x}{\partial c})s_{bc} + 2(\frac{\partial x}{\partial b})(\frac{\partial x}{\partial d})s_{bd} + 2(\frac{\partial x}{\partial c})(\frac{\partial x}{\partial d})s_{cd}$$
$$+ 2(\frac{\partial x}{\partial g})(\frac{\partial x}{\partial a})s_{ag} + 2(\frac{\partial x}{\partial g})(\frac{\partial x}{\partial b})s_{bg} + 2(\frac{\partial x}{\partial g})(\frac{\partial x}{\partial c})s_{cg} + 2(\frac{\partial x}{\partial g})(\frac{\partial x}{\partial d})s_{dg}$$

Upper and lower bounds of a 95% confidence interval (CI) were calculated according to the formula:

$$CI = x_{found} \pm \sqrt{v^2} \cdot t_{1-\alpha/2,\upsilon}$$ [16]

where $x_{found}$ is the predicted concentration and $v^2$ is the uncertainty of the prediction. The student's t-statistic was indexed at the 5% significance level and the degrees of freedom noted in Table S23.

To find the relative widths of the 95% CIs, a range of concentrations to be predicted was defined. To obtain confidence intervals and errors, the concentrations were converted to peak shifts using the 4PL or 5PL model and then a MATLAB function was used to perform the confidence interval predictions. The standard deviation of the measurement ($s_y$) used to make these predictions was the average quantification error computed by propagating the standard deviations of the quantified peak positions from n = 10 peak position measurements before and after each binding shift using custom MATLAB scripts. The average quantification error is calculated from the standard deviation of the resonance peak position in the two regions that were subtracted to obtain the binding shift (i.e., $s_y = \sqrt{s_{y,pre}^2 + s_{y,post}^2}$, where $s_{y,pre}$ is the standard deviation of the quantified peak position prior to the binding shift step and $s_{y,post}$ is the standard deviation of the quantified peak position after the binding shift step). The relative widths of these 95% CIs (normalized to the estimated concentration) were calculated as:

$$CI_{rel} = 100 \cdot \frac{2\sqrt{v^2} \cdot t_{1-\alpha/2,\upsilon}}{estConc}$$

The fitted curve, 95% CIs, and relative CI widths were plotted in MATLAB.
